# Charting structural brain asymmetry across the human lifespan

**DOI:** 10.1101/2025.07.21.665924

**Authors:** Lena Dorfschmidt, Simon White, Margaret Gardner, Saashi Bedford, Gareth Ball, A. David Edwards, Yuanjun Gu, Yuankai He, Hao Huang, Shivaram Karandikar, Vanessa Kyriakopoulou, Minhui Ouyuang, Elmo P. Pulli, Armin Raznahan, Emma C. Robinson, Rafael Romero-García, Theodore D Satterthwaite, Jenna Schabdach, Isaac Sebenius, Zhiqiang Sha, Russell T. Shinohara, Jetro Tuulari, Matilde M Vaghi, Jacob Vogel, František Váša, Logan Williams, Dabriel Zimmerman, Lifespan Brain Chart Consortium, Varun Warrier, Konrad Wagstyl, Edward T Bullmore, Richard A.I. Bethlehem, Jakob Seidlitz, Aaron Alexander-Bloch

**Author notes:** A full list of consortium authors can be found in the appendix. Equal contributions (Jakob Seidlitz, Aaron Alexander-Bloch).

## Abstract

Lateralization is a fundamental principle of structural brain organization. In vivo imaging of brain asymmetry is essential for deciphering lateralized brain functions and their disruption in neurodevelopmental and neurodegenerative disorders. Here, we present a normative framework for benchmarking brain asymmetry across the lifespan, developed from an aggregated sample of 128 primary neuroimaging studies, including 177,701 scans from 138,231 individuals, jointly spanning the age range from 20 post menstrual weeks to 102 years. This resource includes comprehensive, hemisphere-specific brain growth charts for multiple neuroimaging phenotypes: regional cortical grey matter volume, thickness, surface area, and subcortical volumes. Our findings reveal distinct spatial patterns of asymmetry, with early leftward asymmetry observed in association cortices and late rightward asymmetry in sensory regions. These trajectories support theories of the neuroplasticity of asymmetry and the role of both genetic and environmental factors in shaping brain lateralization. Additionally, we provide tools to generate asymmetry centile scores, which allow the quantification of individual deviations from typical asymmetry throughout the lifespan and can be applied to unseen data or clinical populations. We demonstrate the utility of these models by highlighting group-level differences in asymmetry in autism spectrum disorder, schizophrenia, and Alzheimer’s disease, and exploring genetic correlations with hemispheric specialization. To facilitate further research, we have made this normative framework freely available as an interactive open-access resource (upon publication), offering an essential tool to advance both basic and clinical neuroscience.

## Introduction

Structural asymmetry is a fundamental principle of brain organization (Ocklenburg et al. 2024), supporting lateralized cognitive and motor functions and contributing to inter-individual differences in behavior and disease susceptibility (Postema, Hoogman, et al. 2021; Saltoun et al. 2024). However, substantial variability in the degree and pattern of brain asymmetry has been observed between individuals (Maingault et al. 2016). Normative brain asymmetry undergoes profound changes over the course of life and alterations in asymmetry have been observed in disease (Roe, Vidal-Pineiro, et al. 2023). Notably, altered brain asymmetry has been linked to multiple preclinical symptoms (Tomasi and Volkow 2025), as well as neuropsychiatric and neurodegenerative disorders, including autism spectrum disorder (ASD; (Postema, Van Rooij, et al. 2019)), schizophrenia (SCZ), and Alzheimer’s disease(AD; (Roe, Vidal-Piñeiro, et al. 2021)). Further, multiple genetic loci associated with different aspects of brain asymmetry have also been shown to overlap with genetic risk markers of neuropsychiatric disorders (Sha, Schijven, et al. 2021), highlighting the relevance of brain asymmetry as a potential marker of both typical and atypical neurodevelopment.

While certain asymmetries, such as left-hemispheric dominance for language, are considered evolutionarily conserved and adaptive (Corballis 2009), the developmental pathways that give rise to such specialization are not perfectly deterministic. The theory of fluctuating asymmetry — random deviations from bilateral symmetry — suggests that inherent variability or “developmental noise” affects asymmetric morphogenesis. As described by (Van Valen 1962), fluctuating asymmetry reflects an organism’s inability to strictly adhere to genetically prescribed developmental trajectories under varying environmental and internal conditions. In this context, functional asymmetries may emerge from a balance between genetically guided lateralization and stochastic perturbations during neurodevelopment (Hallgrímsson 1993). Fluctuating asymmetry has thus been employed as an indicator of nonspecific developmental stress (Clarke and McKenzie 1992; Townsend 1983) and linked to underlying genetic and environmental instabilities (Bradley 1980). Modeling the trajectories of FA through typical development and aging in neuroimaging-derived phenotypes could shed light on how population variation in symmetry may be associated with risk for neurodevelopmental and neurodegenerative disorders.

Despite growing evidence linking structural asymmetry to both normal and atypical brain function, relatively few studies have modeled asymmetry directly (Roe, Vidal-Piñeiro, et al. 2021; Roe, Vidal-Pineiro, et al. 2023; Maingault et al. 2016; Postema, Van Rooij, et al. 2019; G. Li et al. 2015; Hill, Dierker, et al. 2010; Korbmacher et al. 2024), especially across the full developmental trajectory from prenatal stages through to old age. This gap is particularly relevant in the context of recent work demonstrating the utility of normative modeling approaches for tracking brain development and aging (‘brain charts’) (Bethlehem et al. 2022; Rutherford et al. 2022; Y. Yu et al. 2024; Z. Yang et al. 2024). Brain charts offer an analogy to pediatric growth charts, used in routine clinical care to benchmark measurements of height, weight, and head circumference against population norms. The basic assessment of normalized (per)centile scores from pediatric growth charts allows for early detection and intervention in cases of atypical development of height, weight, or head circumference. In contrast, brain charts (of neuroimaging-derived phenotypes such as volumetrics) could be used to detect atypical neurodevelopmental or neurodegenerative patterns by positioning individual measurements against age-specific norms (Bedford, Seidlitz, and Bethlehem 2022). Given longstanding hypotheses linking brain asymmetry with neuropsychiatric pathologies (Esteves et al. 2021; Ocklenburg et al. 2024), charts of brain asymmetry may be particularly valuable for translational neuroimaging studies.

Here, we propose a population modeling approach to study structural brain asymmetry over the lifespan. Because large and heterogeneous samples are required for replicable and generalizable neuroimaging models (Marek and Laumann 2025; K. Kang et al. 2024), we have aggregated 128 primary studies (including multiple fully open datasets (Nielson et al. 2023; Zareba et al. 2022; Snoek et al. 2021; Bilder et al. 2020; Nárai et al. 2022; Nastase et al. 2025; Spreng et al. 2022; Nugent et al. 2025; Snoek et al. 2020a; Snoek et al. 2020b; Richardson et al. 2019; Garza-Villarreal et al. 2021; J. Wang et al. 2021; Tisdall and Mata 2023; Nielson et al. 2023; Fuchs et al. 2023; Botvinik-Nezer et al. 2023; Bellec et al. 2017; Di Martino et al. 2014; Tanaka et al. 2021; Reynolds et al. 2020); see **SI Fig. 1** and **SI Table 1** for details) comprising 177,701 scans from 138,231 individuals to cover the age range from prenatal development to old age (**SI Fig. 2**), spanning a wide range of image acquisition protocols including ultra low- to high-field MRI. The ultimate goal of these models is to identify neurodevelopmental milestones in asymmetry development and to detect deviations in asymmetry centile scores in patients with psychiatric and neurological disorders. To this end, we first present trajectories of brain development with hemispheric specificity, which we then integrate into explicit models of lifespan asymmetry. In doing so, we release the first comprehensive set of brain charts capturing lifespan asymmetry trajectories across a range of phenotypes, while providing a major improvement over our previous brain charts to refine our understanding of typical and atypical brain maturation. The resulting lifespan atlas of hemispheric development offers novel benchmarks for both basic and clinical neuroscience, with implications for future research on lateralized brain functions in health and disease.

## Results

### Hemispheric differences in development

Using generalized additive models for location, scale and shape (GAMLSS; Stasinopoulos and Rigby (2008) and Borghi et al. (2006)), we derived hemisphere-specific trajectories of regional brain development with age, stratified by sex (**SI Fig. 3**,**4**), for cortical grey matter volume (GM), thickness (CT), surface area (SA), and subcortical volume (**SI Fig. 5-13**). These trajectories were characterized in terms of milestones of development: (i) the peak rate-of-growth; (ii) the peak of the trajectory; and (iii) the peak rate-of decline (**SI Fig. 14**; **SI Table 1**). In line with previous bilaterally-averaged results (Bethlehem et al. 2022), left and right regional tissue trajectories peaked largely in (early) childhood and adolescence (**Fig. 2A**), with left-right age differences in hemispheric peaks of up to 6 years across phenotypes (**Fig. 2B**; see **SI Fig. 15-16** and **SI Table 2** for further details). Prior work has suggested that the timing of brain development relates to vulnerability to decline in later life (Raz 2000). his concept of ‘retrogenesis’ or more coloqially ‘first-in-last-out’ suggests that the ageing brain matures in the reverse sequence of tissue maturation during developing, meaning that late maturing regions are particularly vulnerable to decline (Yeatman, Wandell, and Mezer 2014). We therefore studied the interplay between the peak rate of growth and decline for each phenotype. For grey matter volume, there was a negative relationship between the rate of growth and decline, such that regions that grow first tend to decline last in later life (**Fig. 2C**)). In contrast, the relationship for surface area was positive, such that regions that expand their surface early also decline earlier in later life (**SI Fig. 17**). The peak rate of growth in cortical thickness for all regions (except the cuneus and isthmus cingulate) was at the earliest available age in the aggregated dataset, with most regions showing a correspondingly early decline. While highly informative in their own right, pronounced hemispheric differences in regional trajectories also point to the potential utility of direct models of brain asymmetry across the lifespan. To identify canonical maturational profiles, we conducted a data-driven grouping of brain regions based on shared developmental trajectories. By clustering the grey matter volume trajectories, we identified spatially coherent clusters of development, with early-peaking clusters aligning with primary sensorimotor cortices, and later clusters mapping onto association and limbic areas, consistent with known cortical hierarchies (**Fig 2D**). Lastly, the trajectories allow us to derive subject-specific centile scores, and contrast deviations from the norm in cases and controls (see **SI Fig. 18** for details).

**Fig. 1:**
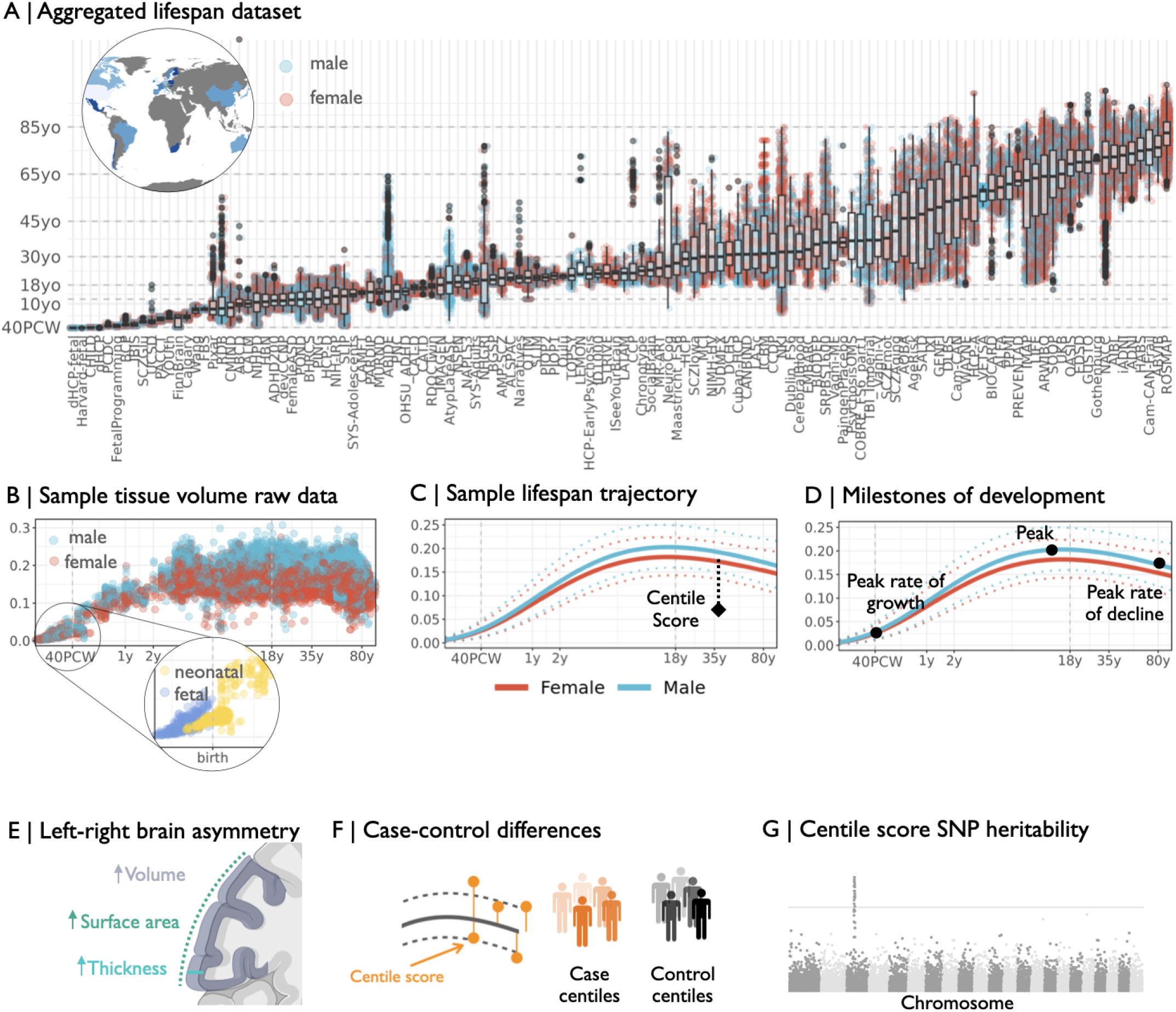
Generating asymmetry brain charts for the human lifespan: (A) We collected T1- and T2 weighted MRI data from 128 primary studies from 25 countries to form an aggregated dataset of 177,701 scans from 138,231 individuals, that collectively spanned the age range from mid-gestation to 102 years. The box-plots show the age distribution for each study, with individual points colored by sex. Term birth is indicated at 40 post conception weeks. The world map depicts the origin of the studies, with darker colors indicating an increased number of subjects from a given country. (B) Sample regions showing the raw gray matter volume of the right amygdala plotted for each cross-sectional scan as a function of age (log-scaled). Colors indicate sex. We further highlight the fact that we included overlapping samples of fetal scans and preterm born neonates to cover the early life period. (C) We estimated normative trajectories of tissue development using GAMLSS, stratified by sex, with site-specific batch effects, resulting in sex-specific lifespan trajectories of development of the median for each phenotype. We show the non-linear trajectory of the sample region’s median gray matter volume (with 2.5 and 97.5% centiles denoted as dotted lines) as a function of age (log-scaled). (D) Next, we estimated trajectories of normative asymmetry development, using GAMLSS. (E) We estimated individual deviations from normative development and estimated group differences in these centile scores in a number of neuropsychiatric and neurodegenerative disorders. (F) Lastly, we quantified SNP heritability and genetic correlations of deviations of regional and asymmetry centile scores.

**Fig. 2:**
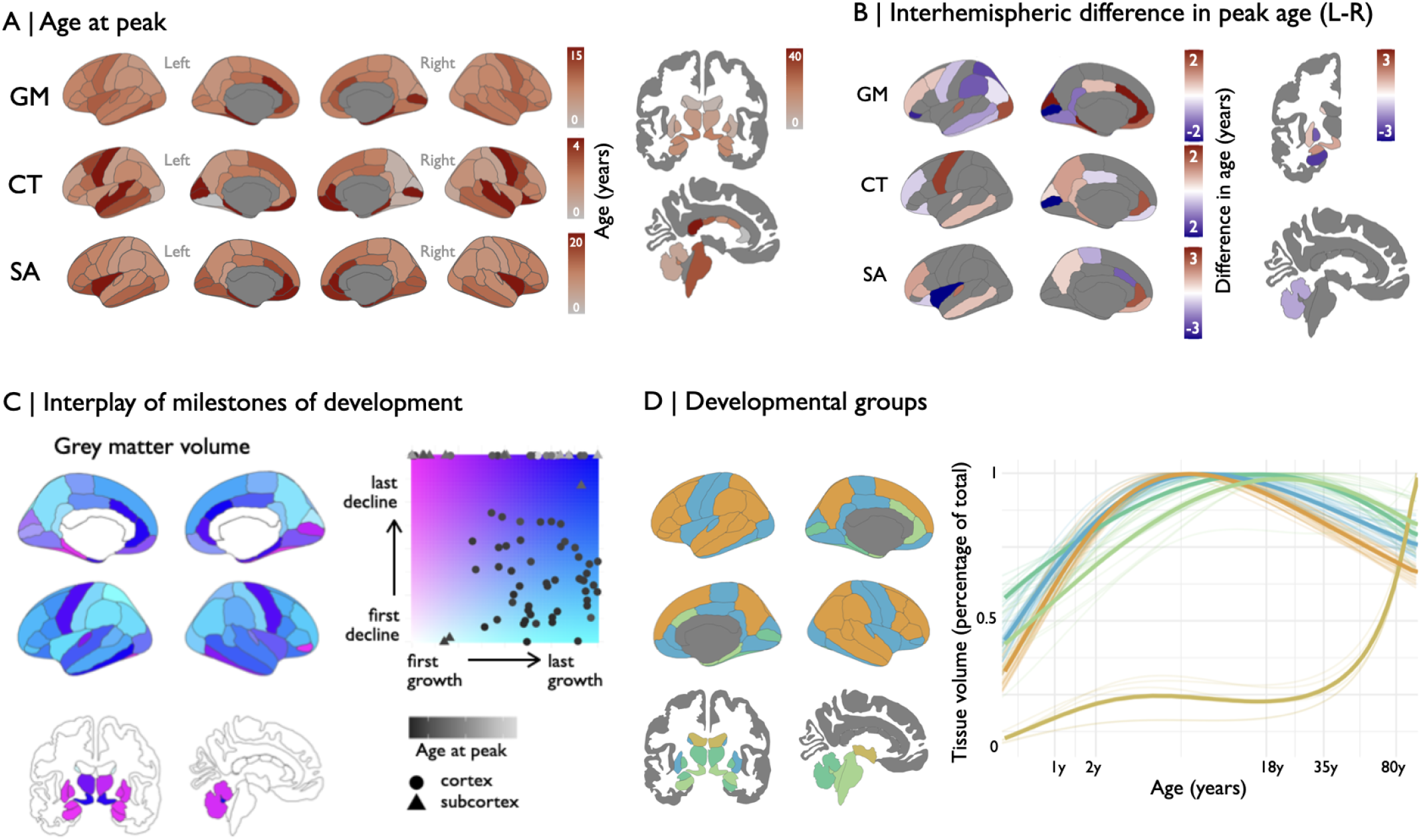
Hemispheric differences in developmental milestones: (A) For each bilateral region, we estimated the age at which three milestones of development were reached: the peak rate of change, the peak of the trajectory, and the peak rate of decline. Here, we illustrate the trajectory peaks. Cortical peaks vary by tissue type and region, with earlier peaks in sensorimotor regions and later peaks in association areas. (B) The spatial maps of developmental milestones reveal striking hemispheric differences in the timing of milestones of development. Regional differences in the timing of peak grey matter volume are visualized as a direct left–right contrast, highlighting systematic hemispheric divergence in developmental timing (in years). (C) The joint mapping of the peak rate of growth and decline uncovers distinct developmental phenotypes, separating regions with rapid early growth and slow decline (“First in last out”, pink) from those with the opposite pattern (see **SI Fig. 15** for other phenotypes). (D) K-means clustering of developmental trajectories (5-cluster solution) identifies canonical maturational profiles, reflecting known cortical hierarchies. Early-developing clusters include primary cortices; later trajectories capture association and limbic regions. Cluster membership was largely symmetric across hemispheres, though asymmetries in trajectory shape were evident within clusters. Mean cluster trajectories are overlaid on top of regional trajectories.

### Lifespan trajectories of asymmetry development

We derived an index of regional brain asymmetry in each phenotype as

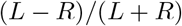

where *L* and *R* are the respective values of the regional phenotype in the left and right hemisphere. Accordingly, an asymmetry score greater than zero indicated a left-dominant region, whereas a score smaller than zero indicated a right-dominant region (**Fig. 3A**). We modeled lifespan development of asymmetry using GAMLSS; see **Supplementary information** for details).

**Fig. 3:**
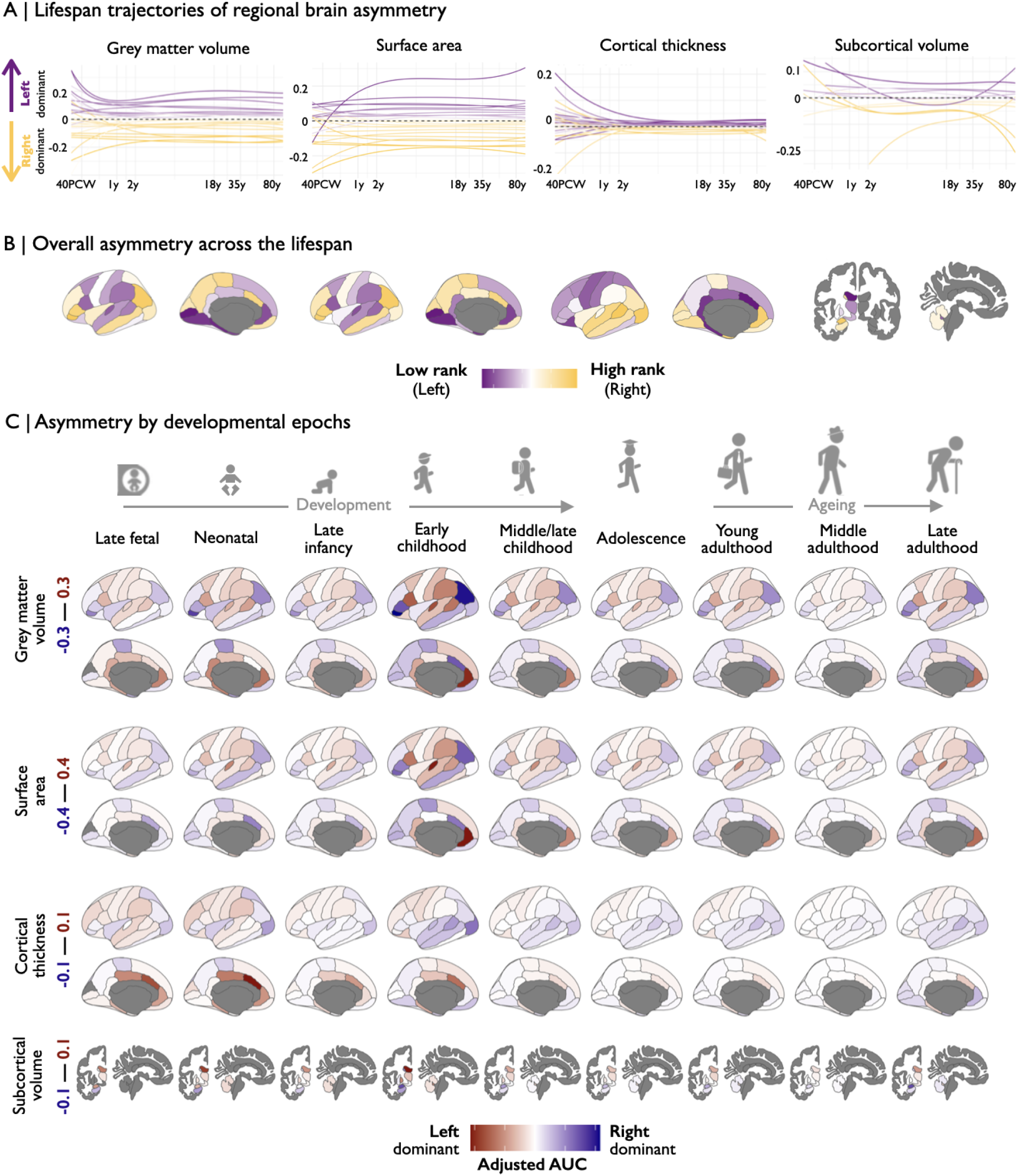
Lifespan development of brain asymmetry: (A) We estimated trajectories of regional asymmetry across the cortex and subcortex. A negative asymmetry index indicates rightward asymmetry, whereas a positive index indicates leftward asymmetry. (B) We then ranked the trajectories by their signed overall lifespan asymmetry. (C) Lastly, we estimated the magnitude of asymmetry as the area under the curve of the asymmetry trajectory in each of nine prior developmental windows ranging from conception to old age (H. J. Kang et al. 2011; **SI Fig. 22**): *late fetal* (24-38 PCW), *early infancy* (0-6 months), *late infancy* (6 months - 1 year), *early childhood* (1-6 years), *mid to late childhood* (6-12 years), *adolescence* (12-20 years), *young adulthood* (20-40 years), *mid adulthood* (40-60 years), *late adulthood* (older than 60 years). We normalized the resulting value by the length (log) of the time window. The color scale is adjusted for each phenotype and min/max values are indicated on the left. This analysis highlighted crucial development of asymmetry in relevant developmental epochs, i.e. language regions show high asymmetry during a period in life during which language is typically developed.

Across the lifespan, regional patterns of hemispheric asymmetry exhibit distinct developmental trajectories. Asymmetries in surface area and grey-matter volume either emerge or consolidate during the first two years of life – with left-dominant association regions (e.g. inferior parietal and superior temporal cortices) gradually amplifying their lateralization – and then remain relatively stable through adulthood and into old age; cortical thickness asymmetries peak neonatally but decline steeply over the first few postnatal years and approach symmetry by adolescence, possibly suggesting an early “fine-tuning” of cortical architecture (**Fig. 3A**; see **SI Fig. 19-21** for bootstrapped trajectories).

To highlight overall trends in asymmetry, we estimated the ranked average area under the curve of the trajetories as an indicator of overall lifetime asymmetry (**SI Table 3**; **Fig. 3B**; **SI Fig. 22**). To add temporal resolution to these analyses, we estimated the average asymmetry as the area under the curve of the trajectory in a given age bin (**SI Table 4**). This analysis highlighted asymmetries in key regions involved in the development of cognitive skills at relevant time periods. For example auditory and language regions (e.g., transverse temporal, pars opercularis and supramarginal cortex) developed a strong leftward asymmetry during early childhood – the critical period for language acquisition (**Fig. 3C**; Panszczyk et al. 2025). In addition, to evaluate patterns of asymmetry change across the lifespan, we estimated the pairwise spatial correlation of these maps of epoch-specific asymmetry and identified two clusters of asymmetry development (early vs late; (**SI Fig. 23**)): early asymmetry prominence, corresponding primarily to association cortical areas that exhibit relatively later peaks in bilateral growth, and late asymmetry prominence, corresponding primarily to sensory areas that reach their peak growth earlier (**SI Fig. 24**)).

We next grouped all subcortical and cortical regions based on their average asymmetry across the lifespan, highlighting distinct temporal patterns: regions with consistent leftward or rightward asymmetry, and regions with developmental reversals – left-to-right or right-to-left shifts – in asymmetry (**Fig. 4A**, *top*). Cortical surface maps of these groupings highlighted spatially coherent clusters of asymmetric development (**SI Table 5**). Association cortices, such as lateral frontal and temporal regions, are prominently represented among the early leftward-asymmetric areas. In contrast, primary sensory and motor regions — including the postcentral gyrus and visual cortex — tend to exhibit later-onset rightward asymmetry (**Fig. 4A**). Regions with developmental reversals are notably distributed across both association and primary cortices (**Fig. 4A**, *bottom*, **SI Fig. 24-25**). Interestingly, regions with greatest magnitude of changes in asymmetry over the lifespan showed an anatomical overlap with the two phylogenetically primitive areas (paleo-cortex and archicortex) of the dual origin theory of cortical evolution (Pandya, Petrides, and Cipolloni 2015; Abbie 1940) (**SI Fig. 26**).

**Fig. 4:**
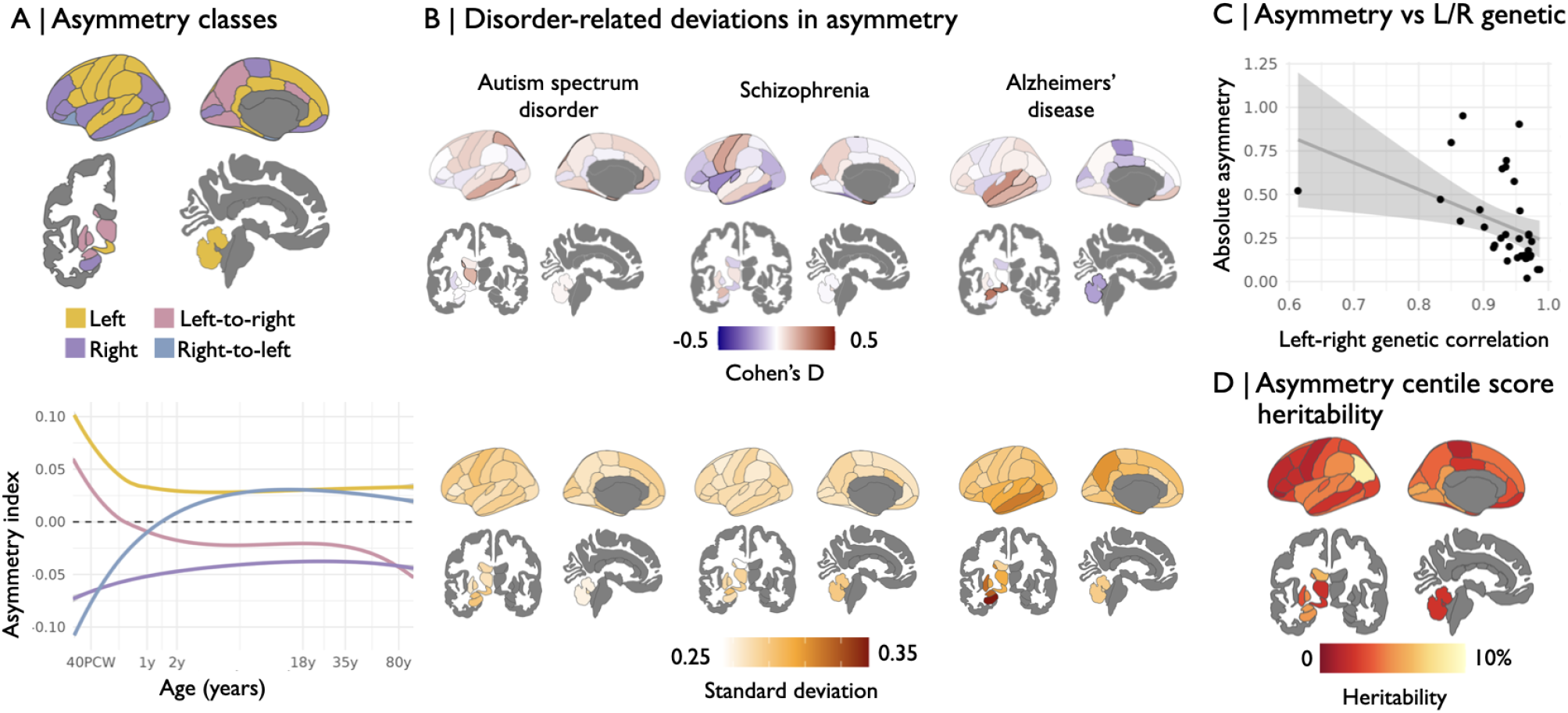
Grey matter volume and subcortical asymmetry centile scores: (A) (*top*) We assigned cortical and subcortical grey matter regions into four asymmetry classes: (i) left dominant; (ii) right dominant; (iii) left-to-right changing; and (iv) right-to-left changing regions. (*bottom*) We then estimated the average within-group trajectory for each class. (B) (*top*) We estimated case-control differences in grey matter volume asymmetry centile scores (black noutlines highlight regions with significant case control differences *P*_*F DR*_ *<* 0.05). (*bottom*) Standard deviation in centile scores of patients with diagnoses of neuropsychiatric disorders (*ρ*_*GM*_ = −0.63, *P*_*spin*_ *<* 0.01). (C) We observe higher asymmetry in regions with lower left-right genetic correlation. (D) We estimated the heritability of grey matter volume asymmetry centile scores using LDSC regression in the UK Biobank sample.

### Disorder-related and other group-level differences in asymmetry

We further derived subject-specific deviations from reference models of asymmetry development and aging, i.e. asymmetry centile scores. We then estimated case-control differences in asymmetry centile scores for individuals with autism spectrum disorder, schizophrenia, and Alzheimer’s disease (**Fig. 4B**; **Fig. 27-28**; **SI Table 6**). The use of centile scores allows us to combine diverse and unmatched cohorts (see **SI Table 2**). Of note, case-control differences in mean group-level asymmetry centile scores were less widespread than for centile scores derived from each hemisphere separately, which may be more sensitive to global alterations in brain size (see **SI Fig. 18A** for left/right regional case-control maps). As expected, we observed far more extreme centile scores in cases compared to controls, most pronounced in temporal lobe regions in Alzheimer’s disease (**Fig. 28**). However, there are pronounced changes in the distribution of asymmetry centile scores in cases compared to controls. More specifically, we observed far more extreme centile scores in cases compared to controls, quantified as a greater standard deviation in centile scores in individuals with neurodevelopmental and neurodegenerative disorders (**SI Fig. 30**).

Given that handedness has been associated with altered patterns of asymmetry and risk for neuropsychiatric disorders (Webb et al. 2013; Sha, Pepe, et al. 2021), we estimated effects of handedness on asymmetry centile scores (**SI Fig. 31A, 32**). We found that non-right-handers across the lifespan exhibited a trend of reduced cortical and subcortical grey matter volume asymmetries, compared to right-handers (**SI Fig. 31B**).

### Genetic association of subject-specific deviations

To the extent that asymmetry is driven by different genetic factors influencing left and right brain development, more asymmetric regions would be expected to vary more in terms of the genetic factors that influence left-right homologs, compared to more symmetric regions. Using UK Biobank data, we estimated the genetic correlation between left and right centile scores for all phenotypes, which quantifies the proportion of interregional covariance due to shared genetic factors (**SI Table 7**). We found that regions with lower left-right genetic correlation tended to show greater asymmetry across the lifespan (*ρ*_*GM*_ = −0.63, *P*_*spin*_ *<* 0.01, **Fig. 4C, SI Table 8**; see **SI Fig. 33** for other phenotypes).

However, prior research has indicated low (Warrier et al. 2023) to modest (Sha, Schijven, et al. 2021) heritability of asymmetry. We assessed genetic variation associated with individual deviations from asymmetry. Most regions showed modest, but significant (*P*_*F DR*_ *<* 0.05), SNP-based heritability values between 2–6%, indicating that common genetic variants account for a small but non-negligible proportion of individual differences in asymmetry trajectories (**Fig. 4D**; see **SI Fig. 34** for all phenotypes). The lower SNP-based heritability in primary sensory and motor regions suggests that these regions may be more shaped by environmental or stochastic developmental factors rather than by inherited variation. In contrast, a number of language regions, including the superior temporal gyrus, transversetemporal, and supramarginal cortices, as well as multiple regions in the association cortex stand out as having higher SNP-based heritability (Correlation with sensorimotor-association: *ρ* = −0.4, *P*_*spin*_ *<* 0.05; see **SI Fig. 35**).

### Promoting access to lifespan trajectories

To promote open science and facilitate future research, we provide the complete set of regional lifespan trajectories and asymmetry trajectories as an openly accessible resource at:http://www.brainchart.io/ (upon publication). These models cover cortical thickness, surface area, grey matter volume, and subcortical volumes with bilateral resolution across the entire lifespan, from prenatal stages to old age. Users can query individual regions of interest, visualize developmental trajectories via an interactive interface, and generate asymmetry centile scores across a range of harmonized image acquisition protocols including for out-of-sample datasets. These reference brain charts are available to support replication, benchmarking, and translational applications across developmental, and clinical neuroscience studies. We further publish relevant derived maps, including case-control effect maps and average asymmetry maps. Lastly, we release the centile score GWAS summary statistics to support further exploration of genetic variants associated with deviations from normative development.

## Discussion

Here we provide lifespan reference models of structural brain asymmetry, spanning from mid-gestation to old age, and encompassing diverse neuroimaging phenotypes. Harnessing a large, globally distributed aggregation of MRI datasets by the Lifespan Brain Chart Consortium (LBCC), we significantly improve and extend upon prior LBCC brain charts (Bethlehem et al. 2022), providing a resource to map typical and atypical patterns of lateralized brain development and asymmetry.

Beyond characterizing structural asymmetry as a normative feature of brain development, our findings align with broader theories of developmental canalization and neuroplasticity. The existence of dynamic, region-specific trajectories supports a model in which lateralization is shaped by both genetically predetermined schedules and environmentally contingent tuning during critical periods (Wu et al. 2025; Van Valen 1962; Hallgrímsson 1993). Notably, we found that regions exhibiting the greatest change in asymmetry across the lifespan align with the dual-origin map of cortical evolution (Pandya, Petrides, and Cipolloni 2015; Abbie 1940). This trend may suggest that these areas lie along a developmental axis especially susceptible to fluctuating asymmetry—where accumulated stochastic influences during sensitive periods may contribute to both structural variability and functional specialization (A. Alexander-Bloch, Giedd, and Bullmore 2013; Hill, Inder, et al. 2010).

Reference brain charts of asymmetry allow for precise quantification of individually-specific deviations from typical asymmetry trajectories across development and aging. When applied to clinical populations, we found that case-control differences in mean asymmetry centile scores were modest and spatially localized, especially compared to centile scores calculated in each hemisphere separately, which are more sensitive to global alterations in brain size. However, we observed marked increases in the variability of asymmetry centile scores among individuals with autism, schizophrenia, and Alzheimer’s disease, consistent with they hypothesis that increased developmental or environmental stress leads to greater heterogeneity in brain lateralization among individuals with these disorders. These findings support longstanding hypotheses that altered asymmetry—and specifically, increased developmental instability reflected in fluctuating asymmetry—may be a transdiagnostic feature of neurodevelopmental and neurodegenerative disorders (Postema, Van Rooij, et al. 2019; Roe, Vidal-Piñeiro, et al. 2021). From a clinical translation perspective, our reference models have the potential to inform precision diagnostics. This could be particularly valuable in stratifying risk during early neurodevelopmental assessments, or in tracking progressive asymmetry shifts in conditions such as Alzheimer’s disease, where early left-right volumetric imbalances have been proposed as prodromal indicators (Roe, Vidal-Piñeiro, et al. 2021; Postema, Hoogman, et al. 2021; Sha, Van Rooij, et al. 2022; Schijven et al. 2023).

Our genetic analyses demonstrated that lower genetic correlation between left and right regional brain measures was associated with greater structural asymmetry. The results are consistent with prior work indicating that brain asymmetry is, at least party, the result of genetically decoupled developmental programs for the two hemispheres (Sha, Schijven, et al. 2021; Warrier et al. 2023). Furthermore, regions with stronger asymmetries exhibited lower genetic coupling, highlighting the potential for both genetically regulated and environmentally modulated contributions to lateralization, and offering a compelling platform to empirically evaluate developmental instability in asymmetry trajectories.

It is worth noting that the growth charts presented here, although analogous to pediatric growth charts, only demonstrate the feasibility of deriving lifespan trajectories of asymmetry development. Future work will have to explore how asymmetry centile scores can support clinical practice (Bedford, Seidlitz, and Bethlehem 2022). Key next steps include extending these charts to examine age-by-sex interactions and incorporating longitudinal data to disentangle within-individual variability and better characterize the origins of fluctuating asymmetry. Such data could help differentiate transient, developmentally constrained asymmetries from more stable, trait-like patterns, while capturing periods of heightened plasticity or vulnerability to environmental or pathological factors. Longitudinal trajectories would further clarify the contribution of left-right differences, including whether asymmetry reflects divergent rates of maturation or degeneration between hemispheres or dynamic shifts in inter-hemispheric balance across the lifespan. We further note that our sample reflects the research landscape of available neuroimaging datasets, which is strongly biased towards western populations (Kopal, Uddin, and Bzdok 2023). True translational potential requires population diversity (Sun et al. 2025). Recent work has demonstrated the ability to construct growth charts from clinically acquired data (Schabdach et al. 2023; Mandal et al. 2024), a development which may facilitate the integration of more diverse populations in neuroimaging work, since each year millions of clinical brain scans are routinely acquired around the world.

Widespread efforts are required to further the democratization of brain MRI research and clinical data. We demonstrate the adaptability and translational potential of centile modeling to varying image resolutions and scanner environments using ultra-low-field MRI systems deployable outside of traditional neuroimaging research and clinical settings (Váša, Bennallick, et al. 2024) (see **SI Fig. 36**). As global neuroimaging consortia grow, difficult questions must be addressed regarding potential benefits and risks of demographically-calibrated brain charts to ensure generalizability and equity in neurodevelopmental research (Garcini et al. 2022). Brain charts tailored to specific genetic syndromes or clinical conditions (such as premature birth) may also increase sensitivity to detect clinically-relevant differences as has been shown in pediatric growth charts (Zemel et al. 2015; Cheung et al. 2024; Fenton and Kim 2013).

Lastly, to facilitate the implementation of asymmetry brain chart models, we provide an openly available resource for future research (http://www.brainchart.io/, updated upon publication). The resource includes code, summary statistics, and reference centile curves that can be applied to independent datasets, enabling individualized assessments of asymmetry for clinical or developmental studies. In doing so, our study provides both a framework and a toolset for investigating brain asymmetry in health and disease.

## Methods

### Preprocessing

The primary studies were available either as open raw data or only as derived phenotypes. For the former, raw imaging data was processed using FreeSurfer for older data, in particular data included in Bethlehem et al. 2022 and subsequently SynthSeg (Billot et al. 2023) for more recently acquired studies. Prior work has demonstrated the reliability of constructing brain-charts from both these processing pipelines (Schabdach et al. 2023). The exception are the dHCP and dHCP-fetal studies, which were processed using their respective published pipelines and registered to the M-CRIB neonatal cortical and subcortical brain atlas (Alexander et al. 2017) using surface registration (Makropoulos et al. 2018; Silva et al. 2025). All other data was parcellated using the Desikan Killiany atlas (Desikan et al. 2006). Due to the number of images, automatic quality control was performed using the Euler Index excluding scans exceeding a threshold. Details of the processing pipelines are in the Supplementary Material.

### Site aggregation

Within the data structure there are three levels: individual, site, and study (four if repeated scans on an individual are included). That is, many studies are actually multi-site studies – with variations in protocol/scanner technology across sites within the same study. As such, our analysis uses only two levels: individual and site, this is to capture as much heterogeneity through site-level effects. However, sites with too few individuals are problematic in our analyses. To address this, small sites within a study are aggregated into a pseudosite for each study. This retains the most data while accounting for site/study heterogeneity.

### Lifespan trajectories

We estimate lifespan trajectories using the Generalized Additive Models for Location, Scale and Shape (GAMLSS) framework. Broadly this is a highly flexible framework for modelling outcome distributions with multiple parameters, enabling the direct modelling of multiple statistical moments (i.e. mean, variance, skew, and kurtosis) where each parameter may depend on dependent variables (e.g. age and sex) and random effects (e.g. site-level effects). An appropriate outcome distribution is determined using standard model selection (i.e. Akaike Information Criteria (AIC)) based on an exploratory subset of the data. Given the GAMLSS outcome distribution, the form of the parameter component regressions must be determined (a two-stage process is necessary to avoid a prohibitive computational burden). GAMLSS allows a very flexible form for the component regressions, following (Bethlehem et al. 2022) we use fractional polynomial models of age effects. As discussed above, the GAMLSS components also include a site-level random effect. To address the robustness of the models to specific studies we perform a leave-one-study-out analysis (note study, not site). To quantify the uncertainty in the models we perform a non-parametric stratified bootstrap (ensuring the bootstrap replicates maintain the sex and site proportions); as expected this is a substantial computational burden but is necessary to understand the uncertainty of key summary statistics.

### Centile score estimation

A consequence of the GAMLSS framework with higher moments (i.e. skew) is that the “standard” summary of z-/t-scores is no longer meaningful or valid. However, we can calculate the centile for each observation based on the parameterised outcome distribution. We can also calculate a rescaled “centile standard score” via the quantile of the normal distribution of the centile. Hence, we consider centiles and centile standard scores as a robust and comparable summary across phenotypes across the lifespan.

### Hemisphere-specific regional models

Initially we fit GAMLSS models to every region across all phenotypes. These lifespan trajectories are used to derive developmental milestones, such as peak development.

### Regional asymmetry models

The focus of our paper is on asymmetry, defined as the normalised ratio of the left and right region pair difference. This induces a non-linear scale where 1 and −1 represent extreme asymmetry while 0 corresponds to a symmetric region.

### Disorder-related differences

The GAMLSS models are estimated using healthy controls. This naturally leads to investigating case–control differences using clinical groups. Centiles for asymmetry are compared for case–control differences.

### SNP heritability analyses

We conducted GWAS analyses of left/right regional, as well as asymmetry centile scores for UKB participants across all phenotypes using FastGWA (v1.93) (Jiang et al. 2019) with the following covariates: age, age^2^, sex, age *×* sex, age^2^ *×* sex, imaging center, first 40 genetic principal components, mean framewise displacement (as obtained from the accompanying resting-state fMRI scan), maximum framewise displacement (as obtained from the accompanying resting-state fMRI scan) and Euler Index (Rosen et al. 2018) as covariates. We computed genetic correlations and SNP-based heritability for the GWAS statistics using LDSC (v1.01) (B. Bulik-Sullivan et al. 2015; B. K. Bulik-Sullivan et al. 2015), using LD weights from the North West European populations. Please refer to **Supplementary Material** for details on genetic quality control.

### Correction for spatial autocorrelation

To control for potential effects of spatial autocorrelation affecting the relationship between any two cortical maps (A. F. Alexander-Bloch et al. 2018), for each spatial correlation of two such maps, we report both a non-parametric *P* − *value* corresponding to the Spearman correlation (*ρ*), as well as a *P* − *value* derived from the more conservative spin-test permutation (Váša, Seidlitz, et al. 2018). Note that when correlations are estimated including subcortical regions, no such correction can be applied.

### Further details

Refer to the **Supplementary Material** for further details and in-depth discussion of practical aspects of the above methods.

## Code Availability

We generated the models using the GAMLSS model code in https://github.com/brainchart/Lifespan. Code for analyses specific to this paper can be accessed at https://github.com/BGDlab/asymmetry-braincharts/ (upon publication). This repository specifically also includes all models described here and a tutorial on how to generate out-of-sample centiles using users’ own data. The repository will be made public upon publication, until then reviewers can access it using the login details for a temporary github account we created.

## Data Availability

Data used in the preparation of this article were obtained from the Lifespan Brain Chart consortium (LBCC), which includes multiple publicly-available MRI datasets. These datasets are subject to their respective data access and sharing policies and can be accessed via the following websites: ABCD (https://abcdstudy.org/); the International Neuroimaging Data-Sharing Initiative (https://fcon_1000.projects.nitrc.org/); the Laboratory of NeuroImaging (https://loni.usc.edu/); the BarcelonaBeta Brain Research Center (https://www.barcelonabeta.org/en); the Alzheimer’s Disease Repository Without Borders (ARWiBo) (https://www.arwibo.it); CamCAN (http://www.mrc-cbu.cam.ac.uk/datasets/camcan/); NIMH Data Archive (https://nda.nih.gov/); the Open Science Framework (https://osf.io/); The Global Alzheimer’s Association Interactive Network (https://www.gaain.org/); CODE (https://www.braincode.ca/); Brain-Science Data Bank (https://www.scidb.cn/en); Longitudinal Online Research and Imaging System (https://www.loris.ca/); the Human Connectome Project (https://www.humanconnectome.org/); Information eXtraction from Images (IXI) (https://brain-development.org/ixi-dataset/); figshare (https://figshare.com/); Collaborative (COINS) Nathional Informatics and Neuroimaging Suite (https://coins.trendscenter.org/); Alzheimer’s Coordinating Center (NACC) (https://naccdata.org/); NeuroImaging Tools and Resources Collaboratory (NITRC) (https://www.nitrc.org/); Neuroscience in Psychiatry Network (NSPN) (https://nspn.org.uk/); Open Access Series of Imaging Studies (OASIS) (https://sites.wustl.edu/oasisbrains/); OpenNeuro (https://openneuro.org/); BICR Resource (https://bicr-resource.atr.jp/); UK Biobank (UKB), a major biomedical database (https://www.ukbiobank.ac.uk/); YOUth Cohort Study (https://www.uu.nl/en/research/youth-cohort-study/data-access); and the Open-Pain project (https://www.openpain.org). Details of each dataset contained within the LBCC are available in **SI Table 1**.

Data used in the preparation of this article were obtained from the Alzheimer’s Disease Neuroimaging Initiative (ADNI) database (adni.loni.usc.edu). The ADNI was launched in 2003 as a public-private partnership, led by Principal Investigator Michael W. Weiner, MD. The primary goal of ADNI has been to test whether serial magnetic resonance imaging (MRI), positron emission tomography (PET), other biological markers, and clinical and neuropsychological assessment can be combined to measure the progression of mild cognitive impairment (MCI) and early Alzheimer’s disease (AD).

ICBM data are disseminated by the Laboratory of Neuro Imaging at the University of Southern California.

Data used in the preparation of this work were obtained from the Mind Clinical Imaging Consortium database through the Mind Research Network (www.mrn.org).

## Acknowledgments

LD, MG, SK, JS, JS, and AAB were supported by R01MH132934, R01MH133843, R01MH134896, and the CHOP Research Institute. T.S. was supported by NIH R01MH113550, NIH R01MH120482, NIH R01MH112847, NIH U24NS130411and the Penn-CHOP Lifespan Brain Institute. M.M.V. was supported by an MRC CDA (MR/Y011384/1).

We would like to acknowledge the individuals and organizations that have made the data used for this research available, including the following consortia: Lifespan Brain Chart Consortium (LBCC); Aging Brain: Vasculature, Ischemia, and Behavior Study (ABVIB); Alzheimer’s Disease Neuroimaging Initiative (ADNI); Australian Imaging, Biomarkers and Lifestyle (AIBL); Alzheimer’s Disease Repository Without Borders (ARWiBo); Biomarkers of Cognitive Decline among Normal Individuals (BIOCARD); Centre for Attention Learning and Memory (CALM); Cambridge Center for Ageing and Neuroscience (Cam-CAN); Chinese Color Nest Project (CCNP); Centers of Biomedical Research Excellence (COBRE); Consortium on Vulnerability to Externalizing Disorders and Addictions (cVEDA); Harvard Aging Brain Study (HABS); International Consortium for Brain Mapping (ICBM); IMAGEN; Neuroscience in Psychiatry Network (NSPN); Province of Ontario Neurodevelopmental Disorders (POND) Network; Pre-symptomatic Evaluation of Experimental or Novel Treatments for Alzheimer Disease (PREVENT-AD); and Vietnam Era Twin Study of Aging (VETSA). Lists of members and their affiliations appears in the Supplementary Information. The consortium investigators who designed and implemented these studies and/or provided data did not necessarily participate in the analysis or writing of this report. This manuscript reflects the views of the authors and may not reflect the opinions or views of the investigators or funders providing data, including those acknowledged below.

YOUth is funded through the Gravitation program of the Dutch Ministry of Education, Culture, and Science and the Netherlands Organization for Scientific Research (NWO grant number 024.001.003). A complete listing of the study investigators and study management can be found at https://www.uu.nl/en/research/youth-cohort-study/about-us/who-is-involved. YOUth investigators and management designed and implemented the study and/or provided data but did not necessarily participate in the analysis or writing of this report. This manuscript reflects the views of the authors and may not reflect the opinions or views of the YOUth study investigators or YOUth management.

We would like to acknowledge the individuals and organizations that have made Data used for this research available including CAM-BIND, the Ontario Brain Institute, the Brain-CODE platform, and the Government of Ontario.

The SRPS1600 dataset was provided by the DecNef Department at the Advanced Telecommunication Research Institute International, Kyoto, Japan.

Data used in the preparation of this article were obtained from the Adolescent Brain Cognitive Development (ABCD) Study (https://abcdstudy.org), held in the NIMH Data Archive (NDA). This is a multisite, longitudinal study designed to recruit more than 10,000 children age 9-10 and follow them over 10 years into early adulthood. The ABCD Study® is supported by the National Institutes of Health and additional federal partners under award numbers U01DA041048, U01DA050989, U01DA051016, U01DA041022, U01DA051018, U01DA051037, U01DA050987, U01DA041174, U01DA041106, U01DA041117, U01DA041028, U01DA041134, U01DA050988, U01DA051039, U01DA041156, U01DA041025, U01DA041120, U01DA051038, U01DA041148, U01DA041093, U01DA041089, U24DA041123, U24DA041147. A full list of supporters is available at https://abcdstudy.org/federal-partners.html. A listing of participating sites and a complete listing of the study investigators can be found at https://abcdstudy.org/consortium_members/. The ABCD data repository grows and changes over time. The ABCD data used in this report came from 10.15154/z563-zd24.

Data used in the preparation of this article were obtained from the Aging Brain: Vasculature, Ischemia, and Behavior Study database (https://ida.loni.usc.edu/login.jsp?project=ABVIB). The ABVIB study was launched in 1994 as a NIA-funded program project led by Principal Investigator Helena C. Chui MD. Data from a second study cohort was started in 2008-2013 and is included in the current database. Data collection and sharing for ABVIB was funded by the National Institutes on Aging (NIA) P01 AG12435.

We would like to acknowledge the primary funding source for the ABIDE II dataset (NIMH 5R21MH107045).

Data collection and sharing for this project was funded by the Alzheimer’s Disease Neuroimaging Initiative (ADNI) (National Institutes of Health Grant U01 AG024904) and DOD ADNI (Department of Defense award number W81XWH-12-2-0012). ADNI is funded by the National Institute on Aging, the National Institute of Biomedical Imaging and Bioengineering, and through generous contributions from the following: AbbVie, Alzheimer’s Association; Alzheimer’s Drug Discovery Foundation; Araclon Biotech; BioClinica, Inc.; Biogen; Bristol-Myers Squibb Company; CereSpir, Inc.; Cogstate; Eisai Inc.; Elan Pharmaceuticals, Inc.; Eli Lilly and Company; EuroImmun; F. Hoffmann-La Roche Ltd and its affiliated company Genentech, Inc.; Fujirebio; GE Healthcare; IXICO Ltd.; Janssen Alzheimer Immunotherapy Research & Development, LLC.; Johnson Johnson Pharmaceutical Research Development LLC.; Lumosity; Lundbeck; Merck Co., Inc.; Meso Scale Diagnostics, LLC.; NeuroRx Research; Neurotrack Technologies; Novartis Pharmaceuticals Corporation; Pfizer Inc.; Piramal Imaging; Servier; Takeda Pharmaceutical Company; and Transition Therapeutics. The Canadian Institutes of Health Research is providing funds to support ADNI clinical sites in Canada. Private sector contributions are facilitated by the Foundation for the National Institutes of Health (www.fnih.org). The grantee organization is the Northern California Institute for Research and Education, and the study is coordinated by the Alzheimer’s Therapeutic Research Institute at the University of Southern California. ADNI data are disseminated by the Laboratory for Neuro Imaging at the University of Southern California.

Data used in the preparation of this article was obtained from the Australian Imaging Biomarkers and Lifestyle flagship study of ageing (AIBL) funded by the Commonwealth Scientific and Industrial Research Organisation (CSIRO) which was made available at the ADNI database (www.loni.usc.edu/ADNI).

ARWiBo data collection and sharing for this project was supported by the Italian Ministry of Health, under the following grant agreements: Ricerca Corrente IRCCS Fatebenefratelli, Linea di Ricerca 2; Progetto Finalizzato Strategico 2000-2001 “Archivio normativo italiano di morfometria cerebrale con risonanza magnetica (età 40+)”; Progetto Finalizzato Strategico 2000-2001 “Decadimento cognitivo lieve non dementigeno: stadio preclinico di malattia di Alzheimer e demenza vascolare. Caratterizzazione clinica, strumentale, genetica e neurobiologica e sviluppo di criteri diagnostici utilizzabili nella realtà nazionale,”; Progetto Finalizzata 2002 “Sviluppo di indicatori di danno cerebrovascolare clinicamente significativo alla risonanza magnetica strutturale”; Progetto Fondazione CARIPLO 2005-2007 “Geni di suscettibilità per gli endofenotipi associati a malattie psichiatriche e dementigene”; “Fitness and Solidarietà”; and anonymous donors.

This paper uses data collected in the Accelerating Medicines Partnership in Schizophrenia (AMP SCZ) project. AMP SCZ is supported by NIMH grants U24MH124629, U01MH124631, U01MH124639. The research was also funded in part by the Welcome Trust (220664/Z/20/Z).

The BCP was supported by grants: U01MH110274, R01MH104324.

Data were provided in part by the Brain Genomics Superstruct Project of Harvard University and the Massachusetts General Hospital, (Principal Investigators: Randy Buckner, Joshua Roffman, and Jordan Smoller), with support from the Center for Brain Science Neuroinformatics Research Group, the Athinoula A. Martinos Center for Biomedical Imaging, and the Center for Human Genetic Research. 20 individual investigators at Harvard and MGH generously contributed data to GSP.

BHRCS was supported with grants from the National Institute of Development Psychiatric for Children and Adolescent (INPD). Grant: Fapesp 2014/50917-0 - CNPq 465550/2014-2

The BIODEP study was sponsored by the Cambridgeshire and Peterborough NHS Foundation Trust and the University of Cambridge, and funded by a strategic award from the Wellcome Trust (104025) in partnership with Janssen, GlaxoSmithKline, Lundbeck and Pfizer.

Data collection and sharing for this project was provided by the Centre for Attention, Learning and Memory (CALM). CALM funding was provided by the UK Medical Research Council and University of Cambridge, UK. Data used in the preparation of this work were obtained from CALM resource – https://calm.mrc-cbu.cam.ac.uk/. The study protocol is reported in Holmes et al. 2019.

We would like to acknowledge the individuals and organizations that have made Data [and analysis] used for this research available including [ID program name/Data producer name], the Ontario Brain Institute, the Brain-CODE platform, the Government of Ontario, as well as [independent collaborator names].”

Data collection and sharing for this project was provided by the Cambridge Centre for Ageing and Neuroscience (CamCAN). CamCAN funding was provided by the UK Biotechnology and Biological Sciences Research Council (grant number BB/H008217/1), together with support from the UK Medical Research Council and University of Cambridge, UK.

The CHILD study was funded by the Autism Research Trust

The Drakenstein Child Health Study is funded by the Bill and Melinda Gates Foundation (OPP 1017641)

The Developing Human Connectome Project was supported by the European Research Council under the European Union Seventh Framework Programme (FP/2007-2013)/ERC Grant Agreement No. 319456

Data were provided in part by the Human Connectome Project, WU-Minn Consortium (Principal Investigators: David Van Essen and Kamil Ugurbil; 1U54MH091657) funded by the 16 NIH Institutes and Centers that support the NIH Blueprint for Neuroscience Research; and by the McDonnell Center for Systems Neuroscience at Washington University.

Research using Human Connectome Project for Early Psychosis (HCP-EP) data reported in this publication was supported by the National Institute of Mental Health of the National Institutes of Health under Award Number U01MH109977. The HCP-EP 1.1 Release data used in this report came from DOI: 10.15154/1522899.

FinnBrain was funded by Jane and Aatos Erkko Foundation, Signe and Ane Gyllenberg Foundation

GUSTO study is supported by the Singapore National Research Foundation under its Translational and Clinical Research (TCR) Flagship Programme and administered by the Singapore Ministry of Health’s National Medical Research Council (NMRC), Singapore - NMRC/TCR/004-NUS/2008; NMRC/TCR/012-NUHS/2014. Additional funding is provided by the Singapore Institute for Clinical Sciences, Agency for Science Technology and Research (A*STAR), Singapore; E.C. is supported by grants: NIMH P50163 MH081755, NIMH R01-MH036840, NIMH R01-MH110558, NIMH U01-MH108898, NIDCD R01DC016385

Data used in the preparation of this work were obtained from the International Consortium for Brain Mapping (ICBM) database (www.loni.usc.edu/ICBM). The ICBM project (Principal Investigator John Mazziotta, M.D., University of California, Los Angeles) is supported by the National Institute of Biomedical Imaging and BioEngineering. ICBM is the result of efforts of co-investigators from UCLA, Montreal Neurologic Institute, University of Texas at San Antonio, and the Institute of Medicine, Juelich/Heinrich Heine University - Germany.

IMAP study (PI (scientific): G Chetelat; PI (MD) V de La Sayette)) was funded by Programme Hospitalier de Recherche Clinique (PHRCN 2011-A01493-38 and PHRCN 2012 12-006-0347) and Agence Nationale de la Recherche (LONGVIE 2007). Dr Chetelat’s research including IMAP was also funded by Institut National de la Santé et de la Recherche Médicale (Inserm), Fondation Plan Alzheimer (Alzheimer Plan 2008-2012)

LIFE is funded by means of the European Union, by the European Regional Development Fund (ERDF) and by funds of the Free State of Saxony within the framework of the excellence initiative.

Data used in the preparation of this work were obtained in part from the Mind Clinical Imaging Consortium database through the Mind Research Network (www.mrn.org). The MCIC project was supported by the Department of Energy under Award Number DE-FG02-08ER64581. MCIC is the result of efforts of co-investigators from the University of Iowa, University of Minnesota, University of New Mexico, and Massachusetts General Hospital, who collected and shared the imaging data and demographic information.

The NACC database is funded by NIA/NIH Grant U01 AG016976. NACC data are contributed by the NIA-funded ADCs: P30 AG019610 (PI Eric Reiman, MD), P30 AG013846 (PI Neil Kowall, MD), P50 AG008702 (PI Scott Small, MD), P50 AG025688 (PI Allan Levey, MD, PhD), P50 AG047266 (PI Todd Golde, MD, PhD), P30 AG010133 (PI Andrew Saykin, PsyD), P50 AG005146 (PI Marilyn Albert, PhD), P50 AG005134 (PI Bradley Hyman, MD, PhD), P50 AG016574 (PI Ronald Petersen, MD, PhD), P50 AG005138 (PI Mary Sano, PhD), P30 AG008051 (PI Thomas Wisniewski, MD), P30 AG013854 (PI Robert Vassar, PhD), P30 AG008017 (PI Jeffrey Kaye, MD), P30 AG010161 (PI David Bennett, MD), P50 AG047366 (PI Victor Henderson, MD, MS), P30 AG010129 (PI Charles DeCarli, MD), P50 AG016573 (PI Frank LaFerla, PhD), P50 AG005131 (PI James Brewer, MD, PhD), P50 AG023501 (PI Bruce Miller, MD), P30 AG035982 (PI Russell Swerdlow, MD), P30 AG028383 (PI Linda Van Eldik, PhD), P30 AG053760 (PI Henry Paulson, MD, PhD), P30 AG010124 (PI John Trojanowski, MD, PhD), P50 AG005133 (PI Oscar Lopez, MD), P50 AG005142 (PI Helena Chui, MD), P30 AG012300 (PI Roger Rosenberg, MD), P30 AG049638 (PI Suzanne Craft, PhD), P50 AG005136 (PI Thomas Grabowski, MD), P50 AG033514 (PI Sanjay Asthana, MD, FRCP), P50 AG005681 (PI John Morris, MD), P50 AG047270 (PI Stephen Strittmatter, MD, PhD). The NACC database is funded by NIA/NIH Grant U24 AG072122. SCAN is a multi-institutional project that was funded as a U24 grant (AG067418) by the National Institute on Aging in May 2020. Data collected by SCAN and shared by NACC are contributed by the NIA-funded ADRCs as follows: Arizona Alzheimer’s Center - P30 AG072980 (PI: Eric Reiman, MD); R01 AG069453 (PI: Eric Reiman (contact), MD); P30 AG019610 (PI: Eric Reiman, MD); and the State of Arizona which provided additional funding supporting our center; Boston University - P30 AG013846 (PI Neil Kowall MD); Cleveland ADRC - P30 AG062428 (James Leverenz, MD); Cleveland Clinic, Las Vegas – P20AG068053; Columbia - P50 AG008702 (PI Scott Small MD); Duke/UNC ADRC – P30 AG072958; Emory University - P30AG066511 (PI Levey Allan, MD, PhD); Indiana University - R01 AG19771 (PI Andrew Saykin, PsyD); P30 AG10133 (PI Andrew Saykin, PsyD); P30 AG072976 (PI Andrew Saykin, PsyD); R01 AG061788 (PI Shannon Risacher, PhD); R01 AG053993 (PI Yu-Chien Wu, MD, PhD); U01 AG057195 (PI Liana Apostolova, MD); U19 AG063911 (PI Bradley Boeve, MD); and the Indiana University Department of Radiology and Imaging Sciences; Johns Hopkins - P30 AG066507 (PI Marilyn Albert, Phd.); Mayo Clinic - P50 AG016574 (PI Ronald Petersen MD PhD); Mount Sinai - P30 AG066514 (PI Mary Sano, PhD); R01 AG054110 (PI Trey Hedden, PhD); R01 AG053509 (PI Trey Hedden, PhD); New York University - P30AG066512-01S2 (PI Thomas Wisniewski, MD); R01AG056031 (PI Ricardo Osorio, MD); R01AG056531 (PIs Ricardo Osorio, MD; Girardin Jean-Louis, PhD); North-western University - P30 AG013854 (PI Robert Vassar PhD); R01 AG045571 (PI Emily Rogalski, PhD); R56 AG045571, (PI Emily Rogalski, PhD); R01 AG067781, (PI Emily Rogalski, PhD); U19 AG073153, (PI Emily Rogalski, PhD); R01 DC008552, (M.-Marsel Mesulam, MD); R01 AG077444, (PIs M.-Marsel Mesulam, MD, Emily Rogalski, PhD); R01 NS075075 (PI Emily Rogalski, PhD); R01 AG056258 (PI Emily Rogalski, PhD); Oregon Health and Science University - P30 AG008017 (PI Jeffrey Kaye MD); R56 AG074321 (PI Jeffrey Kaye, MD); Rush University - P30 AG010161 (PI David Bennett MD); Stanford – P30AG066515; P50 AG047366 (PI Victor Henderson MD MS); University of Alabama, Birmingham – P20; University of California, Davis - P30 AG10129 (PI Charles DeCarli, MD); P30 AG072972 (PI Charles DeCarli, MD); University of California, Irvine - P50 AG016573 (PI Frank LaFerla PhD); University of California, San Diego - P30AG062429 (PI James Brewer, MD, PhD); University of California, San Francisco - P30 AG062422 (Rabinovici, Gil D., MD); University of Kansas - P30 AG035982 (Russell Swerdlow, MD); University of Kentucky - P30 AG028283-15S1 (PIs Linda Van Eldik, PhD and Brian Gold, PhD); University of Michigan ADRC - P30AG053760 (PI Henry Paulson, MD, PhD) P30AG072931 (PI Henry Paulson, MD, PhD) Cure Alzheimer’s Fund 200775 - (PI Henry Paulson, MD, PhD) U19 NS120384 (PI Charles DeCarli, MD, University of Michigan Site PI Henry Paulson, MD, PhD) R01 AG068338 (MPI Bruno Giordani, PhD, Carol Persad, PhD, Yi Murphey, PhD) S10OD026738-01 (PI Douglas Noll, PhD) R01 AG058724 (PI Benjamin Hampstead, PhD) R35 AG072262 (PI Benjamin Hampstead, PhD) W81XWH2110743 (PI Benjamin Hampstead, PhD) R01 AG073235 (PI Nancy Chiaravalloti, University of Michigan Site PI Benjamin Hampstead, PhD) 1I01RX001534 (PI Benjamin Hampstead, PhD) IRX001381 (PI Benjamin Hampstead, PhD); University of New Mexico - P20 AG068077 (Gary Rosenberg, MD); University of Pennsylvania - State of PA project 2019NF4100087335 (PI David Wolk, MD); Rooney Family Research Fund (PI David Wolk, MD); R01 AG055005 (PI David Wolk, MD); University of Pittsburgh - P50 AG005133 (PI Oscar Lopez MD); University of Southern California - P50 AG005142 (PI Helena Chui MD); University of Washington - P50 AG005136 (PI Thomas Grabowski MD); University of Wisconsin - P50 AG033514 (PI Sanjay Asthana MD FRCP); Vanderbilt University – P20 AG068082; Wake Forest - P30AG072947 (PI Suzanne Craft, PhD); Washington University, St. Louis - P01 AG03991 (PI John Morris MD); P01 AG026276 (PI John Morris MD); P20 MH071616 (PI Dan Marcus); P30 AG066444 (PI John Morris MD); P30 NS098577 (PI Dan Marcus); R01 AG021910 (PI Randy Buckner); R01 AG043434 (PI Catherine Roe); R01 EB009352 (PI Dan Marcus); UL1 TR000448 (PI Brad Evanoff); U24 RR021382 (PI Bruce Rosen); Avid Radiopharmaceuticals / Eli Lilly; Yale - P50 AG047270 (PI Stephen Strittmatter MD PhD); R01AG052560 (MPI: Christopher van Dyck, MD; Richard Carson, PhD); R01AG062276 (PI: Christopher van Dyck, MD); 1Florida - P30AG066506-03 (PI Glenn Smith, PhD); P50 AG047266 (PI Todd Golde MD PhD)

Data and/or research tools used in the preparation of this manuscript were obtained from the National Institute of Mental Health (NIMH) Data Archive (NDA). NDA is a collaborative informatics system created by the National Institutes of Health to provide a national resource to support and accelerate research in mental health. Dataset identifier(s): 2400, 2847, 2846, 2275, 2274, 2199, 2021, 1890, 19, 2936, 3142, 2803, 2607. This manuscript reflects the views of the authors and may not reflect the opinions or views of the NIH or of the Submitters submitting original data to NDA.

Data were provided in part by OASIS-3: Longitudinal Multimodal Neuroimaging: Principal Investigators: T. Benzinger, D. Marcus, J. Morris; NIH P30 AG066444, P50 AG00561, P30 NS09857781, P01 AG026276, P01 AG003991, R01 AG043434, UL1 TR000448, R01 EB009352. AV-45 doses were provided by Avid Radiopharmaceuticals, a wholly owned subsidiary of Eli Lilly.

Data used in the preparation of this article was obtained on 2023-09-23 from the Parkinson’s Progression Markers Initiative (PPMI) database (www.ppmi-info.org/access-dataspecimens/download-data; Tier 1), RRID:SCR 006431. For up-to-date information on the study, visit www.ppmi-info.org. PPMI – a public-private partnership – is funded by the Michael J. Fox Foundation for Parkinson’s Research, and funding partners; including including 4D Pharma, Abbvie, AcureX, Allergan, Amathus Therapeutics, Aligning Science Across Parkinson’s, AskBio, Avid Radiopharmaceuticals, BIAL, BioArctic, Biogen, Biohaven, BioLegend, BlueRock Therapeutics, Bristol-Myers Squibb, Calico Labs, Capsida Biotherapeutics, Celgene, Cerevel Therapeutics, Coave Therapeutics, DaCapo Brainscience, Denali, Edmond J. Safra Foundation, Eli Lilly, Gain Therapeutics, GE HealthCare, Genentech, GSK, Golub Capital, Handl Therapeutics, Insitro, Jazz Pharmaceuticals, Johnson Johnson Innovative Medicine, Lundbeck, Merck, Meso Scale Discovery, Mission Therapeutics, Neurocrine Biosciences, Neuron23, Neuropore, Pfizer, Piramal, Prevail Therapeutics, Roche, Sanofi, Servier, Sun Pharma Advanced Research Company, Takeda, Teva, UCB, Vanqua Bio, Verily, Voyager Therapeutics, the Weston Family Foundation and Yumanity Therapeutics.

The POND study was supported by the Ontario Brain Institute (grant number IDS-I 1-02).

The results published here are in whole or in part based on data obtained from the AD Knowledge Portal (https://adknowledgeportal.org). Study data were provided by the Rush Alzheimer’s Disease Center, Rush University Medical Center, Chicago. Data collection was supported through funding by NIA grants P30AG10161 (ROS), R01AG15819 (ROSMAP; genomics and RNAseq), R01AG17917 (MAP), R01AG30146, R01AG36042 (5hC methylation, ATACseq), RC2AG036547 (H3K9Ac), R01AG36836 (RNAseq), R01AG48015 (monocyte RNAseq) RF1AG57473 (single nucleus RNAseq), U01AG32984 (genomic and whole exome sequencing), U01AG46152 (ROSMAP AMP-AD, targeted proteomics), U01AG46161(TMT proteomics), U01AG61356 (whole genome sequencing, targeted proteomics, ROSMAP AMP-AD), the Illinois Department of Public Health (ROSMAP), and the Translational Genomics Research Institute (genomic). Additional phenotypic data can be requested at www.radc.rush.edu.

The content is the sole responsibility of the authors and does not necessarily represent official views of the NIA, NIH, or VA. The U.S. Department of Veterans Affairs, Department of Defense; National Personnel Records Center, National Archives and Records Administration; National Opinion Research Center; National Research Council, National Academy of Sciences; and the Institute for Survey Research, Temple University provided invaluable assistance in the creation of the VET Registry. The Cooperative Studies Program of the U.S. Department of Veterans Affairs provided financial support for development and maintenance of the Vietnam Era Twin Registry. We would also like to acknowledge the continued cooperation and participation of the members of the VET Registry and their families.

Data collection for the Vaghi datasets were supported by a Wellcome Trust Senior Investigator Award Grant No. 104631/Z/14/Z.

## References

[1] Abbie, Andrew Arthur (1940). “Cortical lamination in the monotremata”. In: Journal of Comparative Neurology 72.3, pp. 429–467.

[2] Alexander, Bonnie et al. (2017). “A new neonatal cortical and subcortical brain atlas: the Melbourne Children’s Regional Infant Brain (M-CRIB) atlas”. In: Neuroimage 147, pp. 841–851.

[3] Alexander-Bloch, Aaron, Jay N Giedd, and Ed Bullmore (2013). “Imaging structural co-variance between human brain regions”. In: Nature Reviews Neuroscience 14.5, pp. 322–336.

[4] Alexander-Bloch, Aaron F et al. (2018). “On testing for spatial correspondence between maps of human brain structure and function”. In: Neuroimage 178, pp. 540–551.

[5] Bedford, Saashi A, Jakob Seidlitz, and Richard AI Bethlehem (2022). “Translational potential of human brain charts”. In: Clinical and translational medicine 12.7, e960.

[6] Bellec, Pierre et al. (2017). “The neuro bureau ADHD-200 preprocessed repository”. In: Neuroimage 144, pp. 275–286.

[7] Bethlehem, Richard AI et al. (2022). “Brain charts for the human lifespan”. In: Nature 604.7906, pp. 525– 533.

[8] Bilder, R et al. (2020). “UCLA Consortium for Neuropsychiatric Phenomics LA5c Study”. OpenNeuro. doi: 10.18112/openneuro.ds000030.v1.0.0.

[9] Billot, Benjamin et al. (2023). “SynthSeg: Segmentation of brain MRI scans of any contrast and resolution without retraining”. In: Medical image analysis 86, p. 102789.

[10] Borghi, Elaine et al. (2006). “Construction of the World Health Organization child growth standards: selection of methods for attained growth curves”. In: Statistics in medicine 25.2, pp. 247–265.

[11] Botvinik-Nezer, Rotem et al. (2023). “Paingen?placebo”. OpenNeuro. doi: doi: 10.18112/openneuro.ds004746.v1.0.1.

[12] Bradley, Brian P (1980). “Developmental stability of Drosophila melanogaster under artificial and natural selection in constant and fluctuating environments”. In: Genetics 95.4, pp. 1033–1042.

[13] Bulik-Sullivan, Brendan et al. (2015). “An atlas of genetic correlations across human diseases and traits”. In: Nature genetics 47.11, pp. 1236–1241.

[14] Bulik-Sullivan, Brendan K et al. (2015). “LD Score regression distinguishes confounding from polygenicity in genome-wide association studies”. In: Nature genetics 47.3, pp. 291–295.

[15] Cheung, Moira S et al. (2024). “Growth reference charts for children with hypochondroplasia”. In: American Journal of Medical Genetics Part A 194.2, pp. 243– 252.

[16] Clarke, Geoffrey M and Leslie J McKenzie (1992). “Fluctuating asymmetry as a quality control indicator for insect mass rearing processes”. In: Journal of Economic Entomology 85.6, pp. 2045–2050.

[17] Corballis, Michael C (2009). “The evolution and genetics of cerebral asymmetry”. In: Philosophical Transactions of the Royal Society B: Biological Sciences 364.1519, pp. 867–879.

[18] Desikan, Rahul S et al. (2006). “An automated labeling system for subdividing the human cerebral cortex on MRI scans into gyral based regions of interest”. In: Neuroimage 31.3, pp. 968–980.

[19] Di Martino, Adriana et al. (2014). “The autism brain imaging data exchange: towards a large-scale evaluation of the intrinsic brain architecture in autism”. In: Molecular psychiatry 19.6, pp. 659–667.

[20] Esteves, M et al. (2021). “Asymmetrical brain plasticity: physiology and pathology”. In: Neuroscience 454, pp. 3–14.

[21] Fenton, Tanis R and Jae H Kim (2013). “A systematic review and meta-analysis to revise the Fenton growth chart for preterm infants”. In: BMC pediatrics 13, pp. 1–13.

[22] Fuchs, Bari et al. (2023). “Food and Brain Study”. OpenNeuro. doi: doi: 10.18112/openneuro.ds004697.v1.0.2.

[23] Garcini, Luz M et al. (2022). “Increasing diversity in developmental cognitive neuroscience: A roadmap for increasing representation in pediatric neuroimaging research”. In: Developmental Cognitive Neuroscience 58, p. 101167.

[24] Garza-Villarreal, Eduardo A. et al. (2021). “SUD-MEX?CONN: The Mexican dataset of cocaine use dis-order patients.”. OpenNeuro. doi: doi: 10.18112/openneuro.ds003346.v1.1.2.

[25] Hallgrímsson, Benedikt (1993). “Fluctuating asymmetry in Macaca fascicularis: A study of the etiology of developmental noise”. In: International Journal of Primatology 14, pp. 421–443.

[26] Hill, Jason, Donna Dierker, et al. (2010). “A surface-based analysis of hemispheric asymmetries and folding of cerebral cortex in term-born human infants”. In: Journal of neuroscience 30.6, pp. 2268–2276.

[27] Hill, Jason, Terrie Inder, et al. (2010). “Similar patterns of cortical expansion during human development and evolution”. In: Proceedings of the National Academy of Sciences 107.29, pp. 13135–13140.

[28] Holmes, Joni et al. (2019). “Protocol for a transdiagnostic study of children with problems of attention, learning and memory (CALM)”. In: BMC pediatrics 19, pp. 1–11.

[29] Jiang, Longda et al. (2019). “A resource-efficient tool for mixed model association analysis of large-scale data”. In: Nature genetics 51.12, pp. 1749–1755.

[30] Kang, Hyo Jung et al. (2011). “Spatio-temporal transcriptome of the human brain”. In: Nature 478.7370, pp. 483–489.

[31] Kang, Kaidi et al. (2024). “Study design features increase replicability in brain-wide association studies”. In: Nature, pp. 1–9.

[32] Kopal, Jakub, Lucina Q Uddin, and Danilo Bzdok (2023). “The end game: respecting major sources of population diversity”. In: Nature Methods 20.8, pp. 1122–1128.

[33] Korbmacher, Max et al. (2024). “Brain asymmetries from mid-to late life and hemispheric brain age”. In: Nature Communications 15.1, p. 956.

[34] Li, Gang et al. (2015). “Spatial patterns, longitudinal development, and hemispheric asymmetries of cortical thickness in infants from birth to 2 years of age”. In: Journal of neuroscience 35.24, pp. 9150–9162.

[35] Maingault, Sophie et al. (2016). “Regional correlations between cortical thickness and surface area asymmetries: A surface-based morphometry study of 250 adults”. In: Neuropsychologia 93, pp. 350–364.

[36] Makropoulos, Antonios et al. (2018). “The developing human connectome project: A minimal processing pipeline for neonatal cortical surface reconstruction”. In: Neuroimage 173, pp. 88–112.

[37] Mandal, Ayan S et al. (2024). “Normative trajectories of extra-axial cerebrospinal fluid during childhood and adolescence defined in a clinically-acquired MRI dataset”. In: medRxiv, pp. 2024–09.

[38] Marek, Scott and Timothy O Laumann (2025). “Replicability and generalizability in population psychiatric neuroimaging”. In: Neuropsychopharmacology 50.1, pp. 52–57.

[39] Nárai, Ádám et al. (2022). “Movement-related artefacts (MR-ART) dataset”. OpenNeuro. doi: doi:10.18112/openneuro.ds004173.v1.0.2.

[40] Nastase, Samuel A. et al. (2025). “Narratives”. OpenNeuro. doi: 10.18112/openneuro.ds002345.v1.1.4.

[41] Nielson, Dylan M. et al. (2023). “National Institute of Mental Health Characterization and Treatment of Adolescent Depression (NIMH CAT-D)”. OpenNeuro. doi: doi:10.18112/openneuro.ds004627.v1.0.0.

[42] Nugent, Allison C. et al. (2025). “The NIMH Healthy Research Volunteer Dataset”. OpenNeuro. doi: doi: 10.18112/openneuro.ds005752.v2.1.0.

[43] Ocklenburg, Sebastian et al. (2024). “Clinical implications of brain asymmetries”. In: Nature Reviews Neurology 20.7, pp. 383–394.

[44] Pandya, Deepak, Michael Petrides, and Patsy Benny Cipolloni (2015). Cerebral cortex: architecture, connections, and the dual origin concept. Oxford University Press.

[45] Panszczyk, Daniel et al. (2025). “Hemispheric asymmetry in language-related brain regions”. In: Brain Research 1857, p. 149606.

[46] Postema, Merel C, Martine Hoogman, et al. (2021). “Analysis of structural brain asymmetries in attention-deficit/hyperactivity disorder in 39 datasets”. In: Journal of Child Psychology and Psychiatry 62.10, pp. 1202–1219.

[47] Postema, Merel C, Daan Van Rooij, et al. (2019). “Altered structural brain asymmetry in autism spectrum disorder in a study of 54 datasets”. In: Nature communications 10.1, p. 4958.

[48] Raz, Naftali (2000). “Aging of the brain and its impact on cognitive performance: Integration of structural and functional findings.” In.

[49] Reynolds, Jess E et al. (2020). “Calgary Preschool magnetic resonance imaging (MRI) dataset”. In: Data in brief 29, p. 105224.

[50] Richardson, Hilary et al. (2019). “MRI data of 3–12 year old children and adults during viewing of a short animated film”. OpenNeuro.

[51] Roe, James M, Didac Vidal-Pineiro, et al. (2023). “Tracing the development and lifespan change of population-level structural asymmetry in the cerebral cortex”. In: Elife 12, e84685.

[52] Roe, James M, Didac Vidal-Piñeiro, et al. (2021). “Asymmetric thinning of the cerebral cortex across the adult lifespan is accelerated in Alzheimer’s disease”. In: Nature communications 12.1, p. 721.

[53] Rosen, Adon FG et al. (2018). “Quantitative assessment of structural image quality”. In: Neuroimage 169, pp. 407–418.

[54] Rutherford, Saige et al. (2022). “Charting brain growth and aging at high spatial precision”. In: elife 11, e72904.

[55] Saltoun, Karin et al. (2024). “Longitudinal changes in brain asymmetry track lifestyle and disease”. In: Research Square, rs–3.

[56] Schabdach, Jenna M et al. (2023). “Brain Growth Charts for Quantitative Analysis of Pediatric Clinical Brain MRI Scans with Limited Imaging Pathology”. In: Radiology 309.1, e230096.

[57] Schijven, Dick et al. (2023). “Large-scale analysis of structural brain asymmetries in schizophrenia via the ENIGMA consortium”. In: Proceedings of the National Academy of Sciences 120.14, e2213880120.

[58] Sha, Zhiqiang, Antonietta Pepe, et al. (2021). “Handedness and its genetic influences are associated with structural asymmetries of the cerebral cortex in 31,864 individuals”. In: Proceedings of the National Academy of Sciences 118.47, e2113095118.

[59] Sha, Zhiqiang, Dick Schijven, et al. (2021). “The genetic architecture of structural left–right asymmetry of the human brain”. In: Nature human behaviour 5.9, pp. 1226–1239.

[60] Sha, Zhiqiang, Daan Van Rooij, et al. (2022). “Subtly altered topological asymmetry of brain structural covariance networks in autism spectrum disorder across 43 datasets from the ENIGMA consortium”. In: Molecular psychiatry 27.4, pp. 2114–2125.

[61] Silva, Mariana da et al. (2025). “Differential patterns of cortical expansion in fetal and preterm brain development”. In: bioRxiv, pp. 2025–05.

[62] Snoek, Lukas et al. (2020a). “AOMIC-PIOP1”. OpenNeuro. doi: 10.18112/openneuro.ds002785.v2.0.0.

[63] Snoek, Lukas et al. (2020b). “AOMIC-PIOP2”. OpenNeuro. doi: 10.18112/openneuro.ds002790.v2.0.0.

[64] Snoek, Lukas et al. (2021). “AOMIC-ID1000”. OpenNeuro. doi: 10.18112/openneuro.ds003097.v1.2.1.

[65] Spreng, R. Nathan et al. (2022). “Neurocognitive aging data release with behavioral, structural, and multi-echo functional MRI measures”. OpenNeuro. doi: doi:10.18112/openneuro.ds003592.v1.0.9.

[66] Stasinopoulos, D Mikis and Robert A Rigby (2008). “Generalized additive models for location scale and shape (GAMLSS) in R”. In: Journal of Statistical Software 23, pp. 1–46.

[67] Sun, Lianglong et al. (2025). “Population-specific brain charts reveal Chinese-Western differences in neurodevelopmental trajectories”. In: bioRxiv, pp. 2025– 06.

[68] Tanaka, Saori C et al. (2021). “A multi-site, multi-disorder resting-state magnetic resonance image database”. In: Scientific data 8.1, p. 227.

[69] Tisdall, Loreen and Rui Mata (2023). “AgeRisk”. OpenNeuro. doi: doi: 10.18112/openneuro.ds004711.v1.0.0.

[70] Tomasi, Dardo and Nora D Volkow (2025). “Brain asymmetry and its association with inattention and heritability during neurodevelopment”. In: Translational Psychiatry 15.1, p. 96.

[71] Townsend, Grant C (1983). “Fluctuating dental asymmetry in Down’s syndrome”. In: Australian Dental Journal 28.1, pp. 39–44.

[72] Van Valen, Leigh (1962). “A study of fluctuating asymmetry”. In: Evolution, pp. 125–142.

[73] Váša, František, Carly Bennallick, et al. (2024). “Ultra-low-field brain MRI morphometry: test-retest reliability and correspondence to high-field MRI”. In: bioRxiv, pp. 2024–08.

[74] Váša, František, Jakob Seidlitz, et al. (2018). “Adolescent tuning of association cortex in human structural brain networks”. In: Cerebral Cortex 28.1, pp. 281–294.

[75] Wang, Jin et al. (2021). “A longitudinal neuroimaging dataset on language processing in children ages 5, 7, and 9 years old”. OpenNeuro. doi: 10.18112/openneuro.ds003604.v1.0.2.

[76] Warrier, Varun et al. (2023). “Genetic insights into human cortical organization and development through genome-wide analyses of 2,347 neuroimaging phenotypes”. In: Nature genetics 55.9, pp. 1483–1493.

[77] Webb, Jadon R et al. (2013). “Left-handedness among a community sample of psychiatric outpatients suffering from mood and psychotic disorders”. In: Sage Open 3.4, p. 2158244013503166.

[78] Wu, Xinran et al. (2025). “Developing brain asymmetry shapes cognitive and psychiatric outcomes in adolescence”. In: Nature Communications 16.1, pp. 1–15.

[79] Yang, Zhijian et al. (2024). “Brain aging patterns in a large and diverse cohort of 49,482 individuals”. In: Nature medicine 30.10, pp. 3015–3026.

[80] Yeatman, Jason D, Brian A Wandell, and Aviv A Mezer (2014). “Lifespan maturation and degeneration of human brain white matter”. In: Nature communications 5.1, p. 4932.

[81] Yu, Yuetong et al. (2024). “Brain-age prediction: Systematic evaluation of site effects, and sample age range and size”. In: Human Brain Mapping 45.10, e26768.

[82] Zareba, Michal Rafal et al. (2022). “Structural (t1) images of 136 young healthy adults; study of effects of chronotype, sleep quality and daytime sleepiness on brain structure.”. OpenNeuro. doi: doi:10.18112/openneuro.ds003826.v3.0.1.

[83] Zemel, Babette S et al. (2015). “Growth charts for children with Down syndrome in the United States”. In: Pediatrics 136.5, e1204–e1211.

