## Supplementary Information for "Charting structural brain asymmetry across the human lifespan"

[a]Lifespan Brain Institute, The Children's Hospital of Philadelphia and Penn Medicine, Philadelphia PA 19139, USA
[b]Department of Child and Adolescent Psychiatry and Behavioral Science, The Children's Hospital of Philadelphia, Philadelphia, PA, USA
[c]Department of Psychiatry, University of Pennsylvania, Philadelphia, PA, USA
[d]Department of Psychiatry, University of Cambridge, Cambridge, UK
[e]MRC Biostatistics Unit, University of Cambridge, Cambridge, UK
[f]Department of Psychology, University of Cambridge, Cambridge, UK
[g]Developmental Imaging, Murdoch Children's Research Institute, The Royal Children's Hospital Melbourne, Parkville, Victoria, Australia
[h]Department of Paediatrics, University of Melbourne, Melbourne, Victoria, Australia
[i]Department of Early Life Imaging, School of Biomedical Engineering and Imaging Sciences, King's College London, London, UK
[j]Medical Research Council Centre for Neurodevelopmental Disorders, King's College London, London, UK
[k]Department of Psychiatry, University of Cambridge, Cambridge, UK
[l]Department of Radiology, Children's Hospital of Philadelphia, Philadelphia, PA, United States
[m]Department of Radiology, Perelman School of Medicine, University of Pennsylvania, Philadelphia, PA, United States
[n]Research Department of Biomedical Computing, School of Biomedical Engineering and Imaging Sciences, King's College London, London, United Kingdom
[o]Clinical Neurosciences, University of Turku, Turku, Finland.
[p]FinnBrain Birth Cohort Study, Turku Brain and Mind Center, Department of Clinical Medicine, University of Turku and Turku University Hospital, Turku, Finland.
[q]Centre for Population Health Research, Turku University Hospital and University of Turku, Turku, Finland
[r]School of Psychological Sciences, Birkbeck University of London, London, UK
[s]Centre for Brain and Cognitive Development, Birkbeck College, University of London, London, UK
[t]Department of Clinical Sciences, SciLifeLab, Lund University, Lund, Sweden
[u]A full list of consortium authors can be found in the appendix
[v]Penn Statistics in Imaging and Visualization Center, Department of Biostatistics, Epidemiology, and Informatics, University of Pennsylvania
[w]Penn Lifespan Informatics and Neuroimaging Center (PennLINC), University of Pennsylvania, Philadelphia PA, USA
[x]Developmental Neuroscience, UCL Great Ormond Street Institute of Child Health, UCL, London, UK
[y]Department of Medical Physiology and Biophysics, University of Seville; Seville, Spain
[z]Instituto de Biomedicina de Sevilla (IBiS) HUVR/CSIC/University of Seville / CIBERSAM, ISCIII; Seville, Spain
[aa]Department of Neuroimaging, King's College London, London, UK
[ba]Section on Developmental Neurogenomics, Human Genetics Branch, National Institute of Mental Health, Bethesda, MD, 20892, USA
[*]Equal contributions
[**†] Corresponding authors

### Consortium member information

The members of the consortia used in this manuscript include:

##### Lifespan Brain Chart Consortium (LBCC)

Lena Dorfschmidt[1,2,3], Chris Adamson[4,5], Sophie Adler[6] [a], Evdokia Anagnostou[7,8] [b], Kevin M. Anderson[9], Annemieke M. Apergis-Schoute[10], Ariosky Areces-Gonzalez[11,12], Duncan E. Astle[13] [c], Bonnie Auyeung[14,15], Muhammad Ayub[16] [d], Jong Bin Bae[17], Gareth Ball[4,18] [e], Paula Banca[19,20], Simon Baron-Cohen[15,21], Richard Beare[4,5] [f], Saashi A. Bedford[15], Vivek Benegal[22] [g], Jonathan Berken[23,24], Richard A.I. Bethlehem[15,25], Frauke Beyer[26], Marjan Biria[h], John Blangero[29] [i], Manuel Blesa Cábez[30,31], James P. Boardman[30,31] [j], Matthew Borzage[32,33,34], Jorge F. Bosch-Bayard[35,36] [k], Niall Bourke, Matthew Buczek[23,24], Vince D. Calhoun[38] [l], Mallar M. Chakravarty[36,39] [m], Christina Chen[40], Casey Chertavian[3], Gaël Chetelat[41] [n], Yap S. Chong[42,43] [o], Aiden Corvin[44], Manuela Costantino[45,46], Eric Courchesne[47,48] [p], Fabrice Crivello[49], Vanessa L. Cropley[50], Jennifer Crosbie[51], Nicolas Crossley[52,53] [q], Kat Cyr[23,24], Anthony S. David[54], Marion Delarue[41], Richard Delorme[55,56], Sylvane Desrivieres, Gabriel A. Devenyi[58,59], Maria A. Di Biase[50,60], Ray Dolan[61,62,63] [r], Kirsten A. Donald[64] [s], Gary Donohoe, Katharine Dunlop[66,67] [t], Anthony D. Edwards[68,69,70] [u], Jed T. Elison[71] [v], Cameron T. Ellis[72,73], Jeremy A. Elman[74], Lisa Eyler[75,76], Damien A. Fair[71], Eric Feczko[77,78], Paul C. Fletcher[79,80] [w], Peter Fonagy[81,82] [x], Carol E. Franz[83] [y], Lidice Galan-Garcia[84], Margaret Gardner[23,24,85], Ali Gholipour[86] [z], Jay Giedd[87,88], John H. Gilmore[89], Juan Domingo Gispert López[90,91,92,93], David C. Glahn[94,95] [aa], Ian M. Goodyer[96], P. E. Grant[97], Nynke A. Groenewold[64], Faith M. Gunning[99], Raquel E. Gur[1,3], Ruben C. Gur[1,100], Christopher F. Hammill[51,101], Oskar Hansson[102,103], Trey Hedden[104,105] [bb], Andreas Heinz[106], Richard N. Henson[13,107] [cc], Katja Heuer[108], Jacqueline Hoare[109] [dd], Bharath Holla[110,111], Avram J. Holmes[112], Hao Huang[113] [ee], Kiho Im[114,115], Jonathan Ipser[116], Clifford R. Jack Jr[117], Andrea P. Jackowski[118,119], Tianye Jia[120,121,122], David T. Jones[123,124] [ff], Peter B. Jones[96,125] [gg], Benjamin Jung[23,24], Eren Kafadar[23,24,85], Rene S. Kahn[126,127], Shivaram Karandikar[23,24], Hasse Karlsson[128,129] [hh], Linnea Karlsson[129,130] [ii], Ryuta Kawashima[131], Elizabeth A. Kelley[132] [jj], Silke Kern[133,134] [kk], Ki-Woong Kim[135,136,137,138] [ll], Manfred G. Kitzbichler[96] [mm], William S. Kremen[74] [nn], François Lalonde[139], Brigitte Landeau[41], Jason Lerch[140,141,142] [oo], John D. Lewis[143], Jiao Li[144] [pp], Wei Liao[144] [qq], Conor Liston[145], Michael V. Lombardo[15,146], Pedro Luque Laguna[147], Jinglei Lv[50], Briana Macedo[23,24,85], Travis T. Mallard[149], Ayan Mandal[23,24,85], Machteld Marcelis[150], Samuel R. Mathias[94], Bernard Mazoyer[49,151], Philip McGuire[152], Michael J. Meaney[153], Andrea Mechelli[152], Laura Mercedes[23,24], Harri Merisaari[130], Kate Merritt[54], Bratislav Misic[154], Marcella Montagnese, Sarah E. Morgan[156,157,158], David Mothersill[159,161], Cynthia Ortinau[162] [rr], Rik Ossenkoppele[163,164], Minhui Ouyang[113], Eva Palacios Martínez[90], Lena Palaniyappan[165,166] [ss], Leo Paly[41], Pedro M. Pan[167,168] [tt], Christos Pantelis[169,170,171] [uu], Min Tae M. Park[172,173], Tomas Paus[174,175], Zdenka Pausova[51,176], Deirel Paz-Linares[11,177], Alexa Pichet Binette[178,179], Karen Pierce[180] [vv], Smrithi Prem[23,24,85], Elmo P. Pulli[130], Xing Qian[181], Anqi Qiu[182] [ww], Armin Raznahan[139], Timothy Rittman[183] [xx], Trevor W. Robbins[184], Amanda Rodrigue[94], Caitlin K. Rollins[185,186] [yy], Rafael Romero-Garcia[96,187] [zz], Lisa Ronan[96] [aaa], Monica D. Rosenberg[188] [bbb], David H. Rowitch[189] [ccc], Giovanni A. Salum[190,191,192], Theodore D. Satterthwaite[1] [ddd], H. Lina Schaare[194], Russell J. Schachar[51], Aaron P. Schultz[95,195,196], Michael Schöll[197,198,199] [eee], Jakob Seidlitz[1,2,3], Zhiqiang Sha[23,24,85], David Sharp[200], Russell T. Shinohara[40,202] [fff], Ingmar Skoog[133,134] [ggg], Christopher D. Smyser[203] [hhh], Reisa A. Sperling[95,195,204] [iii], Dan J. Stein, Aleks Stolicyn[206] [jjj], John Suckling[96,207], Gemma Sullivan[208], Kevin Sun[23,24,85], Benjamin Thyreau[131], Roberto Toro[108,209], Nicolas Traut[210,211], Kamen A. Tsvetanov[183,212] [kkk], Elise Turk[213,214], Nicholas B. Turk-Browne[9,215], Jetro J. Tuulari[129,130,216] [lll], Christophe Tzourio[217] [mmm], Étienne Vachon-Presseau[218,219,220], Mitchell J. Valdes-Sosa[84], Pedro A. Valdes-Sosa[222], Sofie L. Valk[223], Therese van Amelsvoort, Simon N. Vandekar[225,226] [nnn], Lana Vasung[227], Lindsay W. Victoria[99], Sylvia Villeneuve[178,179,228], Arno Villringer[26,229], Jacob W. Vogel[1] [ooo], Petra E. Vértes[96,230] [ppp], Konrad Wagstyl[231] [qqq], Yin-Shan S. Wang[232,233,234,235], Simon K. Warfield[86] [rrr], Varun Warrier[96], Eric Westman[236,237], Margaret L. Westwater[96], Heather C. Whalley[206,238], Simon R. White[96,239] [sss], Remo Williams[23,24], A. Veronica Witte[26,229,240] [ttt], Ning Yang[232,233,234,235], B.T. Thomas Yeo[241,244] [uuu], Hyuk Jin Yun[245], Andrew Zalesky[246] [vvv], Heather J. Zar[www], Anna Zettergren[133] [xxx], Juan H. Zhou[181,241,248] [yyy], Hisham Ziauddeen[96,249,250], Dabriel Zimmerman[1,2,3], Andre Zugman[168,251,252], Xi-Nian N. Zuo[232,233,234,253,254] [zzz], Edward T. Bullmore[96], Aaron F. Alexander-Bloch[1,2,3]

[1]Department of Psychiatry, University of Pennsylvania, Philadelphia, PA 19104 [2]Department of Child and Adolescent Psychiatry and Behavioral Science, The Children's Hospital of Philadelphia, Philadelphia, PA 19104 [3]Lifespan Brain Institute, The Children's Hospital of Philadelphia, Philadelphia, PA 19104 [4]Developmental Imaging, Murdoch Children's Research Institute, Melbourne, Victoria, Australia [5]Department of Medicine, Monash University, Melbourne, Victoria, Australia [6]UCL Great Ormond Street Institute for Child Health, 30 Guilford St, Holborn, London WC1N 1EH [7]Department of Pediatrics University of Toronto [8]Holland Bloorview Kids Rehabilitation Hospital, Toronto, Canada [9]Department of Psychology, Yale University, New Haven, CT, USA [10]School of Biological and Behavioural Sciences, Centre for Brain and Behaviour, Queen Mary University of London, London, United Kingdom [11]The Clinical Hospital of Chengdu Brain Science Institute, MOE Key Lab for NeuroInformation, University of Electronic Science and Technology of China, No. 2006, Xiyuan Ave., West Hi-Tech Zone, Chengdu, 611731, China [12]University of Pinar del Río "Hermanos Saiz Montes de Oca", Cuba [13]MRC Cognition and Brain Sciences Unit, Uni-

versity of Cambridge, Cambridge UK [14]Department of Psychology, School of Philosophy, Psychology and Language Sciences, University of Edinburgh, Edinburgh, United Kingdom [15]Autism Research Centre, Department of Psychiatry, University of Cambridge, Cambridge, CB2 0SZ, UK [16]University College London, Mental Health Neuroscience Research Department, Division of Psychiatry, London UK [17]Department of Neuropsychiatry, Seoul National University Bundang Hospital, Seongnam, Korea [18]Department of Paediatrics, University of Melbourne, Melbourne, Victoria, Australia [19]Department of Neuroscience, Faculty of Medicine and Nursing, University of the Basque Country, UPV/EHU, Vitoria-Gasteiz, Spain [20]IKERBASQUE, Basque Foundation for Science, Bilbao, Spain [21]Cambridge Lifetime Asperger Syndrome Service (CLASS), Cambridgeshire and Peterborough NHS Foundation Trust, Cambridge, United Kingdom [22]Centre for Addiction Medicine, National Institute of Mental Health and Neurosciences (NIMHANS), Bengaluru, India 560029 [23]Lifespan Brain Institute, The Children's Hospital of Philadelphia and Penn Medicine, Philadelphia PA 19139, USA [24]Department of Child and Adolescent Psychiatry and Behavioral Science, The Children's Hospital of Philadelphia, Philadelphia, PA, USA [25]Brain Mapping Unit, Department of Psychiatry, University of Cambridge, Cambridge, CB2 0SZ, UK [26]Department of Neurology, Max Planck Institute for Human Cognitive and Brain Sciences, Leipzig, 04103, Germany [27]The Institute of Psychiatry, Psychology & Neuroscience (IoPPN), King's College London, UK [28]Department of Psychology King's College London South London & Maudsley NHS Foundation Trust, UK [29]Department of Human Genetics, South Texas Diabetes and Obesity Institute, University of Texas Rio Grande Valley [30]Centre for Reproductive Health, Institute for Regeneration and Repair, University of Edinburgh, UK [31]Centre for Clinical Brain Sciences, University of Edinburgh, Edinburgh, UK [32]Fetal and Neonatal Institute, Division of Neonatology, Children's Hospital Los Angeles, Department of Pediatrics, Keck School of Medicine, University of Southern California, Los Angeles, California USA [33]Alfred E. Mann Department of Biomedical Engineering, Viterbi School of Engineering, University of Southern California, Los Angeles, USA [34]Department of Regulatory and Quality Sciences, Alfred E. Mann School of Pharmacy and Pharmaceutical Sciences, University of Southern California, Los Angeles, USA [35]McGill Centre for Integrative Neuroscience, Ludmer Centre for Neuroinformatics and Mental Health, Montreal Neurological Institute [36]McGill University [37]Department of Brain Sciences, Imperial College London, London UK & Care Research & Technology Centre, UK Dementia Research Institute [38]Tri-institutional Center for Translational Research in Neuroimaging and Data Science, Georgia State University, Georgia Institute of Technology, and Emory University, Atlanta, GA, USA [39]Computational Brain Anatomy (CoBrA) Laboratory, Cerebral Imaging Centre, Douglas Mental Health University Institute [40]Penn Statistics in Imaging and Visualization Center, Department of Biostatistics, Epidemiology, and Informatics, Perelman School of Medicine, University of Pennsylvania, Philadelphia, PA, USA [41]Normandie Univ, UNICAEN, INSERM, U1237, PhIND "Physiopathology and Imaging of Neurological Disorders", Institut Blood and Brain @ Caen-Normandie, Cyceron, 14000 Caen, France [42]Singapore Institute for Clinical Sciences, Agency for Science, Technology and Research, Singapore [43]Department of Obstetrics and Gynaecology, Yong Loo Lin School of Medicine, National University of Singapore, Singapore [44]Department of Psychiatry, Trinity College, Dublin, Ireland [45]Cerebral Imaging Centre, Douglas Mental Health University Institute, Verdun, Canada [46]Undergraduate program in Neuroscience, McGill University, Montreal, Canada [47]Department of Neuroscience, University of California, San Diego, San Diego, CA 92093, USA [48]Autism Center of Excellence, University of California, San Diego, San Diego, CA 92037, USA [49]Institute of Neurodegenerative Disorders, CNRS UMR5293, CEA, University of Bordeaux [50]Melbourne Neuropsychiatry Centre, University of Melbourne, Melbourne, Australia [51]The Hospital for Sick Children, Toronto, Canada [52]Department of Psychiatry, School of Medicine, Pontificia Universidad Católica de Chile, Diagonal Paraguay 362, Santiago 8330077, Chile [53]Department of Psychiatry, University of Oxford OX3 7JX [54]Division of Psychiatry, University College London, London, UK [55]Child and Adolescent Psychiatry Department, Robert Debré University Hospital, AP-HP, F-75019, Paris France [56]Human Genetics and Cognitive Functions , Institut Pasteur, F-75015, Paris France [57]Social, Genetic and Developmental Psychiatry Centre, Institute of Psychiatry, Psychology & Neuroscience, King's College London, London, United Kingdom [58]Cerebral Imaging Centre, Douglas Mental Health University Institute, Montreal, QC, Canada, McGill Department of Psychiatry, Montreal, QC, Canada [59]Department of Psychiatry, McGill University, Montreal, QC, Canada [60]Department of Psychiatry, Brigham and Women's Hospital, Harvard Medical School, Boston, Massachusetts, United States [61]Max Planck UCL Centre for Computational Psychiatry and Ageing Research, University College London, London, UK [62]Wellcome Centre for Human Neuroimaging, University College London, London, UK [63]Wellcome Centre for Human Neuroimaging, 12 Queen Square, London WC1N 3AR [64]Neuroscience Institute, University of Cape Town, Cape Town, South Africa [65]Center for Neuroimaging, Cognition & Genomics (NICOG), School of Psychology, National University of Ireland Galway, Galway, Ireland [66]Department of Psychiatry, University of Toronto, Toronto, Ontario, Canada [67]Centre for Depression and Suicide Studies, Unity Health Network, Toronto, Ontario, Canada [68]Centre for the Developing Brain, King's College London, London, UK [69]Evelina London Children's Hospital [70]MRC Centre for Neurodevelopmental Disorders, London [71]Institute of Child Development, Department of Pediatrics, Masonic Institute for the Developing Brain, University of Minnesota, Minneapolis, MN, United States [72]Department of Psychology, Stanford University, Stanford, CA, USA [73]Haskins Laboratories, New Haven, CT, USA [74]Department of Psychiatry, Center for Behavior Genetics of Aging, University of California, San Diego, La Jolla, CA [75]Desert-Pacific Mental Illness Research Education and Clinical Center, VA San Diego Healthcare, San Diego, CA, USA [76]Department of Psychiatry, University of California San Diego, Los Angeles, CA, USA [77]Masonic Institute for the Developing Brain, University of Minnesota, Minneapolis, Minnesota [78]Department of Pediatrics, University of Minnesota, Minneapolis, Minnesota [79]Department of Psychiatry, University of Cambridge, and Wellcome Trust MRC Institute of Metabolic Science, Cambridge Biomedical Campus, Cambridge, United Kingdom [80]Cambridgeshire and Peterborough NHS Foundation Trust [81]Department of Clinical, Educational and Health Psychology, University College London, London, UK [82]Anna Freud National Centre for Children and Families, London UK [83]Department of Psychiatry, Center for Behavior Genetics of Aging, University of California, San Diego, La Jolla, CA 92093 [84]Cuban Center for Neuroscience, La Habana, Cuba [85]Department of Psychiatry, University of Pennsylvania, Philadelphia, PA, USA [86]Computational Radiology Laboratory, Boston Children's Hospital, Boston, MA 02115 [87]Department of Child and Adolescent Psychiatry, University of California, San Diego, San Diego, CA 92093, USA [88]Department of Psychiatry, University of California San Diego, San Diego, CA, USA [89]Department of Psychiatry, University of North Carolina, Chapel Hill, NC, USA [90]Barcelona$\beta$eta Brain Research Center (BBRC), Pasqual Maragall Foundation, Barcelona, Spain [91]Hospital del Mar Research Institute, Barcelona, Spain [92]Centro de Investigación Biomédica en Red Bioingeniería, Biomateriales y Nanomedicina, Instituto de Salud Carlos III, Madrid, Spain [93]Centro Nacional de Investigaciones Cardiovasculares (CNIC), Madrid, Spain [94]Department of Psychiatry, Boston Children's Hospital and Harvard Medical School, Boston, MA 02115 [95]Harvard Medical School, Boston, MA 02115 [96]Department of Psychiatry, University of Cambridge, Cambridge, CB2 0SZ, UK [97]Division of Newborn Medicine and Neuroradiology, Fetal Neonatal Neuroimaging and Developmental Science Center, Boston Children's Hospital, Harvard Medical School, Boston, MA 02115, USA [98]Department of Paediatrics and Child Health, Red Cross War Memorial Children's Hospital, SA-MRC Unit on Child & Adolescent Health, University of Cape Town, South Africa [99]Weill Cornell Institute of Geriatric Psychiatry, Department of Psychiatry, Weill Cornell Medicine [100]Lifespan Brain Institute, The Children's Hospital of Philadelphia, Philadelphia, PA 19105 [101]Mouse Imaging Centre, Toronto, Canada [102]Clinical Memory Research Unit, Department of Clinical Sciences Malmö, Lund University, Malmö, Sweden [103]Memory Clinic, Skåne University Hospital, Malmö, Sweden [104]Department of Neurology, Icahn School of Medicine at Mount Sinai, New York, NY 10029, USA [105]Athinoula A. Martinos Center for Biomedical Imaging, Department of Radiology, Massachusetts General Hospital, Harvard Medical School, Boston, MA 02129, USA [106]Department of Psychiatry and Psychotherapy, Charite University Hospital Berlin, Berlin, Germany [107]Department of Psychiatry, University of Cambridge, Cambridge, UK [108]Institut Pasteur, Université Paris Cité, Unité de Neuroanatomie Appliquée et Théorique, F-75015 Paris, France [109]Department of Psychiatry, University of Cape Town, Cape Town, South Africa [110]Department of Integrative Medicine, NIMHANS, Bengaluru-560029, India [111]Accelerator Program for Discovery in Brain disorders using Stem cells (ADBS), Department of Psychiatry, NIMHANS, Bengaluru-560029, India [112]Department of Psychiatry, Brain Health Institute, Rutgers University, Piscataway, NJ, USA [113]Department of Radiology, Children's Hospital of Philadelphia and University of Pennsylvania, Philadelphia, PA 19104 [114]Division of Newborn Medicine, Fetal Neonatal Neuroimaging and Developmental Science Center, Boston Children's Hospital, Harvard Medical School, Boston, MA 02115, USA [115]Boston Children's Hospital, Boston, MA 02115 [116]Department of Psychiatry and Mental Health, Clinical Neuroscience Institute, University of Cape Town [117]Department of Radiology, Mayo Clinic, Rochester, MN 55905, USA [118]Department of Psychiatry, Universidade Federal de São Paulo [119]National Institute of Developmental Psychiatry, CNPq [120]Institute of Science and Technology for Brain-Inspired Intelligence, Fudan University, Shanghai, 200433, China [121]Key Laboratory of Computational Neuroscience and BrainInspired Intelligence (Fudan University), Ministry of Education, Shanghai, China [122]Centre for Population Neuroscience and Precision Medicine (PONS), Institute of Psychiatry, Psychology and Neuroscience, SGDP Centre, King's College London, London SE5 8AF, UK [123]Department of Neurology, Mayo Clinic, Rochester, MN, USA [124]Department of Radiology, Mayo Clinic, Rochester, MN, USA [125]Cambridgeshire and Peterborough NHS Foundation Trust, Huntingdon, United Kingdom [126]Department of Psychiatry, Icahn School of Medicine at Mount Sinai, New York, NY, USA [127]Department of Psychiatry, Icahn School of Medicine, Mount Sinai, New York, USA [128]Department of Clinical Medicine, Department of Psychiatry and Turku Brain and Mind Center, FinnBrain Birth Cohort Study, University of Turku and Turku University Hospital, Turku, Finland [129]Centre for Population Health Research, Turku University Hospital and University of Turku, Turku, Finland [130]FinnBrain Birth Cohort Study, Turku Brain and Mind Center, Department of Clinical Medicine, University of Turku and Turku University Hospital, Turku, Finland [131]Institute of Development, Aging and Cancer, Tohoku University, Seiryocho, Aobaku, Sendai 980-8575, Japan [132]Queen's University, Departments of Psychology and Psychiatry, Centre for Neuroscience Studies, Kingston, Ontario, Canada [133]Neuropsychiatric Epidemiology Unit, Department of Psychiatry and Neurochemistry, Institute of Neuroscience and Physiology, the Sahlgrenska Academy, Centre for Ageing and Health (AGECAP) at the University of Gothenburg, Sweden [134]Region Västra Götaland, Sahlgrenska University Hospital, Psychiatry, Cognition and Old Age Psychiatry Clinic, Gothenburg, Sweden [135]Department of Brain and Cognitive Sciences, Seoul National University College of Natural Sciences, Seoul, Republic of Korea [136]Department of Neuropsychiatry, Seoul National University Bundang Hospital, Seongnam, Republic of Korea [137]Department of Psychiatry, Seoul National University College of Medicine, Seoul, Republic of Korea [138]Department of Brain and Cognitive Science, Seoul National University College of Natural Sciences [139]Section on Developmental Neurogenomics, Human Genetics Branch, National Institute of Mental Health, Bethesda, MD, USA [140]Department of Medical Biophysics, University of Toronto, Toronto, ON, Canada [141]Mouse Imaging Centre, The Hospital for Sick Children, Toronto, ON, Canada [142]Wellcome Centre for Integrative Neuroimaging, FMRIB, Nuffield Department of Clinical Neuroscience, University of Oxford, Oxford, UK [143]Montreal Neurological Institute, McGill University, Montreal, Canada [144]The Clinical Hospital of Chengdu Brain Science Institute, University of Electronic Science and Technology of China, Chengdu 611731, China [145]Department of Psychiatry and Brain and

Mind Research Institute, Weill Cornell Medicine [146]Laboratory for Autism and Neurodevelopmental Disorders, Center for Neuroscience and Cognitive Systems @UniTn, Istituto Italiano di Tecnologia, Rovereto, Italy [147]The Cardiff University Brain Research Imaging Centre (CUBRIC), Cardiff University, Cardiff, UK [148]School of Biomedical Engineering & Brain and Mind Centre, The University of Sydney, Sydney, NSW, Australia [149]Department of Psychology, University of Texas, Austin, Texas 78712, USA [150]Department of Psychiatry and Neuropsychology, School of Mental Health and Neuroscience, EURON, Maastricht University Medical Centre, PO Box 616, 6200 MD, Maastricht, the Netherlands; Institute for Mental Health Care Eindhoven (GGzE), Eindhoven, the Netherlands [151]Bordeaux University Hospital [152]Professor, Department of Psychosis Studies, Institute of Psychiatry, Psychology and Neuroscience, King's College London, UK [153]Ludmer Centre for Neuroinformatics and Mental Health, Douglas Mental Health University Institute, McGill University, Montreal, Quebec, Canada; Singapore Institute for Clinical Sciences, Singapore [154]McConnell Brain Imaging Centre, Montreal Neurological Institute, McGill University, Montreal, QC H3A 2B4, Canada [155]Department of Neuroimaging, Institute of Psychiatry, Psychology & Neuroscience, King's College London, London, United Kingdom [156]School of Biomedical Engineering and Imaging Sciences, King's College London, London UK [157]Department of Computer Science and Technology, University of Cambridge, Cambridge CB3 0FD, United Kingdom [158]Department of Psychiatry, University of Cambridge, Cambridge CB2 0SZ, United Kingdom [159]Department of Psychology, School of Business, National College of Ireland, Dublin, Ireland [160]School of Psychology & Center for Neuroimaging and Cognitive Genomics, National University of Ireland Galway, Galway, Ireland [161]Department of Psychiatry, Trinity College Dublin, Dublin, Ireland [162]Department of Pediatrics, Washington University in St. Louis, St. Louis, Missouri, United States [163]Alzheimer Center Amsterdam, Department of Neurology, Amsterdam Neuroscience, Vrije Universiteit Amsterdam, Amsterdam UMC, Amsterdam, The Netherlands [164]Lund University, Clinical Memory Research Unit, Lund, Sweden [165]Douglas Mental Health University Institute, Department of Psychiatry, McGill University, Quebec, Canada [166]Robarts Research Institute, University of Western Ontario, London, Ontario, Canada [167]Department of Psychiatry, Universidade Federal de São Paulo, Brazil [168]National Institute of Developmental Psychiatry for Children and Adolescents (INPD), Brazil [169]Melbourne Neuropsychiatry Centre, Department of Psychiatry, The University of Melbourne and Melbourne Health, Carlton South, Victoria, Australia [170]Melbourne School of Engineering, The University of Melbourne, Parkville, Victoria, Australia [171]Florey Institute of Neuroscience and Mental Health, Parkville, VIC, Australia [172]Cerebral Imaging Centre, Douglas Mental Health University Institute, Montreal, Canada [173]Integrated Program in Neuroscience, McGill University, Montreal, Canada [174]Department of Psychiatry, Faculty of Medicine and Centre Hospitalier Universitaire Sainte-Justine, University of Montreal, Montreal, Quebec, Canada [175]Departments of Psychiatry and Psychology, University of Toronto, Toronto, ON, Canada [176]Departments of Physiology and Nutritional Sciences, University of Toronto, Toronto, Canada [177]Cuban Neuroscience Center, Havana, Cuba [178]Department of Psychiatry, Faculty of Medicine, McGill University, Montreal, Qc, H3A 1Y2, Canada [179]Douglas Mental Health University Institute, Montreal, Qc, H4H 1R3, Canada [180]Department of Neurosciences, University of California, San Diego La Jolla, CA, USA [181]Center for Sleep and Cognition, Yong Loo Lin School of Medicine, National University of Singapore, Singapore [182]Department of Health Technology and Informatics, Mental Health Research Centre, The Hong Kong Polytechnic University [183]Department of Clinical Neurosciences, University of Cambridge, Cambridge UK [184]Department of Psychology and the Behavioural and Clinical Neuroscience Institute, University of Cambridge, Cambridge, CB2 3EB, UK [185]Department of Neurology, Harvard Medical School [186]Department of Neurology, Boston Children's Hospital, Boston, MA 02115 [187]Instituto de Biomedicina de Sevilla (IBiS) HUVR/CSIC/Universidad de Sevilla, Dpto. de Fisiología Médica y Biofísica, Spain [188]Department of Psychology, Neuroscience Institute, University of Chicago [189]Department of Paediatrics and Wellcome-MRC Cambridge Stem Cell Institute, University of Cambridge, Hills Road, Cambridge, UK [190]Department of Psychiatry, Universidade Federal do Rio Grande do Sul (UFRGS) [191]National Institute of Developmental Psychiatry (INPD) [192]Child Mind Institute, New York, USA [193]Lifespan Informatics & Neuroimaging Center, University of Pennsylvania, Philadelphia, PA 19104 [194]U Bremen Research Alliance, Bremen, Germany [195]Harvard Aging Brain Study, Department of Neurology, Massachusetts General Hospital, Boston, MA 02114 [196]Athinoula A. Martinos Center for Biomedical Imaging, Department of Radiology, Massachusetts General Hospital, Charlestown, MA 02129, USA [197]Wallenberg Centre for Molecular and Translational Medicine, University of Gothenburg, Gothenburg, Sweden [198]Department of Psychiatry and Neurochemistry, University of Gothenburg, Sweden [199]Dementia Research Centre, Queen's Square Institute of Neurology, University College London, UK [200]Department of Brain Sciences, Imperial College London, London UK [201]Care Research & Technology Centre, UK Dementia Research Institute [202]Center For AI And Data Science For Integrated Diagnostics, Department of Radiology, Perelman School of Medicine, University of Pennsylvania, Philadelphia, PA, USA [203]Departments of Neurology, Pediatrics, and Radiology, Washington University School of Medicine, St. Louis, United States [204]Center for Alzheimer Research and Treatment, Department of Neurology, Brigham and Women's Hospital, Boston, MA 02115 [205]SA MRC Unit on Risk & Resilience in Mental Disorders, Dept of Psychiatry and Neuroscience Institute, University of Cape Town, Cape Town, South Africa [206]Division of Psychiatry, Centre for Clinical Brain Sciences, University of Edinburgh, UK [207]Cambridge and Peterborough Foundation NHS Trust [208]MRC Centre for Reproductive Health, University of Edinburgh, UK [209]Université de Paris, Paris, France [210]Department of Neuroscience, Institut Pasteur, Paris, France [211]Center for Research and Interdisciplinarity (CRI), Université Paris Descartes, Paris, France [212]Department of Psychology, University of Cambridge, Cambridge, UK [213]Department of Cognitive Neuropsychology, Tilburg University, Warandelaan 2, 5000 LE, Tilburg, the Netherlands [214]Department of Neonatology, University Medical Center Utrecht, Utrecht University Heidelberglaan 100, 3584 CX, Utrecht, the Netherlands [215]Wu Tsai Institute, Yale University, New Haven, CT, USA [216]Neurocenter, Turku University Hospital, Turku, Finland [217]Univ. Bordeaux, Inserm, Bordeaux Population Health Research Center, U1219, CHU Bordeaux, F-33000 Bordeaux, France [218]Faculty of Dental Medicine and Oral Health Sciences, McGill University, Montreal, Qc, H3A 1G1, Canada [219]Faculty of Dentistry, McGill University, Montreal, Qc, H3A 1G1, Canada [220]Alan Edwards Centre for Research on Pain (AECRP), McGill University, Montreal, Qc, H3A 1G1, Canada [221]Brain-Computer Interface & Brain-Inspired Intelligence Key Laboratory of Sichuan Province, Chengdu, Sichuan, China [222]University of Electronic Science and Technology of China/Cuban Center for Neuroscience [223]Institute for Neuroscience and Medicine 7, Forschungszentrum Juelich; Max Planck Institute for Human Cognitive and Brain Sciences [224]Department of Psychiatry & Neurosychology, Maastricht University, Maastricht, The Netherlands [225]Department of Biostatistics, Vanderbilt University, Nashville, Tennessee, USA [226]Department of Biostatistics, Vanderbilt University Medical Center, Nashville, Tennessee, USA [227]Division of Newborn Medicine, Fetal Neonatal Neuroimaging and Developmental Science Center, Department of Pediatrics, Boston Children's Hospital, Boston, MA 02115 [228]McConnell Brain Imaging Center, Montreal Neurological Institute, McGill University, Montreal, Quebec, Canada [229]Clinic for Cognitive Neurology, University of Leipzig Medical Center, Leipzig, 04103, Germany [230]The Alan Turing Institute, London NW1 2DB, UK [231]Wellcome Centre for Human Neuroimaging, Institute of Neurology, University College London, WC1N 3AR [232]State Key Laboratory of Cognitive Neuroscience and Learning, Beijing Normal University, Beijing 100875, China [233]Developmental Population Neuroscience Research Center, IDG/McGovern Institute for Brain Research, Beijing Normal University, Beijing 100875, China [234]National Basic Science Data Center, Beijing 100190, China [235]Research Center for Lifespan Development of Brain and Mind, Institute of Psychology, Chinese Academy of Sciences, Beijing 100101, China [236]Division of Clinical Geriatrics, Center for Alzheimer Research, Department of Neurobiology, Care Sciences and Society, Karolinska Institutet, Stockholm, Sweden [237]Ageing Epidemiology Research Unit, School of Public Health, Imperial College London, UK [238]Generation Scotland, University of Edinburgh [239]MRC Biostatistics Unit, University of Cambridge, Cambridge, England [240]Faculty of Medicine, CRC 1052 'Obesity Mechanisms', University of Leipzig, Leipzig, 04103, Germany [241]Department of Electrical and Computer Engineering, National University of Singapore, Singapore [242]Centre for Sleep & Cognition and Centre for Translational MR Research, Yong Loo Lin School of Medicine, National University of Singapore, Singapore [243]N.1 Institute for Health & Institute for Digital Medicine, National University of Singapore, Singapore [244]Integrative Sciences and Engineering Programme (ISEP), National University of Singapore, Singapore [245]Fetal Neonatal Neuroimaging and Developmental Science Center, Division of Newborn Medicine, Boston Children's Hospital, Harvard Medical School, Boston, MA 02115, USA [246]Melbourne Neuropsychiatry Centre, University of Melbourne, Melbourne, Australia; Department of Biomedical Engineering, University of Melbourne, Melbourne, Australia [247]SAMRC Unit on Child & Adolescent Health, University of Cape Town, South Africa [248]Center for Translational Magnetic Resonance Research, Yong Loo Lin School of Medicine, National University of Singapore, Singapore [249]Wellcome Trust-MRC Institute of Metabolic Science, University of Cambridge, Cambridge, CB2 0SZ [250]Cambridgeshire and Peterborough Foundation Trust, Cambridge, CB21 5EF [251]National Institute of Mental Health (NIMH), National Institutes of Health (NIH), Bethesda, Maryland, USA [252]Department of Psychiatry, Escola Paulista de Medicina, São Paulo, Brazil [253]Research Center for Lifespan Development of Brain and Mind, Institute of Psychology, Chinese Academy of Sciences, Beijing 100101, China [254] Key Laboratory of Brain and Education, School of Education Science, Nanning Normal University, Nanning 530001, China

[a]S.A was supported by the Rosetrees Trust (A2665) [b]E.A and POND collection is funded by the Ontario Brain Institute [c]D.E.A. was supported by MRC Programme Grant MC-A0606-5PQ41 [d]M.A. is supported by NIH 1R01MH112904-01 [e]G.B. was supported by an NHMRC Investigator Grant (1194497) and the Royal Children's Hospital Foundation. [f]R.B was supported by a Royal Children's Hospital Foundation Grant [g]cVEDA is jointly funded by the Indian Council for Medical Research (ICMR/MRC/3/M/2015-NCD-I) and the Newton Grant from the Medical Research Council(MR/N000390/1), United Kingdom. [h]MB was supported by the MHRUK and Angharad-Dodds Bursaries [i]The Genetics of Brain Structure and Function project was funnded by the NIH (MH078143, MH083824, and MH078111) [j]TEBC data collection was funded by Theirworld [k]JBB was supported by Brain Canada (243030), the Fonds de recherche du Québec (FRQ) Healthy Brains Healthy Liles (HBHL) FRQ/Canada-Cuba-China Axis (246117), the Canada First Research Excellence Fund (CFREF)/HBHL BigBrain Analytics and Learning Laboratory (HIBALL), and Helmholtz (252428) [l]VDC was suppored by NSF 2112455 and NIH R01MH123610 [m]MMC is supported by Fonds Recherche [n]IMAP study (PI (scientific): G Chetelat; PI (MD) V de La Sayette)) was funded by Programme Hospitalier de Recherche Clinique (PHRCN 2011-A01493-38 and PHRCN 2012 12-006-0347) and Agence Nationale de la Recherche (LONGVIE 2007). Dr Chetelat's research including IMAP was also funded by Institut National de la Santé et de la Recherche Médicale (Inserm), Fondation Plan Alzheimer (Alzheimer Plan 2008-2012); Région Basse-Normandie; Association France Alzheimer et maladies apparentées, Fondation Vaincre Alzheimer. [o]GUSTO study is supported by the Singapore National Research Foundation under its Translational and Clinical Research (TCR) Flagship Programme and administered by the Singapore Ministry of Health's National Medical Research Council (NMRC), Singapore - NMRC/TCR/004-NUS/2008; NMRC/TCR/012-NUHS/2014. Additional funding is provided by the Singapore Institute for Clinical Sciences, Agency for Science Technology and Research (A*STAR), Singapore [p]E.C. is supported by grants: NIMH P50-MH081755, NIMH R01-MH036840, NIMH R01-MH110558, NIMH U01-

MH108898, NIDCD R01-DC016385 [q]NC is supported by the Agencia Nacional de Investigación y Desarrollo (ANID) Chile through grants FONDECYT regular 1200601, ANILLO PIA ACT192064 [r]RJD is supported by the Max Planck Society (MPS) [s]The Drakenstein Child Health Study is funded by the Bill and Melinda Gates Foundation (OPP 1017641). Additional support for HJZ and DJS was provided by the Medical Research Council of South Africa. KAD and aspects of the research are additionally supported by the NRF, an Academy of Medical Sciences Newton Advanced Fellowship (NAF002/1001) funded by the UK Government's Newton Fund, by NIAAA via (R21AA023887), by the Collaborative Initiative on Fetal Alcohol Spectrum Disorders (CIFASD) developmental grant (U24 AA014811), and by the US Brain and Behaviour Foundation Independent Investigator grant (24467) [t]KD is supported by a Canadian Institutes for Health Research Banting Postdoctoral Fellowship. [u]The Developing Human Connectome Project was supported by the European Research Council under the European Union Seventh Framework Programme (FP/2007-2013)/ERC Grant Agreement No. 319456 [v]The BCP was supported by grants: U01MH110274 & R01MH104324 [w]PCF is supported by the Wellcome Trust (Reference No. Reference No. 206368/Z/17/Z) and by the Bernard Wolfe health Neuroscience Fund [x]PF is supported by edical Research Council (MRC) (reference MR/V049941/1) and National Institute for Health Research (NIHR) (reference NIHR131339) [y]C.E.F. is supported by grants: R01s AG050595, AG022381, AG037985; & P01 AG055367; The content is the sole responsibility of the authors and does not necessarily represent official views of the NIA, NIH, or VA. The U.S. Department of Veterans Affairs, Department of Defense; National Personnel Records Center, National Archives and Records Administration; National Opinion Research Center; National Research Council, National Academy of Sciences; and the Institute for Survey Research, Temple University provided invaluable assistance in the creation of the VET Registry. The Cooperative Studies Program of the U.S. Department of Veterans Affairs provided financial support for development and maintenance of the Vietnam Era Twin Registry. We would also like to acknowledge the continued cooperation and participation of the members of the VET Registry and their families. [z]A.G. is supported by grants: NIH R01 EB018988, R01 NS106030, and R01 EB031849 [aa]The Genetics of Brain Structure and Function project was funded by the NIH (MH078143, MH083824, and MH078111) [bb]Contributed data were collected with support from P01AG036694 and R01AG053509. T.H. is supported by R01AG053509 and P30AG066514. [cc]RNH is supported by MRC Programme Grant SUAG/046 G101400 [dd]R01HD074051 [ee]H.H is funded by NIH R01MH092535, R01EB031285, and R01MH125333. [ff]U01 AG06786 [gg]PBJ was funded by the Wellcome Trust (095844/Z/11/Z) and NIHR RP-PG-0616-20003 [hh]FinnBrain was funded by Jane and Aatos Erkko Foundation, Signe and Ane Gyllenberg Foundation [ii]LK was funded by the Brain and Behavior Research Foundation, NARSAD YI Grant 1956 and the Academy of Finland Profi 5 325292 [jj]E.A.K. is funded by the Masonic Foundation of Ontario, the National Institute of Mental Health, and the Ontario Brain Institute [kk]SK was financed by grants from the Swedish state under the agreement between the Swedish government and the county councils, the ALF-agreement ( ALFGBG-965923,ALFGBG-81392, ALF GBG-771071). The Alzheimerfonden (AF-842471, AF-737641, AF-939825 ). The Swedish Research Council (2019-02075). SK has served at scientific advisory boards and / or as consultant for Geras Solutions and Biogen. [ll]The KNE96 study was supported by a grant of the Korean Health Technology R&D Project, Ministry for Health, Welfare, and Family Affairs, Republic of Korea (grant number I09C1379(A092077) [mm]The BIODEP study was sponsored by the Cambridgeshire and Peterborough NHS Foundation Trust and the University of Cambridge, and funded by a strategic award from the Wellcome Trust (104025) in partnership with Janssen, GlaxoSmithKline, Lundbeck and Pfizer. [nn]W.S.K. was supported by NIA grants R01s AG050595, AG022381, AG037985, and P01 AG055367. [oo]The POND study was supported by the Ontario Brain Institute (grant number IDS-I 1-02). This organization did not play a role in thedesign of the study, the collection, analysis and interpretation ofthe data, and in writing the manuscript. [pp]J.L. was supported by the China Postdoctoral Science Foundation (BX2021057). [qq]W.L. was supported by the National Natural Science Foundation of China (61871077). [rr]Supported by the NIH NHLBI K23HL141602 (C.M.O) and the Mend A Heart Foundation (C.M.O) [ss]This study was funded by CIHR Foundation Grant (375104/2017); Schulich School of Medicine Clinical Investigator Fellowship; Bucke Family Fund; Grad student salary support by NSERC Discovery Grant (No. RGPIN2016-05055); Canada Graduate Scholarship. Data acquisition was supported by the Canada First Excellence Research Fund to BrainSCAN, Western University (Imaging Core); Compute Canada Resources were used in the storage of imaging data. L. Palaniyappan's research is supported by the Monique H. Bourgeois Chair in Developmental Disorders and the Graham Boeckh Foundation. He receives a salary award from the Fonds de recherche du Québec-Santé (FRQS 366934). Data acquisition supported by Drs. Khan, Gati, and the research staff at the CFMM, Robarts Imaging and the clinical staff at the PEPP Clinic, London, Ontario. This study was conducted according to the approval by the Research Ethics Board of the University of Western Ontario (Project 108268; October 19, 2020). [tt]The study was funded by Conselho Nacional de Desenvolvimento Científico e Tecnológico (CNPq grant numbers 573974/2008-0 and 465550/2014-2), Fundação de Amparo à Pesquisa do Estado de São Paulo (FAPESP grant numbers: 2008/57896-8, 2013/08531-5, 2014/50917-0, 2020/06172-1, 2021/05332-8, 2021/12901-9), European Research Council (ERC grant numbers: 337673 and 101057390), UK Medical Research Council (MRC grant number: MR/R022763/1), Ministério da Saúde (Decit/SECTICS/MS Grant number: 888379/2019 - Portaria Nº 1.949, 04/08/2020) and Banco Industrial do Brasil S/A (CISM grant). Scholarship received financial support from Conselho Nacional de Desenvolvimento Científico e Tecnológico (CAPES). Collaboration between the BHRC and other cohorts has been funded by the National Institutes of Health (NIMH grant number: R01MH120482-01). Involvement of NIMH Intramural investigators has been funded by NIMH-Intramural Research Program Project MH 002782. [uu]CP was supported by a National Health and

Medical Research Council (NHMRC) Senior Principal Research Fellowship (1105825), an NHMRC L3 Investigator Grant (1196508) and NHMRC Program Grant (ID: 1150083). [vv]K.P. is supported by grants: NIMH R01-MH080134, NIMH R01-MH104446 [ww]This research/project is supported by the National Science Foundation (NSF:2010778) and National Research Foundation, Singapore under its AI Singapore Programme (AISG Award No: AISG-GC-2019-002). Additional funding is provided by the Singapore Ministry of Education (Academic research fund Tier 1; NUHSRO/2017/052/T1-SRP-Partnership/01), NUS Institute of Data Science [xx]ADNI: NIH funding. Dataused in preparation of this article were obtained from the Alzheimer's Disease Neuroimaging Initiative (ADNI) database (adni.loni.usc.edu).As such, the investigators within the ADNI contributed to the design and implementation of ADNI and/or provided data but did not participate in analysis or writing of this report. A complete listing of ADNI investigators can be found at:http://adni.loni.usc.edu/wp-content/uploads/how_to_apply/ADNI_Acknowledgement_List.pdf. NACC: The NACC database is funded by NIA/NIH Grant U01 AG016976. NACC data are contributed by the NIA-funded ADCs: P30 AG019610 (PI Eric Reiman, MD), P30 AG013846 (PI Neil Kowall, MD), P50 AG008702 (PI Scott Small, MD), P50 AG025688 (PI Allan Levey, MD, PhD), P50 AG047266 (PI Todd Golde, MD, PhD), P30 AG010133 (PI Andrew Saykin, PsyD), P50 AG005146 (PI Marilyn Albert, PhD), P50 AG005134 (PI Bradley Hyman, MD, PhD), P50 AG016574 (PI Ronald Petersen, MD, PhD), P50 AG005138 (PI Mary Sano, PhD), P30 AG008051 (PI Thomas Wisniewski, MD), P30 AG013854 (PI Robert Vassar, PhD), P30 AG008017 (PI Jeffrey Kaye, MD), P30 AG010161 (PI David Bennett, MD), P50 AG047366 (PI Victor Henderson, MD, MS), P30 AG010129 (PI Charles DeCarli, MD), P50 AG016573 (PI Frank LaFerla, PhD), P50 AG005131 (PI James Brewer, MD, PhD), P50 AG023501 (PI Bruce Miller, MD), P30 AG035982 (PI Russell Swerdlow, MD), P30 AG028383 (PI Linda Van Eldik, PhD), P30 AG053760 (PI Henry Paulson, MD, PhD), P30 AG010124 (PI John Trojanowski, MD, PhD), P50 AG005133 (PI Oscar Lopez, MD), P50 AG005142 (PI Helena Chui, MD), P30 AG012300 (PI Roger Rosenberg, MD), P30 AG049638 (PI Suzanne Craft, PhD), P50 AG005136 (PI Thomas Grabowski, MD), P50 AG033514 (PI Sanjay Asthana, MD, FRCP), P50 AG005681 (PI John Morris, MD), P50 AG047270 (PI Stephen Strittmatter, MD, PhD). [yy]Supported the NIH including the NINDS K23NS101120 (C.K.R.), the NHLBI K23HL141602 (C.M.O), NIBIB R01EB013248 (S.K.W.), R01EB018988 and R01NS106030 (A.G.), and a NHLBI Pediatric Heart Network Scholar Award (C.K.R.); the American Academy of Neurology Clinical Research Training Fellowship (C.K.R.); the Brain and Behavior Research Foundation NARSAD Young Investigator (C.K.R.) and Distinguished Investigator (S.K.W.) Awards; the McKnight Foundation Technological Innovations in Neuroscience Award (A.G.); Office of Faculty Development at Boston Children's Hospital Career Development Awards (A.G., C.K.R.); and the Mend A Heart Foundation (C.M.O.). [zz]RRG was supported by the Guarantors of Brain, Cancer Research UK Cambridge Centre and the EMERGIA Junta de Andalucía program [aaa]LR was supported by the Bernard Wolfe Health Neuroscience Fellowship [bbb]M.D.R. is supported by Bill & Melinda Gates Foundation INV-015711 [ccc]NIHR Cambridge BRC [ddd]T.D.S. is supported by grants: NIH R01MH112847, R01MH120482, & R01MH113550 [eee]M.S. is supported by the Knut and Alice Wallenberg Foundation (Wallenber Centre for Molecular and Translational Medicine), the Swedish Research Council, the Swedish Alzheimer Association, the Swedish Brain Foundation and the Swedish State under the ALF-agreement. [fff]RTS is supported by R01MH112847. [ggg]I.S is supported by the Swedish Research Council (2019-01096 ), Swedish state under the the ALF-agreement, Swedish Brain Foundation, Swedish Alzheimer Foundation [hhh]C.D.S. effort on this project is supported by Bill & Melinda Gates Foundation INV-015711 and NIH P50 HD103525 [iii]RAS is supported by P01 AG036694 and R01 AG03689 [jjj]STRADL study was supported and funded by the Wellcome Trust Strategic Award "Stratifying Resilience and Depression Longitudinally" (ref. 104036/Z/14/Z). Data processing used the resources provided by the Edinburgh Compute and Data Facility (ECDF) (http://www.ecdf.ed.ac.uk/). A.S. was funded as part of the STRADL study and indirectly through the Lister Institute of Preventive Medicine award ref. 173096. [kkk]K.A.T. was supported by the Alzheimer's Society (Grant number 602). [lll]JJT was supported by the Finnish Medical Foundation, Sigrid Juselius Foundation and Emil Aaltonen Foundation [mmm]The preparation and initiation of the i-Share project was funded by the program 'Invest for future' (reference ANR-10-COHO-05). The i-Share Project is currently supported by an unrestricted grant of the Nouvelle-Aquitaine Regional Council (Conseil Régional Nouvelle-Aquitaine) (grant N° 4370420) and by the Bordeaux 'Initiatives d'excellence' (IdEx) program of the University of Bordeaux (ANR-10-IDEX-03-02). It has also received grants from the Nouvelle-Aquitaine Regional Health Agency (Agence Régionale de Santé Nouvelle-Aquitaine, grant N°6066R-8), Public Health France (Santé Publique France, grant N°19DPPP023-0), , and The National Institute against cancer INCa (grant N°INCa_11502). [nnn]SNV was supported by NIH R01MH123563 [ooo]J.W.V. was supported by NIH T32MH019112 [ppp]PEV is a fellow of MQ:Transforming Mental Health (MQF_17_24) [qqq]K.S.W. was supported by the Wellcome Trust (215901/Z/19/Z) [rrr]S.K.W. was supported in part by NIH R01 EB013248 and R01 EB019483. [sss]S.R.W was funded by UKRI Medical Research Council MC_UU_00002/2 and was supported by the NIHR Cambridge Biomedical Research Centre (BRC-1215-20014). The views expressed are those of the author(s) and not necessarily those of the NIHR or the Department of Health and Social Care [ttt]LIFE is funded by means of the European Union, by the European Regional Development Fund (ERDF) and by funds of the Free State of Saxony within the framework of the excellence initiative. A.V.W. was supported by grants from the German Research Foundation (WI 3342/3-1; 209933838-02) [uuu]B.T.T.Y. is supported by the Singapore National Research Foundation (NRF) Fellowship (Class of 2017), the NUS Yong Loo Lin School of Medicine (NUHSRO/2020/124/TMR/LOA), the Singapore National Medical Research Council (NMRC) LCG (OFLCG19May-0035), NMRC STaR (STaR20nov-0003), Singapore Ministry of Health (MOH) Centre Grant (CG21APR1009) and the United States National Institutes of

Health (R01MH120080). [vvv]AZ was supported by an NHMRC Senior Research Fellowship (ID: 1136649) [www]The DCHS was funded by the Bill & Melinda Gates Foundation (OPP1017641). HJZ is funded by the SA-MRC [xxx]AZ was supported by the Swedish Alzheimer Foundation (AF-968431, AF-939988, AF-930582, AF-646061, AF-741361) [yyy]J.H.Z. is funded by National Medical Research Council, Singaore and the National University of Singapore Yong Loo Lin School of Medicine (NUHSRO/2020/124/TMR/LOA) [zzz]X.N.Z has received funding supports from the Child Brain-Mind Development Cohort Study in China Brain Initiative (SQ2021AAA010024), the National Natural Science Foundation of China (81220108014), the National Basic Research (973) Program (2015CB351702), the National Basic Science Data Center 'Chinese Data-sharing Warehouse for In-vivo Imaging Brain' Program (NBSDC-DB-15), the Major Project of National Social Science Foundation of China (20&ZD296), the Beijing Municipal Science and Technology Commission (Z161100002616023, Z181100001518003), the China - Netherlands CAS-NWO Programme (153111KYSB20160020), the Startup Funds for Leading Talents at Beijing Normal University, Guangxi BaGui Scholarship (201621), and the Key Realm R&D Program of Guangdong Province (2019B030335001).

##### Aging Brain: Vasculature, Ischemia, and Behavior Study (ABVIB)

Helena C. Chui M.D.[1,2] (Principal Investigator), Charles C. DeCarli, M.D.[3], William G. Ellis, M.D.[3], William J. Jagust, M.D.[4,5], Joel H. Kramer, Ph.D.[6], Meng Law, M.D.[7,8], Dan Mungas Ph,D.[3], Bruce R. Reed, Ph.D.[9], Nerses Sanossian, M.D.[2,10], Michael W. Weiner, M.D.[11,12,13,14,15,16], Wendy J. Mack, Ph.D.[17], Harry V. Vinters, M.D.[18,19], Chris Zarow, Ph.D.[2], Ling Zheng, Ph.D.[2]

[1]Alzheimer's Disease Research Center, Keck School of Medicine, University of Southern California, Los Angeles, CA, USA [2]Department of Neurology, Keck School of Medicine, University of Southern California, Los Angeles, CA, USA [3]Department of Neurology, University of California Davis School of Medicine, Sacramento, California, USA [4]Department of Neuroscience, University of California Berkeley, Berkeley, CA, USA [5]Lawrence Berkeley National Laboratory, Berkeley, CA, USA [6]Memory and Aging Center, Department of Neurology, University of California, San Francisco, San Francisco, CA, USA [7]Department of Radiology, The Alfred, Melbourne, VIC, Australia [8]Department of Neuroscience, School of Translational Medicine, Monash University, Clayton, VIC, Australia [9]National Institutes of Health Center for Scientific Review, Bethesda, MD, USA [10]Roxanna Todd Hodges Stroke Program, University of Southern California, Los Angeles, CA, USA [11]Department of Veterans Affairs Medical Center, Center for Imaging of Neurodegenerative Diseases, San Francisco, California, USA [12]Department of Radiology and Biomedical Imaging, University of California San Francisco, San Francisco, California, USA [13]Department of Medicine, University of California San Francisco, San Francisco, California, USA [14]Department of Psychiatry and Behavioral Sciences, University of California San Francisco, San Francisco, California, USA [15]Department of Neurology, University of California San Francisco, San Francisco, California, USA [16]Northern California Institute for Research and Education (NCIRE), San Francisco, California, USA [17]Population and Public Health Sciences, Keck School of Medicine of the University of Southern California, Los Angeles, California, USA [18]Department of Pathology and Laboratory Medicine, David Geffen School of Medicine, University of California Los Angeles, Los Angeles, California, USA [19]Department of Neurology, David Geffen School of Medicine, University of California Los Angeles, Los Angeles, California, USA

##### Alzheimer's Disease Neuroimaging Initiative (ADNI)

A complete listing of ADNI investigators can be found at: `http://adni.loni.usc.edu/wp-content/uploads/how_to_apply/ADNI_Acknowledgement_List.pdf`

##### Australian Imaging, Biomarkers and Lifestyle (AIBL)

AIBL researchers are listed at `www.aibl.csiro.au`.

##### Alzheimer's Disease Repository Without Borders (ARWiBo)

The Principal Investigator of ARWIBO is Giovanni B. Frisoni, MD, University Hospitals and University of Geneva, Geneva, Switzerland, and IRCCS Fatebenefratelli, The National Centre for Alzheimer's and Mental Diseases, Brescia, Italy. ARWIBO is the result of effort of many researchers of IRCCS Fatebenefratelli: G. Binetti, MD, Neurobiology; L. Bocchio-Chiavetto, PhD, Neuropharmacology; M. Cotelli, PhD, Neuropsychology Unit; C. Minussi, PhD, Neurophysiology; M. Gennarelli, PhD, Genetic Unit; R. Ghidoni, PhD, Proteomics Unit; D. Moretti, MD, and O. Zanetti, MD, Alzheimer's Unit. A complete listing of ARWiBo researchers can be found at: `https://www.arwibo.it/acknowledgments.pdf`

##### Biomarkers of Cognitive Decline among Normal Individuals: the BIOCARD Study

A listing of BIOCARD investigators can be found on the BIOCARD website, `https://biocard.pathology.jhu.edu/our-team/`.

#### Centre for Attention Learning and Memory (CALM)

Duncan E. Astle[1,2]. More information on CALM team members can be found at: https://calm.mrc-cbu.cam.ac.uk/team/

[1]MRC Cognition and Brain Sciences Unit, University of Cambridge, Cambridge, UK [2]Department of Psychiatry, University of Cambridge, Cambridge, UK

#### Cambridge Center for Ageing and Neuroscience (Cam-CAN) study

A complete list of Cam-CAN reasearchers can be found at: https://cam-can.mrc-cbu.cam.ac.uk/people/

#### Chinese Color Nest Project (CCNP)

Yin-Shan Wang[1,2], Ning Yang[1] and Xi-Nian Zuo[1,2,3,4]

[1]State Key Laboratory of Cognitive Neuroscience and Learning, Beijing Normal University, Beijing 100875, China [2]Developmental Population Neuroscience Research Center, International Data Group/McGovern Institute for Brain Research, Beijing Normal University, Beijing 100875, China [3]Research Center for Lifespan Development of Mind and Brain, Institute of Psychology, Chinese Academy of Sciences, Beijing 100101, China [4]Department of Psychology, University of Chinese Academy of Sciences, Beijing 100049, China

#### Centers of Biomedical Research Excellence (COBRE)

Vince D. Calhoun[1]

[1]Tri-Institutional Center for Translational Research in Neuroimaging and Data Science (TReNDS), Georgia State University, Georgia Institute of Technology, and Emory University, Atlanta, Georgia, USA

#### Developing Human Connectome Project (dHCP)

Stephen M. Smith[1], Daniel Rueckert[2,3], Joseph V. Hajnald[4,5] and A. David Edwards[6,7]

[1]Nuffield Department of Clinical Neurosciences, Wellcome Centre for Integrative Neuroimaging, FMRIB, University of Oxford, Oxford, UK [2]Chair for AI in Healthcare and Medicine, Technical University of Munich (TUM) and TUM University Hospital, Munich, Germany [3]Department of Computing, Imperial College London, London, UK [4]Early Life Imaging Department, School of Biomedical Engineering and Imaging Sciences, King's College London, London, UK [5]Imaging Physics and Engineering Department, School of Biomedical Engineering and Imaging Sciences, King's College London, London, UK [6]Centre for the Developing Brain, Research Department of Early Life Imaging, School of Biomedical Engineering and Imaging Sciences, King's College London, London SE1 7EH, United Kingdom [7]Medical Research Council Centre for Neurodevelopmental Disorders, King's College London, London SE1 1UL, United Kingdom

#### Harvard Aging Brain Study (HABS)

Reisa A. Sperling[1,2] and Keith A. Johnson[1,2,3,4,5]

[1]Department of Neurology, Massachusetts General Hospital, Harvard Medical School, Boston, MA, USA [2]Department of Neurology, Brigham and Women's Hospital, Harvard Medical School, Boston, MA, USA [3]Molecular Neuroimaging, Massachusetts General Hospital, Boston, Massachusetts, USA [4]arvard Medical School, Boston, Massachusetts, USA [5]Department of Radiology, Massachusetts General Hospital, Boston, Massachusetts, USA

#### Human Connectome Project (HCP)

The investigators for each HCP substudy can be found at: https://www.humanconnectome.org/

#### International Consortium for Brain Mapping (ICBM)

John C Mazziotta[1]

[1]UCLA School of Medicine, Los Angeles, USA

#### IMAGEN

Sylvane Desrivieres[1], Andreas Heinz[2], Tianye Jia1[3,4], and Gunter Schumann[5,6]

[1]Social, Genetic and Developmental Psychiatry Centre, Institute of Psychiatry, Psychology and Neuroscience, King's College, London, UK [2]Department of Psychiatry and Psychotherapy CCM, Charité-Universitätsmedizin Berlin, corporate member of Freie Universität Berlin, Humboldt-Universität zu Berlin, and Berlin Institute of Health, Berlin,

Germany [3]Institute of Science and Technology for Brain-Inspired Intelligence (ISTBI), Fudan University, Shanghai 200433, China [4]Key Laboratory of Computational Neuroscience and Brain-Inspired Intelligence, Fudan University, Ministry of Education, Shanghai 200433, China. [5]Centre for Population Neuroscience and Stratified Medicine (PONS), Department of Psychiatry and Neuroscience, Charité Universitätsmedizin Berlin, Berlin, Germany [6]Centre for Population Neuroscience and Precision Medicine (PONS), Institute for Science and Technology of Brain-inspired Intelligence (ISTBI), Fudan University, Shanghai, China

##### Neuroscience in Psychiatry Network (NSPN)

A full list of NSPN consortium members can be found at: `https://www.nspn.org.uk/nspn-team/`

##### Province of Ontario Neurodevelopmental Disorders (POND) Network

Evdokia Anagnostou[1,2], Muhammad Ayub[3,4], Jennifer Crosbie[5,6,7,8], Christopher F. Hammill[9], Elizabeth A. Kelley[3,10,11], Jason Lerch[7,9,12], and Russell J. Schachar[5,6,7,8]

[1]Autism Research Centre, Bloorview Research Institute, Holland Bloorview Kids Rehabilitation Hospital, Toronto, ON, Canada [2]Department of Pediatrics, Temerty Faculty of Medicine, University of Toronto, Toronto, Ontario, Canada [3]Department of Psychiatry, Queen's University, Kingston, ON, Canada [4]Division of Psychiatry, University College London, London, UK [5]Department of Psychiatry, The Hospital for Sick Children, Toronto, ON, Canada [6]Department of Psychiatry, Temerty Faculty of Medicine, University of Toronto, Toronto, ON, Canada [7]Program in Neurosciences and Mental Health, Research Institute, The Hospital for Sick Children, Toronto, ON, Canada [8]Genetics and Genome Biology, The Hospital for Sick Children, Toronto, ON, Canada [9]Mouse Imaging Centre, Hospital for Sick Children, Toronto, ON, Canada [10]Department of Psychology, Queen's University, Kingston, ON, Canada [11]Centre for Neuroscience Studies, Queen's University, Kingston, ON, Canada [12]Wellcome Centre for Integrative Neuroimaging, FMRIB, Nuffield Department of Clinical Neurosciences, University of Oxford, Oxford, UK

##### Pre-symptomatic Evaluation of Experimental or Novel Treatments for Alzheimer Disease (PREVENT-AD)

Alexa Pichet Binette[1] and Sylvia Villeneuve[2,3]

[1]Clinical Memory Research Unit, Department of Clinical Sciences Malmö, Lund University, Lund, Sweden [2]Douglas Mental Health University Institute Research Centre, McGill University, Montréal H4H 1R3, Canada [3]Department of Psychiatry, McGill University, Montréal H3A 1A1, Canada

##### Vietnam Era Twin Study of Aging (VETSA)

A complete list of VETSA researchers can be found at: `https://www.vetsatwins.org/people/`

### Supplementary methods

#### Aggregated lifespan dataset

##### Demographics

We aggregated data across 128 primary cross-sectional and longitudinal neuroimaging studies, covering an age range from mid gestation (180 days post conception), to 100 years. Details of each individual study are compiled further below.

We collected a total of 179,151 scans from 139,842 individual subjects across 128 studies.

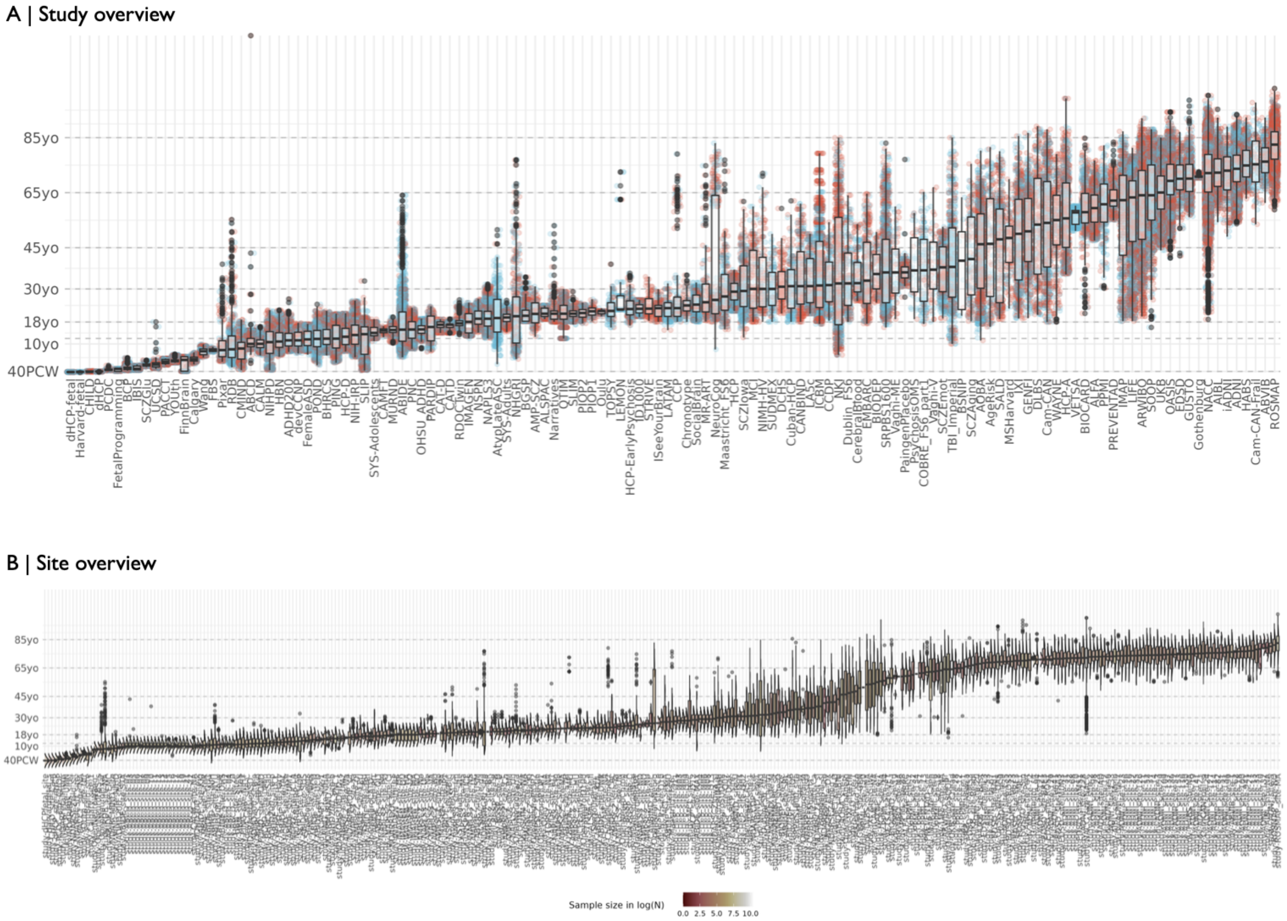

**Fig. 1: Aggregated lifespan sample:** We collected T1- and T2-weighted MRI data from 128 primary studies to form an aggregated dataset of 179,151 scans from 139,842 subjects that collectively spanned the age range from mid-gestation to 102 postnatal years. (A) The box-plots show the age distribution for each study, with individual points colored by sex. (B) Many studies were acquired at multiple scanning locations, or sites. Here, we show the size and median age of all sites included.

| Study | Sites | Session | Age range | $N \| N_{female}$ | $N_{pat.}$ | $pipeline_{GM}$ | $pipeline_{SA}$ | $pipeline_{CT}$ | Reference |
|---|---|---|---|---|---|---|---|---|---|
| ABCD | 22 | 3 | 8y - 17y | 11791 \| 5634 f | 2985 | FS7_T1 | FS7_T1 | FS7_T1 | [27], http://dx.doi.org/10.15154/3er3-dc69 |
| ABIDE | 25 | 1 | 5y - 64y | 1865 \| 389 f | 985 | FS6_T1 | FS6_T1 | FS6_T1 | [35] |
| ABVIB | 1 | 3 | 61y - 92y | 197 \| 76 f | 214 | FS6_T1 | FS6_T1 | FS6_T1 | [99] |
| ADHD200 | 10 | 2 | 7y - 26y | 931 \| 356 f | 363 | FS6_T1 | FS6_T1 | FS6_T1 | [12] |
| ADNI | 3 | 10 | 54y - 93y | 1704 \| 767 f | 1291 | FS6_T1 | FS6_T1 | FS6_T1 | [60] |
| AIBL | 3 | 5 | 54y - 95y | 624 \| 356 f | 175 | FS6_T1 | FS6_T1 | FS6_T1 | [38] |
| ALFA | 1 | 2 | 48y - 77y | 1285 \| 816 f | 0 | FS7_T1+ | FS7_T1+ | FS7_T1+ | [84] |
| ALSPAC | 2 | 1 | 19y - 24y | 423 \| 257 f | 155 | FS6_T1 | FS6_T1 | FS6_T1 | [85] |
| AMP-SCZ | 14 | 1 | 13y - 31y | 137 \| 79 f | 112 | SS | RAC | | [147], http://dx.doi.org/10.15154/3er3-dc69 |
| AOBA | 1 | 1 | 11y - 80y | 1139 \| 586 f | 180 | FS6_T1 | FS6_T1 | FS6_T1 | [115] |
| ARWIBO | 3 | 3 | 20y - 92y | 1827 \| 1147 f | 758 | FS6_T1 | FS6_T1 | FS6_T1 | [41] |
| AtypLateASC | 1 | 5 | 3y - 51y | 60 \| 8 f | 41 | SS | RAC | | http://dx.doi.org/10.15154/3er3-dc69 |
| BCP | 1 | 6 | 0y - 5y | 119 \| 52 f | 0 | FS6_T1, SS | FS6_T1, RAC | RAC | [59] |
| BGSP | 4 | 1 | 18y - 34y | 1557 \| 897 f | 0 | FS6_T1 | FS6_T1 | FS6_T1 | [57] |
| BHRCS | 2 | 3 | 6y - 22y | 719 \| 355 f | 461 | FS6_T1 | FS6_T1 | FS6_T1 | [114] |
| BIOCARD | 1 | 5 | 21y - 85y | 206 \| 131 f | 206 | FS7_T1+ | FS7_T1+ | FS7_T1+ | [5] |
| BIODEP | 1 | 1 | 25y - 51y | 99 \| 70 f | 58 | FS6_T1 | FS6_T1 | FS6_T1 | [66] |
| Calgary | 1 | 7 | 3y - 7y | 82 \| 37 f | 0 | FS6_T1, SS | FS6_T1, RAC | FS6_T1, RAC | [105] |
| CALM | 1 | 1 | 6y - 16y | 385 \| 135 f | 311 | FS6_T1, FS6_T1+ | FS6_T1, FS6_T1+ | FS6_T1, FS6_T1+ | [58] |
| Cam-CAN | 1 | 1 | 18y - 88y | 632 \| 324 f | 0 | FS5.3 | FS5.3 | FS5.3 | [120, 131] |
| Cam-CAN-Frail | 1 | 1 | 53y - 98y | 122 \| 55 f | 74 | FS7_T1 | FS7_T1 | FS7_T1 | [67] |
| CAMFT | 1 | 1 | 12y - 16y | 68 \| 35 f | 0 | FS6_T1 | FS6_T1 | FS6_T1 | [78] |
| CANBIND | 3 | 1 | 18y - 61y | 275 \| 181 f | 178 | FS7_T1 | FS7_T1 | FS7_T1 | [71] |
| CHILD | 1 | 2 | 10PCW - 3m | 43 \| 22 f | 11 | Custom | Custom | Custom | [144] |
| CCP | 1 | 1 | 18y - 79y | 176 \| 84 f | 0 | SS | RAC | | [48] |
| COBRE_FS6 | 1 | 1 | 17y - 65y | 146 \| 35 f | 70 | FS6_T1 | FS6_T1 | FS6_T1 | [4] |
| Cuban-HCP | 1 | 1 | 18y - 62y | 181 \| 44 f | 0 | FS6_T1 | FS6_T1 | FS6_T1 | [138] |
| DCHS | 1 | 2 | 21y - 59y | 127 \| 127 f | 0 | FS7_T1 | FS7_T1 | FS7_T1 | |
| devCCNP | 1 | 1 | 7y - 18y | 176 \| 98 f | 0 | FS7_T1 | FS7_T1 | FS7_T1 | [155, 75, 39] |
| dHCP | 1 | 2 | 11PCW - 1m | 505 \| 241 f | 0 | SS | RAC | Custom | [79, 36] |
| dHCP-fetal | 1 | 3 | 19PCW - 0m | 246 \| 112 f | 0 | SS | RAC | Custom | [121] |
| DLBS | 1 | 1 | 21y - 89y | 296 \| 186 f | 0 | FS6_T1 | FS6_T1 | FS6_T1 | [65] |
| Dublin_FS6 | 1 | 2 | 18y - 64y | 206 \| 94 f | 56 | FS6_T1 | FS6_T1 | FS6_T1 | [87, 86] |
| EDSD | 5 | 1 | 42y - 87y | 381 \| 204 f | 221 | FS6_T1 | FS6_T1 | FS6_T1 | [22] |
| FinnBrain | 1 | 3 | 0y - 6y | 245 \| 119 f | 0 | SS | RAC | FSInfant | [29] |
| GENFI | 5 | 3 | 19y - 87y | 365 \| 204 f | 110 | FS6_T1 | FS6_T1 | FS6_T1 | [108] |
| Gothenburg | 1 | 1 | 70y - 72y | 724 \| 381 f | 0 | FS7_T1 | FS7_T1 | FS7_T1 | [112] |
| GUSTO | 1 | 6 | 56y - 98y | 301 \| 157 f | 0 | FS7_T1 | FS7_T1 | FS7_T1 | [73] |
| HABS | 1 | 4 | 63y - 92y | 280 \| 167 f | 7 | FS6_T1 | FS6_T1 | FS6_T1 | [31] |
| Harvard-fetal | 1 | 1 | 19PCW - -1m | 32 \| 9 f | 0 | Custom | Custom | | [142, 109, 143] |

| Study | Sites | Session | Age range | $N \vert N_{female}$ | $N_{pat.}$ | $pipeline_{GM}$ | $pipeline_{SA}$ | $pipeline_{CT}$ | Reference |
|---|---|---|---|---|---|---|---|---|---|
| HBN | 3 | 1 | 5y - 22y | 1007 \| 407 f | 563 | Custom | Custom | Custom | [7] |
| HCP | 13 | 1 | 22y - 37y | 1113 \| 606 f | 0 | FS5.3 | FS5.3 | FS5.3 | [49] |
| HCP-A | 1 | 1 | 35y - 99y | 667 \| 380 f | 0 | FS6_T1+ | FS6_T1+ | FS6_T1+ | [56], http://dx.doi.org/10.15154/3er3-dc69 |
| HCP-D | 4 | 1 | 7y - 21y | 652 \| 330 f | 0 | FS6_T1+ | FS6_T1+ | FS6_T1+ | [56], http://dx.doi.org/10.15154/3er3-dc69 |
| HCP-EarlyPsych | 3 | 1 | 17y - 36y | 174 \| 66 f | 120 | SS | RAC | | http://dx.doi.org/10.15154/3er3-dc69 |
| iADNI | 5 | 1 | 49y - 89y | 201 \| 143 f | 235 | FS6_T1 | FS6_T1 | FS6_T1 | [28] |
| ICBM | 1 | 1 | 17y - 79y | 565 \| 658 f | 0 | FS6_T1 | FS6_T1 | FS6_T1 | [82] |
| IMAGEN | 8 | 3 | 12y - 25y | 1942 \| 1008 f | 0 | FS6_T1, FS6_T1+ | FS6_T1, FS6_T1+ | FS6_T1, FS6_T1+ | [118], https://github.com/imagen2/imagen˙mri/tre |
| IMAP | 1 | 3 | 18y - 87y | 342 \| 171 f | 131 | FS6_T1 | FS6_T1 | FS6_T1 | [45] |
| ISeeYourBrain | 1 | 1 | 18y - 30y | 202 \| 149 f | 0 | FS7_T1 | FS7_T1 | FS7_T1 | [43, 44] |
| IXI | 3 | 1 | 19y - 86y | 560 \| 327 f | 0 | FS6_T1, FS6_T1+ | FS6_T1, FS6_T1+ | FS6_T1, FS6_T1+ | [70], https://brain-development.org/ixi-dataset/ |
| LATAM | 1 | 1 | 15y - 32y | 72 \| 22 f | 0 | FS6_T1 | FS6_T1 | FS6_T1 | [30] |
| LEMON | 1 | 1 | 22y - 72y | 82 \| 14 f | 0 | FS7_T1+ | FS7_T1+ | FS7_T1+ | [11] |
| LIFE | 1 | 1 | 18y - 81y | 2525 \| 1207 f | 134 | FS5.3 | FS5.3 | FS5.3 | [77] |
| Maastricht_FS6 | 1 | 1 | 15y - 77y | 255 \| 122 f | 112 | FS6_T1 | FS6_T1 | FS6_T1 | [87] |
| MCI | 3 | 1 | 18y - 60y | 194 \| 141 f | 104 | FS6_T1 | FS6_T1 | FS6_T1 | [53, 52] |
| NACC | 18 | 10 | 19y - 100y | 5161 \| 3111 f | 2126 | FS6_T1, FS6_T1+ | FS6_T1, FS6_T1+ | FS6_T1, FS6_T1+ | [14] |
| NAPLS3 | 4 | 6 | 12y - 32y | 503 \| 208 f | 433 | SS | RAC | | [3] |
| ND_EMBARC | 1 | 1 | 17y - 65y | 316 \| 208 f | 277 | FS6_T1 | FS6_T1 | FS6_T1 | http://dx.doi.org/10.15154/3er3-dc69 |
| ND_FemaleASD | 3 | 3 | 6y - 17y | 388 \| 212 f | 143 | FS6_T1 | FS6_T1 | FS6_T1 | http://dx.doi.org/10.15154/3er3-dc69 |
| ND_FetalProgram | 1 | 2 | 0y - 1y | 56 \| 20 f | 0 | SS | RAC | FSInfant | http://dx.doi.org/10.15154/3er3-dc69 |
| ND_IBIS | 4 | 3 | 0y - 3y | 224 \| 79 f | 31 | SS | RAC | RAC | [74], http://dx.doi.org/10.15154/3er3-dc69 |
| ND_NHGRI | 1 | 1 | 4y - 77y | 401 \| 183 f | 0 | FS6_T1 | FS6_T1 | FS6_T1 | https://nda.nih.gov/edit˙collection.html?id=2936, h |
| ND_NIH-IRP | 1 | 6 | 5y - 26y | 769 \| 411 f | 0 | FS7_T1 | FS7_T1 | FS7_T1 | [104], http://dx.doi.org/10.15154/3er3-dc69 |
| ND_OHSU_ADHD | 1 | 1 | 8y - 19y | 218 \| 389 f | 614 | FS7_T1+ | FS7_T1+ | FS7_T1+ | [103], http://dx.doi.org/10.15154/3er3-dc69 |
| ND_PARDIP | 2 | 1 | 7y - 24y | 107 \| 50 f | 66 | FS7_T1 | FS7_T1 | FS7_T1 | http://dx.doi.org/10.15154/3er3-dc69 |
| ND_SCZIowa | 4 | 5 | 12y - 62y | 205 \| 75 f | 109 | FS6_T1 | FS6_T1 | FS6_T1 | [10],http://dx.doi.org/10.15154/3er3-dc69 |
| ND_UCSD | 1 | 3 | 1y - 18y | 340 \| 177 f | 117 | FS6_T1, SS | FS6_T1, RAC | FS6_T1, RAC | [40], http://dx.doi.org/10.15154/3er3-dc69 |
| ND_CCDP | 1 | 1 | 18y - 54y | 203 \| 72 f | 148 | SS | RAC | | http://dx.doi.org/10.15154/3er3-dc69 |
| ND_CMIND | 2 | 5 | 0y - 19y | 177 \| 114 f | 0 | SS | RAC | | [139], http://dx.doi.org/10.15154/3er3-dc69 |
| ND_MURND | 1 | 1 | 10y - 20y | 339 \| 192 f | 0 | SS | RAC | | http://dx.doi.org/10.15154/3er3-dc69 |
| ND_PsychCB | 1 | 1 | 26y - 53y | 68 \| 98 f | 111 | SS | RAC | | http://dx.doi.org/10.15154/3er3-dc69 |
| ND_PsychosisOMS | 1 | 1 | 19y - 58y | 38 \| 10 f | 18 | SS | RAC | | http://dx.doi.org/10.15154/3er3-dc69 |
| ND_RDOCTwin | 1 | 1 | 13y - 26y | 360 \| 188 f | 0 | SS | RAC | | [51], http://dx.doi.org/10.15154/3er3-dc69 |
| ND_SCZAging | 1 | 1 | 18y - 75y | 160 \| 74 f | 116 | SS | RAC | | http://dx.doi.org/10.15154/3er3-dc69 |
| ND_SCZEmot | 1 | 5 | 19y - 53y | 149 \| 69 f | 104 | SS | RAC | | http://dx.doi.org/10.15154/3er3-dc69 |
| ND_SCZGlu | 1 | 1 | 2y - 4y | 86 \| 24 f | 53 | SS | RAC | | http://dx.doi.org/10.15154/3er3-dc69 |
| ND_SocialBrain | 1 | 1 | 18y - 32y | 130 \| 212 f | 337 | SS | RAC | | http://dx.doi.org/10.15154/3er3-dc69 |
| NIHPD | 1 | 8 | 0y - 22y | 389 \| 197 f | 0 | FS6_T1, SS | FS6_T1, RAC | FS6_T1, RAC | [8] |
| NKI | 1 | 3 | 6y - 85y | 666 \| 435 f | 0 | FS6_T1 | FS6_T1 | FS6_T1 | [134] |

| Study | Sites | Session | Age range | $N \mid N_{female}$ | $N_{pat.}$ | $pipeline_{GM}$ | $pipeline_{SA}$ | $pipeline_{CT}$ | Reference |
|---|---|---|---|---|---|---|---|---|---|
| NSPN | 3 | 3 | 14y - 26y | 294 \| 149 f | 0 | FS5.3 | FS5.3 | FS5.3 | [151] |
| OASIS | 2 | 6 | 41y - 94y | 895 \| 820 f | 484 | FS6_T1, FS6_T1+ | FS6_T1, FS6_T1+ | FS6_T1, FS6_T1+ | [72] |
| ON_CAT-D | 1 | 3 | 14y - 19y | 124 \| 142 f | 167 | FS7_T1 | FS7_T1 | FS7_T1 | [93, 113, 26] |
| ON_Chronotype | 1 | 1 | 18y - 35y | 110 \| 70 f | 0 | FS7_T1 | FS7_T1 | FS7_T1 | [153, 154] |
| ON_ID1000 | 1 | 1 | 20y - 26y | 926 \| 1438 f | 0 | FS6_T1 | FS6_T1 | FS6_T1 | [122, 125] |
| ON_LA5c | 2 | 1 | 21y - 50y | 257 \| 108 f | 136 | FS6_T1 | FS6_T1 | FS6_T1 | [16, 101] |
| ON_MR-ART | 1 | 1 | 18y - 75y | 142 \| 94 f | 0 | FS7_T1 | FS7_T1 | FS7_T1 | [89, 90] |
| ON_Narratives | 1 | 1 | 18y - 53y | 295 \| 177 f | 0 | FS7_T1 | FS7_T1 | FS7_T1 | [91, 92] |
| ON_NeuroCog | 2 | 1 | 8y - 83y | 287 \| 161 f | 0 | FS7_T1 | FS7_T1 | FS7_T1 | [119, 126] |
| ON_NIMH-HV | 1 | 1 | 18y - 72y | 143 \| 95 f | 0 | FS7_T1 | FS7_T1 | FS7_T1 | [95, 96] |
| ON_PIOP1 | 1 | 1 | 18y - 26y | 72 \| 43 f | 0 | FS6_T1+ | FS6_T1+ | FS6_T1+ | [125, 123] |
| ON_PIOP2 | 1 | 1 | 17y - 25y | 224 \| 128 f | 0 | FS6_T1 | FS6_T1 | FS6_T1 | [125, 124] |
| ON_Pixar | 1 | 1 | 4y - 39y | 152 \| 83 f | 0 | FS7_T1 | FS7_T1 | FS7_T1 | [107, 106] |
| ON_SUDMEX | 1 | 1 | 18y - 50y | 130 \| 20 f | 71 | FS7_T1 | FS7_T1 | FS7_T1 | [46, 102, 47] |
| ON_Wang | 1 | 3 | 5y - 10y | 215 \| 125 f | 0 | FS7_T1 | FS7_T1 | FS7_T1 | [145, 146] |
| ON_AgeRisk | 1 | 1 | 16y - 81y | 182 \| 95 f | 0 | SS | RAC | | [133, 132] |
| ON_CATD | 1 | 3 | 14y - 19y | 121 \| 147 f | 175 | SS | RAC | | [113, 93] |
| ON_FBS | 1 | 1 | 7y - 9y | 58 \| 27 f | 0 | SS | RAC | | [42] |
| ON_Paingen | 1 | 1 | 27y - 77y | 383 \| 228 f | 0 | SS | RAC | | [20, 19] |
| ON_SOOP | 1 | 1 | 16y - 88y | 1030 \| 487 f | 1030 | SS | RAC | | [110] |
| Oulu | 1 | 1 | 20y - 23y | 101 \| 64 f | 0 | FS6_T1 | FS6_T1 | FS6_T1 | [63] |
| PACCT | 1 | 1 | 2y - 5y | 242 \| 125 f | 0 | FS7_T1 | FS7_T1 | FS7_T1 | [94] |
| PCDC | 1 | 5 | 0y - 2y | 44 \| 28 f | 0 | SS | RAC | RAC | [15] |
| PING | 4 | 1 | 3y - 21y | 713 \| 352 f | 0 | FS7_T1, FS7_T1+ | FS7_T1, FS7_T1+ | FS7_T1, FS7_T1+ | [61], http://dx.doi.org/10.15154/3er3-dc69 |
| PNC | 1 | 1 | 8y - 23y | 1509 \| 797 f | 0 | FS6_T1 | FS6_T1 | FS6_T1 | [116] |
| POND | 1 | 6 | 2y - 24y | 605 \| 171 f | 511 | FS6_T1 | FS6_T1 | FS6_T1 | [88] |
| PPMI | 3 | 1 | 30y - 82y | 239 \| 86 f | 157 | FS7_T1 | FS7_T1 | FS7_T1 | [80, 9] |
| PREVENTAD | 1 | 1 | 55y - 83y | 369 \| 273 f | 0 | FS6_T1 | FS6_T1 | FS6_T1 | [136] |
| QTIM | 1 | 1 | 12y - 30y | 1042 \| 669 f | 0 | FS7_T1 | FS7_T1 | FS7_T1 | [128] |
| RDB | 1 | 1 | 1y - 55y | 492 \| 126 f | 325 | FS6_T1 | FS6_T1 | FS6_T1 | [135] |
| ROSMAP | 2 | 7 | 59y - 103y | 858 \| 673 f | 183 | FS6_T1 | FS6_T1 | FS6_T1 | [13] |
| SALD | 1 | 1 | 19y - 80y | 472 \| 295 f | 0 | FS6_T1 | FS6_T1 | FS6_T1 | [149] |
| SLIM | 1 | 3 | 17y - 29y | 650 \| 342 f | 0 | FS6_T1 | FS6_T1 | FS6_T1 | [76] |
| SLIP | 12 | 3 | 0y - 33y | 1657 \| 1185 f | 0 | SS | RAC | RAC | [117] |
| SRPBS1600 | 12 | 1 | 16y - 80y | 1548 \| 638 f | 638 | FS7_T1 | FS7_T1 | FS7_T1 | [130, 152] |
| STRIVE | 1 | 1 | 18y - 34y | 84 \| 84 f | 54 | FS6_T1 | FS6_T1 | FS6_T1 | [150] |
| SYS-Adolescents | 1 | 1 | 11y - 19y | 1029 \| 533 f | 0 | FS5.3 | FS5.3 | FS5.3 | [100] |
| SYS-Adults | 1 | 1 | 14y - 26y | 670 \| 363 f | 0 | FS5.3 | FS5.3 | FS5.3 | [100] |
| TBI_Imperial | 1 | 4 | 10y - 85y | 347 \| 129 f | 208 | FS6_T1 | FS6_T1 | FS6_T1 | [21, 54] |
| TOPSY | 1 | 1 | 16y - 39y | 105 \| 25 f | 74 | FS6_T1 | FS6_T1 | FS6_T1 | [33] |

| Study | Sites | Session | Age range | $N \| N_{female}$ | $N_{pat.}$ | $pipeline_{GM}$ | $pipeline_{SA}$ | $pipeline_{CT}$ | Reference |
|---|---|---|---|---|---|---|---|---|---|
| UKB | 4 | 2 | 44y - 85y | 65517 \| 34312 f | 12125 | FS7_T1 | FS7_T1 | FS7_T1 | [83] |
| Vaghi-ME | 1 | 1 | 19y - 60y | 83 \| 40 f | 42 | FS5.3 | FS5.3 | FS5.3 | [137] |
| Vaghi-V | 1 | 1 | 18y - 67y | 53 \| 31 f | 21 | FS5.3 | FS5.3 | FS5.3 | |
| VETSA | 2 | 1 | 51y - 60y | 528 \| 0 f | 0 | FS6_T1 | FS6_T1 | FS6_T1 | [69] |
| WAYNE | 1 | 2 | 18y - 79y | 198 \| 132 f | 0 | FS6_T1 | FS6_T1 | FS6_T1 | [32] |
| YOUth | 1 | 2 | 3y - 6y | 1117 \| 644 f | 0 | FS7_T1 | FS7_T1 | FS7_T1 | [97] |

**Table 1: Dataset overview:** Above, we summarize datasets included in the LBCC cohort. For each study, we list the number of sites ($Sites$), and the maximum sessions (timepoints) ($Sesssion$), the age range, the total number of individual subjects ($N$), as well as the number of females included ($N_{female}$), the number of with a neuropsychiatric or neurodevelopmental diagnosis ($N_{patient}$). We further specify the processing pipeline used for each phenotype: cortical and subcortical g volume ($pipeline_{GM}$), surface area ($pipeline_{SA}$) and cortical thickness ($pipeline_{CT}$). We lastly reference relevant resources for each study. ND = NDAR stud OpenNeuro studies.

Previous work has defined major epochs of lifespan brain development, ranging from conception to old age [64]. No neuroimaging data is available prior to about 16 PCW, but our aggregated dataset includes subjects from all developmental stages from 24 PCW: it includes 265 scans from the *late fetal* (24-38 PCW) stage, 557 scans from *early infancy* (0-6 months), 242 scans from *late infancy* (6 months - 1 year), 3,106 scans from *early childhood* (1-6 years), 17,851 scans from *mid to late childhood* (6-12 years), 13,332 scans from *adolescence* (12-20 years), 13,607 scans from *young adulthood* (20-40 years), 27,196 scans from *mid adulthood* (40-60 years), and 62,649 scans from *late adulthood* (older than 60 years). Notably, the ages <2 years, as well as the mid-life range from 25 to 50 years, spanning two developmental stages, are represented by comparatively few scans.

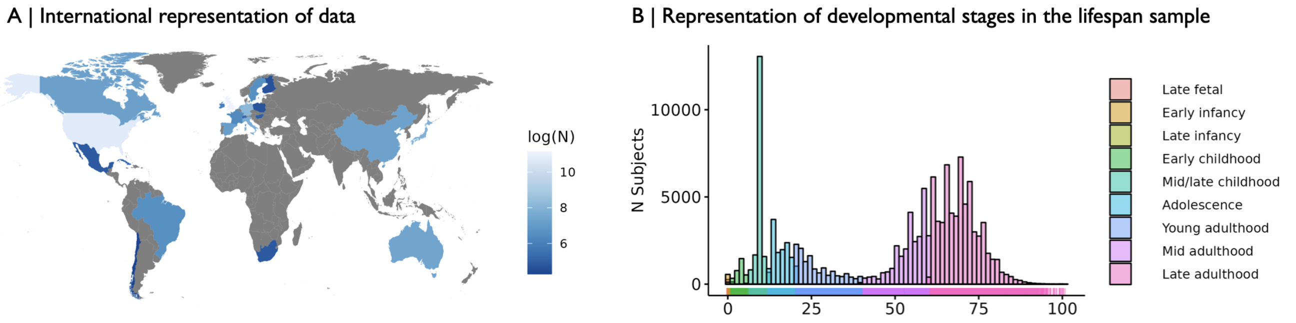

**Fig. 2: Demographic over view of the lifespan sample:** (A) Histogram and density plot indicating the number of subjects by age. Further, the coloring indicates which developmental stage the subject falls into. Notably, the dataset includes fewer scans early in life, in the prenatal window until early childhood, as well as between 25 and 50 years of age.

An ideal sample to estimate generalisable brain "growth charts" should be as representative of the world's population as possible, and cover the entire lifespan. However, in general, the landscape of openly available neuroimaging datasets is strongly biased in favour of (mostly affluent) participants from white Western European and North American populations [68]. This increasingly well-recognised demographic bias in the field at large was inevitably also reflected in our aggregated dataset. While we were able to gather data from three continents, further work is needed to increase the geographical, and thereby ethnic, diversity of the studies included in this work to enable more population-representative normative trajectories.

#### Site aggregation

Many studies included in this work are multi-center (multi-site) studies. While some studies included multi-site acquisition as per stuy protocol, often as part of larger study consortia (i.e. ABCD, UKB), a number of studies, however, included multi-site data, that was closer to multi-study data, i.e. indepently acquired data were post-hoc combined to form a single larger dataset. Further, some studies had very small sample sizes as some sites. Prior work on braincharts has controlled for variability at the study level [15]. In an effort to enhance our ability to derive accurate centile scores for individuals from multi-center studies, here, we aimed to control for random effects of site. To ensure better stability of our models, we opted to include the largest sample size possible, while still controlling for site. We therefore proceeded in a two step approach. First, we found all sites with less than 40 subjects. For each study that had such sites, we combined all subjects from such small sites, to form an aggregated artificial site. Then, we re-estimated the number of subjects per site. In cases where studies still included sites with less than 30 subjects, we opted to remove these sites from our analyses. This process means that subjects from the new artificial aggregated sites can still contribute to the population trajectories, while we accept the fact that there centile scores will be somewhat less accurate.

#### Preprocessing

The primary studies included in this dataset ranged from openly available data, to data only shareable in derived format, i.e. regional volume values rather than raw imaging files. Whenever possible, the data were processed locally on two servers located at Cambridge, UK, and Philadelphia, USA. We strived to use the most up-to-date MRI image processing time line available at the time. Until 2023, the most recent FreeSurfer version at the time was used. When synthseg became available in 2023, we increasingly used synthseg processing. We were able to include a large number of studies originally pre-processed for a previous publication [15] using FreeSurfer 6.0.142. However, [15] only aggregated the phenotypes used in their study, such that here, we went back to the original freesurfer output files where available to extract subcortical data from the *aseg.stats* files. We further added a number of new studies not included in the prior publication.

Until 2023, for newly added data, we used the most recent FreeSurfer version available, version 7.2. Wherever T1- and T2/FLAIR-weighted raw images were available, these data were processed with FreeSurfer's combined T1-T2 recon-all pipeline. If only raw T1-weighted data were available, and subjects were aged over 2 years, the data were processed with a FreeSurfer standard recon-all pipeline. From 2023 onwards we used synthseg-robust processing. Lastly, if subjects were aged 0–2 years, regardless of of when we received data, we aimed to process it with synthseg for segmentation and grey matter volume extraction, and reconall-clinical for extraction of cortical thickness and surface area values. The exception are the dHCP-fetal and dHCP datasets. Processing details for those datasets can the found elsewhere [79, 121], but briefly, we used the preprocessed version of the data supplied by the dHCP/dHCP-fetal consortia. We aligned the surfaces into dHCP surface template space (`https://github.com/ecr05/dHCP_template_alignment`) and parcellated them using the M-CRIB neonatal cortical and subcortical brain atlas [6]. An overview of processing versions used for each dataset (and site) is provided in **Supplementary Table 1**.

Briefly, the recon-all processing function includes the following steps: non-uniformity correction, projection to Talairach space, intensity normalisation, skull-stripping, automatic tissue and subcortical segmentation, surface interpolation, tessellation and registration.

Regional cortical and subcortical volumes, as well as cortical surface area and thickness defined by the aparc and aseg parcellation templates were extracted.

#### Lifespan trajectories

We used GAMLSS, a robust and flexible framework for modelling non-linear growth trajectories recommended by the World Health Organization [18, 127], to derive developmental curves from the aggregated life-spanning neuroimaging dataset. This modelling strategy allowed us to estimate non-linear heteroskedastic age-related trends (i.e. the mean and variance had non-straight line changes with age) stratified by sex over the entire lifespan. To account for site- or study-specific effects on MRI phenotypes random effects were included within the GAMLSS model. To estimate these models, we used the code published by [15], which is available on github: https://github.com/brainchart/Lifespan

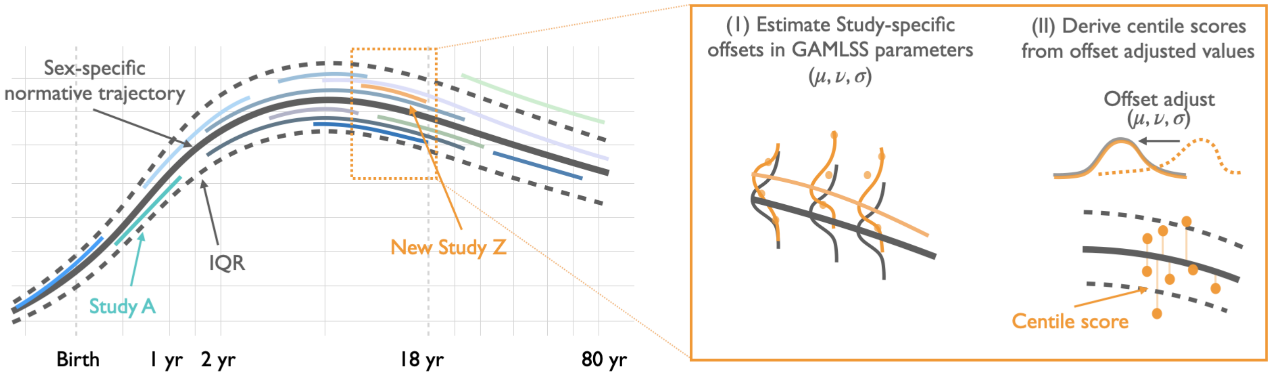

**Fig. 3: Normative modelling:** (*left*) We estimated normative trajectories of various imaging phenotypes as a non-linear function of age, stratified by sex, using GAMLSS models on an aggregated dataset of cross-sectional primary studies. This resulted in sex-specific lifespan trajectories of development of the median of each phenotype. (*right*) We controlled for acquisition site-specific offsets, or random effects, in the first two parameters of the underlying statistical distributions, $\mu$, $\sigma$. After correction for these site-specific offsets, we can derive subject-specific centile scores, which measure an individual's deviation from the benchmark of the normative age- and sex-matched growth chart.

GAMLSS is a class of regression models where all the parameters of the outcome distribution can be modelled as additive functions of the explanatory variables. A strong asset of GAMLSS models is that they allow modeling not only of the central tendency of the outcome distribution ($Y$), but also other parameters of the distribution of $Y$, as linear, nonlinear, parametric, or additive non-parametric functions of explanatory variables and random effects. Prior evidence suggests that brain phenotypes do not only vary in mean, but also in variance across the lifespan. With GAMLSS we chose a modelling framework that could account for variation with age in the first (mean) and second (variance) moment of the outcome distributions. Further, GAMLSS models allow for the distribution of the response variable to be drawn from a general family of distributions which includes, among others, skewed and kurtotic continuous and discrete distributions.

In the GAMLSS framework, the outcome vector $Y$, consisting of independent observations $y_1, y_2, y_3, ...y_N$, follows the probability distribution $F$, with:

$$F \sim F(\mu, \sigma, \nu, \tau) \tag{1}$$

where $F$ is parameterised by typically up to four distribution parameters $(\mu, \sigma, \nu, \tau)$. These parameters can correspond to the mean, variance, skewness, and kurtosis of the outcome distribution, i.e. the first four moments. However, for many distributions the moments of the distribution are a function of these parameters. For example the Beta distribution within GAMLSS is parameterised such that $\mathrm{E}[X_{BE}] = \mu$ (mean) and $\mathrm{V}(X_{BE}) = \sigma^s \mu (1 - \mu)$ (variance) with $0 < (\mu, \sigma) < 1$.

More specifically, each of these components, $k$, is defined in terms of a link-function $g_k$, i.e. a regression on potential covariates. Importantly, the covariates do not have to be the same between the parameters and in fact parameters can reduce to constants, a fact that becomes relevant further below. The link function, $g_k$, thus includes $N_k$ fixed effects, parametrised by their coefficients $\beta_k = (\beta_{k,1}, \beta_{k,2}, ..., \beta_{k,J_k})$, and their design matrix, $X_k$; random effects, $\gamma_k$, with design matrix $Z_k$,; and non-parametric smoothing functions $s_{k,i}$ applied to the $i$th covariate for each parameter, with $i = 1, 2, ...N_k$:

$$g_\mu(\mu) = X_\mu \beta_\mu + Z_\mu \gamma_\mu + \sum_{i=1}^{N_\mu} s_{\mu,i}(x_i) \tag{2}$$

$$g_\sigma(\sigma) = X_\sigma \beta_\sigma + Z_\sigma \gamma_\sigma + \sum_{i=1}^{N_\sigma} s_{\sigma,i}(x_i) \tag{3}$$

$$g_\nu(\nu) = X_\nu \beta_\nu + Z_\nu \gamma_\nu + \sum_{i=1}^{N_\nu} s_{\nu,i}(x_i) \tag{4}$$

$$g_\tau(\tau) = X_\tau \beta_\tau + Z_\tau \gamma_\tau + \sum_{i=1}^{N_\tau} s_{\tau,i}(x_i) \tag{5}$$

The outcome distribution determines the appropiate link function and which parameters (i.e. how many) are modelled; for example the Beta distribution within GAMLSS requires $0 < (\mu, \sigma) < 1$ which naturally leads to a logit-link function for these component regressions. [15] optimised GAMLSS model specification and parameterization to estimate non-linear normative growth trajectories of lifespan brain structural development and determined that the generalized gamma distribution appropriately models their data using a model comparison approach; this is a three-component model $(\mu, \sigma, \tau)$ such that the outcome is positive $(y > 0)$. Here, we use a largely overlapping sample on similar phenotypes, thus we also model a generalized gamma distribution.

We used fractional polynomials to model age-related changes in MRI phentoypes. Using the alternative, non-parametric smoothers (i.e. smoothing splines), would have been more flexible, but also more unstable in terms of model fitting/convergence. Within the GAMLSS framework the appropriate power of the fractional polynomials is chosen in an iterative fitting process across the "standard" set of powers, $p \in \{-2, -1, -0.5, 0, 0.5, 1, 2, 3\}$.

All trajectories fitted here are represented in terms of centile scores rather than the outcome measures directly, or $Z$-scores. The reasoning behind this is that GAMLSS models allow for the outcome distribution to be highly skewed, in which case $Z$-scoring is invalid. For clarity, it is worth emphasizing that while the GAMLSS parameters can map to the moments of the outcome distribution, this depends on the specific distribution that is used. In the case of the generalized gamma distribution used here, the distributional parameters $(\mu, \sigma, \nu)$ do not directly map to the mean, variance and skewness of the distribution. Thus, while as specified in **Equation 4** $\nu$ does not change with age, the skewness of the outcome distribution can. All trajectories shown are the median (50th percentile) of the distribution of normative trajectories.

#### Left/right regional models

We estimated GAMLSS models to derive trajectories of regional brain development. Briefly, in a sex-stratified approach, we estimated lifespan development in the first order $(\mu)$ and second order $(\sigma)$ distributional parameters of brain phenotypes using GAMLSS with fixed effects of age and pre-processing pipeline, and a random effect of site, using fractional polynomials to model non-linear age-related trends.

We modeled the effect of site as a random effect, as opposed to a fixed effect. These random effects are assumed to follow a normal distribution with mean zero and a variance term (recall this is on the link-scale, hence the random effects will likely induce non-normal variability on the outcome-scale). We chose to use the simple case of random intercept, i.e. a group-level intercept. The advantage of this procedure is that we can estimate the (random) effect of a new site, i.e., a site that is not included in the original dataset used to fit the normative growth chart, from the random-effect covariance structure.

The third distributional parameter, $\nu$, was only modeled as a constant, since previous work using GAMLSS models to estimate lifespan trajectories [15], found that including age, sex and study parameters for this term lead to numerical instability. Since we have no *a priori* reason to assume age-dependent or random effects on skewness in subcortical volume, we chose not to model lifespan changes in this parameter. This means that **Equation 4** reduces to:

$$\nu = \alpha_\nu \tag{6}$$

##### Regional asymmetry models

Prior to model-fitting, we performed a family selection to determine the underlying distribution family for the asymmetry models. *A priori*, we decided to evaluate three parameter distributions, with a range from $-\infty$ to $\infty$, since we knew from prior work, that it was unlikely four parameter distributions would fit on neuroimaging data [15]. We thus fit models using three distributions included in the gamlss package: SN1, SN2 and exGAUS. For simplicity, we fit models to the 'global' phenotypes of hemispheric asymmetry in grey matter volume, cortical thickness and surface area, using random effect ofsite. We then compared the AIC of the highest order model we could fit for each phenotype and condition. Thus for all regional asymmetry phenotypes, we used SN2 as the underlying distribution.

For each bilateral region $i$, we estimated the regional asymmetry in cortical and subcortical volumes, as well as cortical surface area and thickness, as the ratio of the difference in left $(L_i)$ and right $(R_i)$ regional values, over the sum of them:

$$A_i = (L_i - R_i)/(L_i + R_i) \tag{7}$$

Given that both regional values are necessarily always positive, $A_i$ is bounded between $-1$ and $1$.
For modelling purposes, we transform $A_i$ to range from $-\infty$ to $\infty$ using:

$$tA_i = tan(\frac{A_i * \pi}{2}) \tag{8}$$

As an aside, asymmetry is a $(-1, 1)$ bounded outcome and it was also possible to model directly using a scaled-and-shifted Beta distribution (among other bounded distribution options within the gamlss-package) or to consider truncated distributions. Truncated distributions were deemed less directly interpretable and they can exhibit strange edge effects when fitting; although a normal truncated to $(-1, 1)$ would be valid transforming the assymetry measure to the real line is less complex. Equally a transformation to $(-1, 1)$ is equivalent to $(-\infty, \infty)$, the latter being more familiar and using a more familiar normal distribution was deemed more appropriate.

##### Developmental milestones

We estimated the age at which three key developmental milestones were reached in each of the modelled trajectories. The three milestones were: (i) The peak rate-of-growth, estimated as the peak of the first derivative of each normative median trajectory; (ii) the peak of the trajectory, determined as the peak of the median trajectory of each regional phenotype; and (iii) the peak rate-of-decline, was estimated as the trough of the first derivative of each normative median trajectory.

We further estimated the hemispheric difference $(\Delta M)$ in the age at which the milestones were reached. We determined the significance of the difference in a bootstrap approach as follows: We resampled with replacement the data (see details in section **Bootstrap analyses**). We re-estimated the milestones in each bootstrap for the left and right phenotype. We then estimated the difference in those phenotypes in each bootstrap iteration. We then estimated the standard deviation of the difference across bootstraps. We then determined the significance of the hemispheric difference in milestones based on whether the interval $[\Delta M - 1.96 * SD, \Delta M + 1.96 * SD]$ included $0$.

##### Centile score estimation

We obtained the relative distance of each individual's observation from the normative trajectory of each brain phenotype, as the relative distance from the median of the age-normed distributions of the reference model, stratified by sex. This distance is termed "centile" and describes the percentage rank of the individual's brain phenotype measurement benchmarked by the normative distribution of the corresponding phenotype. More specifically, we derived the study-specific centile $q_i$ for an individual's observation, $i$, as:

$$q_i = F'(y, x|\beta, z) \tag{9}$$

where $F'$ is the inverse cumulative density function of (**Equation 1**) of a brain phenotype, $\beta$ are the coefficients of the fixed effects, $z$ is the random effect of study, $x$ are the individual's covariates, and $y$ is the outcome measure, i.e., the brain MRI phenotype.

| dx | N | female | male |
|---|---|---|---|
| Autism Spectrum Disorder (ASD) | 2421 | 464 | 1957 |
| Attention Deficit and Hyperactivity Disorder (ADHD) | 1795 | 520 | 1274 |
| Major Depressive Disorder (MDD) | 8212 | 5379 | 2833 |
| Schizophrenia (SCZ) | 1479 | 589 | 890 |
| Mild Cognitive Impariment (MCI) | 1535 | 758 | 777 |
| Alzheimer's Disease (AD) | 3070 | 1667 | 1403 |
| Other disorder | 13799 | 7206 | 6592 |

**Table 2: Subjects with psychopathology:** Our dataset includes data from subjects with dozens of disorders, including ASD, ADHD, MDD, SCZ, MCI, AD and others.

##### Case control differences

It is worth reiterating that the normative model is base on healthy controls and, as such, being in the $1^{st}$ of $99^{th}$ is not indicative of any clinical issue (by definition there are people in the top 1% and bottom 1% of a healthy population).

Using the centile scores derived above, we estimated deviations from normative development in normative controls in multiple psychiatric and developmental disorders.

To this end, the aggregated dataset included a total of 2421 subjects autism spectrum disorder, 1479 subjects with schizophernia, and 3070 with Alzheimer's disease (**Table 1**).

We estimated differences in centile scores between healthy controls and patients using a bootstrapped (500 bootstraps) non-parametric generalization of Welch's one-way ANOVA.

We then conducted post-hoc comparisons for all case-control combinations using a non-parametric Monte Carlo permutation test with 10,000 permutations. The results were corrected for multiple comparisons using FDR correction. Lastly, we estimated effect sizes of the case-control differences using Cohen's $d$.

#### Quality control

##### Automated quality control

Given the very large number of scans in this aggregated dataset, we did not perform manual quality control of each individual scan. Rather, we made use of the Synthseg-generated quality control scores for synthseg-processed scans, and we estimated the Euler Index (EI) for each freesurfer-processed scan as a measure of image quality. The EI is an automated, quantitative measure of data quality in scans processed by FreeSurfer and as such is only available for FreeSurfer processed data. It measures the number of "surface holes", or topological defects, in the cortical surface reconstruction, across hemispheres prior to correction. More specifically, surface holes are regions on the surface mesh where there are missing or disconnected vertices, resulting in gaps in the surface for example as a result of image artifacts, or partial volume effects. This work focuses on volumetric measures which are likely less affected by surface holes, however the EI has previously been used as a general measure of raw scan quality, which is how we have used it here. Previous work has provided evidence that there likely is no single EI threshold that is generalizable as a valid criterion of image quality across studies [111].

Prior work has shown robustness of braincharts morphometric centile scores to image quality [15]. Here, we assess the robustness of asymmetry centile scores to image quality, using the example of grey matter volume.

First, we assessed the potential relationship between age and image quality. Specifically, in a sex-stratified approach, we estimated the linear effect of age on QC measures using linear mixed effects models with a random effect of site:

$$\mathrm{QC} \sim 1 + \beta_{age} * age + \beta_{sex} * sex + \gamma_{site} * (1|site) + \epsilon \quad (10)$$

where QC is the minimum synthseg QC score or the Euler index for a given subject, $\beta$ refers to the coefficients for the fixed effects, $\gamma_{subject}$ refers to the coefficients for random effects, and $\epsilon$ represents the residual error.

We find that there indeed is a significant effect of age on QC metric, such that younger subjects tend to have lower image quality (SynthSeg: $t = 8.7, P < 0.001$; Freesurfer: $t = 49; P < 0.001$). Young subjects were processed using SynthSeg, such that below we illustrate the effect of age on the SynthSeg QC metric (**SI Fig. 4**).

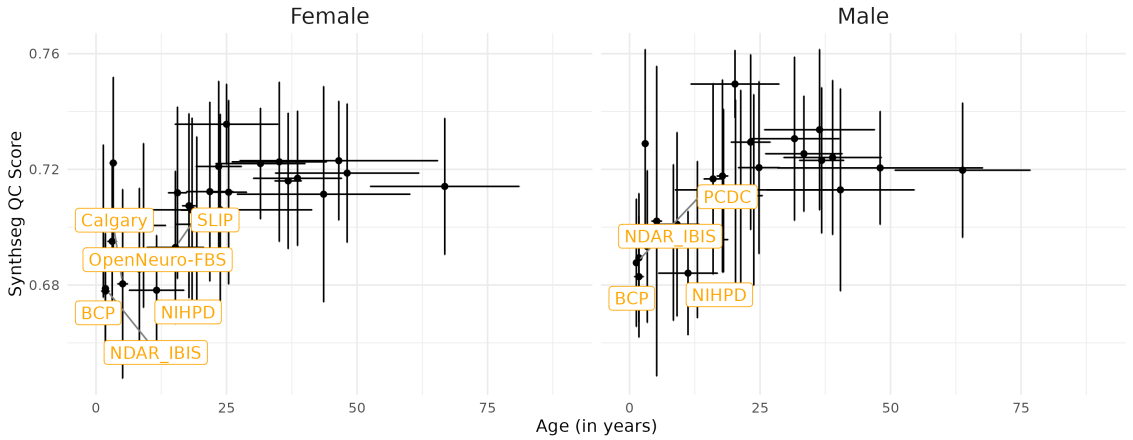

**Fig. 4: Effect of age on SynthSeg QC metrics:**

Secondly, we estimated the effects of image quality on model-derived centile scores. In a sex-stratified approach, for each site, we estimated the Spearman correlation between the subject-specific centile scores and the QC measure. We find that the magnitude of the relationship between QC scores and centile scores was negligable for both Freesurfer and synthseg processed scans, however correlations were sometimes significant, presumably due to the large saple size (**SI Table 4-3**).

##### Bootstrap analyses

In an effort to evaluate the reliability and stability of the derived lifespan trajectories, we estimated confidence intervals around all parameters by a bootstrap procedure. More specifically, we randomly resampled with replacement 1,000 times and re-estimated the lifespan trajectories for each regional phenotype. Each of these bootstrap iterations was restrained to ensure a random dataset that was comparable to our original dataset in terms of the sex distribution, the age distribution, and the relative size of the original primary studies. Retaining the relative proportions of males and females in the resampled dataset is relevant, since we stratified our original models by sex, i.e. we included sex as a fixed effect. Our original models included a random effect of study. In order to derive bootstrap confidence intervals around this study parameter, we constrained the resampling to retain the original age distribution and study size. Failing to constrain the resampling to consider study size could lead to individual studies being omitted in some iterations, and otherwise skew the bootstrap value, leading to inappropriate comparisons between original and bootstrap values.

##### Quantification of asymmetry magnitude

We summarized the magnitude of asymmetry in each region across (i) the entire lifespan, and (ii) developmental periods more specifically by estimating the area under the curve of the asymmetry trajectories. More specifically, in R, we used the `stats` package (Version 3.6.2), to first interpolate the asymmetry index as a function of age using `aproxfun`. We then estimated the (signed) area under the curve of this function using the `integrate` function. In order to obtain the absolute (unsigned) area of the curve, we additionally segmented the asymmetry curves into segments whenever the median trajectory crosses zero.

We applied this approach (i) across the entire lifespan, resulting in an estimate of overall/lifespan magnitude of asymmetry in a given region (**Main Fig 2B**); and (ii) in each prior developmental window [64], resulting in an estimate of asymmetry magnitude in relevant periods of brain development (**Fig. 3**). Since the developmental windows have vastly different lengths, we adjusted the resulting integral by the length of the development window using the log-transformed number of days.

a

#### GWAS analyses of centile scores

Variation in human brain anatomy is partially genetic and prior work has identified a number of genetic loci associated with multiple (bilaterally averaged, raw) imaging phenotypes [37, 148, 55]. Here, we aimed to assess the genetic architecture of individual deviations from normative brain development. To this end, we conducted genome-wide association analyses of centile scores for both the left/right regional models, as well as asymmetry centile scores across four phenotypes: grey matter volume, surface area, cortical thickness and subcortical structures. We conducted our analyses in the UK Biobank Sample [25].

| regions | pval_Female | pval_Male | corval_Female | corval_Male |
|---|---|---|---|---|
| bankssts | 0.73 | 0.21 | -0.00 | 0.01 |
| caudalanteriorcingulate | 0.38 | 0.16 | -0.00 | 0.01 |
| caudalmiddlefrontal | 0.26 | 0.74 | 0.00 | 0.00 |
| cuneus | 0.01 | 0.11 | -0.01 | -0.01 |
| entorhinal | 0.00 | 0.19 | -0.02 | -0.01 |
| frontalpole | 0.02 | 0.00 | -0.01 | -0.01 |
| fusiform | 0.01 | 0.00 | 0.01 | 0.02 |
| inferiorparietal | 0.00 | 0.00 | 0.02 | 0.02 |
| inferiortemporal | 0.00 | 0.11 | -0.01 | -0.01 |
| insula | 0.00 | 0.00 | 0.07 | 0.08 |
| isthmuscingulate | 0.18 | 0.87 | 0.01 | -0.00 |
| lateraloccipital | 0.51 | 0.10 | -0.00 | 0.01 |
| lateralorbitofrontal | 0.01 | 0.01 | -0.01 | -0.01 |
| lingual | 0.00 | 0.04 | -0.01 | -0.01 |
| medialorbitofrontal | 0.00 | 0.00 | -0.03 | -0.03 |
| middletemporal | 0.00 | 0.00 | -0.03 | -0.04 |
| paracentral | 0.74 | 0.11 | 0.00 | 0.01 |
| parahippocampal | 0.24 | 0.64 | 0.00 | 0.00 |
| parsopercularis | 0.06 | 0.02 | -0.01 | -0.01 |
| parsorbitalis | 0.00 | 0.12 | -0.02 | -0.01 |
| parstriangularis | 0.00 | 0.00 | -0.02 | -0.02 |
| pericalcarine | 0.18 | 0.72 | 0.01 | -0.00 |
| postcentral | 0.55 | 0.02 | -0.00 | -0.01 |
| posteriorcingulate | 0.88 | 0.00 | 0.00 | 0.02 |
| precentral | 0.77 | 0.22 | 0.00 | -0.01 |
| precuneus | 0.01 | 0.96 | -0.01 | 0.00 |
| rostralanteriorcingulate | 0.00 | 0.00 | 0.02 | 0.02 |
| rostralmiddlefrontal | 0.00 | 0.09 | -0.02 | -0.01 |
| superiorfrontal | 0.33 | 0.47 | -0.00 | -0.00 |
| superiorparietal | 0.00 | 0.00 | 0.02 | 0.03 |
| superiortemporal | 0.00 | 0.00 | -0.02 | -0.01 |
| supramarginal | 0.00 | 0.00 | 0.01 | 0.02 |
| temporalpole | 0.29 | 0.00 | 0.00 | 0.02 |
| transversetemporal | 0.16 | 0.86 | -0.01 | 0.00 |

**Table 3: Effect of synthseg QC scores on asymmetry GM centile scores:**

| regions | pval_Female | pval_Male | corval_Female | corval_Male |
|---|---|---|---|---|
| bankssts | 0.73 | 0.21 | -0.00 | 0.01 |
| caudalanteriorcingulate | 0.38 | 0.16 | -0.00 | 0.01 |
| caudalmiddlefrontal | 0.26 | 0.74 | 0.00 | 0.00 |
| cuneus | 0.01 | 0.11 | -0.01 | -0.01 |
| entorhinal | 0.00 | 0.19 | -0.02 | -0.01 |
| frontalpole | 0.02 | 0.00 | -0.01 | -0.01 |
| fusiform | 0.01 | 0.00 | 0.01 | 0.02 |
| inferiorparietal | 0.00 | 0.00 | 0.02 | 0.02 |
| inferiortemporal | 0.00 | 0.11 | -0.01 | -0.01 |
| insula | 0.00 | 0.00 | 0.07 | 0.08 |
| isthmuscingulate | 0.18 | 0.87 | 0.01 | -0.00 |
| lateraloccipital | 0.51 | 0.10 | -0.00 | 0.01 |
| lateralorbitofrontal | 0.01 | 0.01 | -0.01 | -0.01 |
| lingual | 0.00 | 0.04 | -0.01 | -0.01 |
| medialorbitofrontal | 0.00 | 0.00 | -0.03 | -0.03 |
| middletemporal | 0.00 | 0.00 | -0.03 | -0.04 |
| paracentral | 0.74 | 0.11 | 0.00 | 0.01 |
| parahippocampal | 0.24 | 0.64 | 0.00 | 0.00 |
| parsopercularis | 0.06 | 0.02 | -0.01 | -0.01 |
| parsorbitalis | 0.00 | 0.12 | -0.02 | -0.01 |
| parstriangularis | 0.00 | 0.00 | -0.02 | -0.02 |
| pericalcarine | 0.18 | 0.72 | 0.01 | -0.00 |
| postcentral | 0.55 | 0.02 | -0.00 | -0.01 |
| posteriorcingulate | 0.88 | 0.00 | 0.00 | 0.02 |
| precentral | 0.77 | 0.22 | 0.00 | -0.01 |
| precuneus | 0.01 | 0.96 | -0.01 | 0.00 |
| rostralanteriorcingulate | 0.00 | 0.00 | 0.02 | 0.02 |
| rostralmiddlefrontal | 0.00 | 0.09 | -0.02 | -0.01 |
| superiorfrontal | 0.33 | 0.47 | -0.00 | -0.00 |
| superiorparietal | 0.00 | 0.00 | 0.02 | 0.03 |
| superiortemporal | 0.00 | 0.00 | -0.02 | -0.01 |
| supramarginal | 0.00 | 0.00 | 0.01 | 0.02 |
| temporalpole | 0.29 | 0.00 | 0.00 | 0.02 |
| transversetemporal | 0.16 | 0.86 | -0.01 | 0.00 |

**Table 4: Effect of synthseg Euler index on asymmetry GM centile scores:**

We used the genotype data provided by the UK Biobank following centralized quality control and imputation procedures [25]. All analyses were restricted to participants who self-identified as of European ancestry to minimize population stratification effects. We applied additional subject-level quality control by excluding individuals who were more than five standard deviations from the mean of the first two genetic principal components, those with a genotyping rate below 95%, individuals with excessive heterozygosity, and those whose genetic sex did not match their self-reported sex.

At the variant level, we applied standard quality control thresholds: Analyses were restricted to variants with a minor allele frequency (MAF) of at least 0.1% MAF. We excluded variants with genotype call rates below 95% or that deviated from Hardy-Weinberg equilibrium in the European sample ($P < 110^{-6}$). For imputed variants, we required an imputation quality score (INFO or $R^2$) greater than 0.4 to ensure reliability of genotype dosage estimates.

First, we derived centile scores for left/right regional and asymmetry trajectories for each UKB subject. We then conducted GWAS analyses of these centile scores for all phenotypes using FastGWA (v1.93) [62] with the following covariates: age, age$^2$, sex, age $\times$ sex, age$^2$ $\times$ sex, imaging center, first 40 genetic principal components, mean framewise displacement (as obtained from the accompanying resting-state fMRI scan), maximum framewise displacement (as obtained from the accompanying resting-state fMRI scan) and Euler Index [111] as covariates. We next computed pairwise genetic correlations for each left-right combination of regions within a phenotype, as well as SNP heritability for the GWAS statistics using LDSC (v1.01) [23, 24], using LD weights from the North West European populations. Lastly, we estimated the heritability of asymmetry centile scores using LDSC (v1.01) [23, 24].

### Supplementary results

#### Regional trajectories of brain development

In the following pages we visually display the left and right regional trajectories of brain development for grey matter volume (GM), surface area (SA), cortical thickness (CT) and subcortical structures (SUBC).

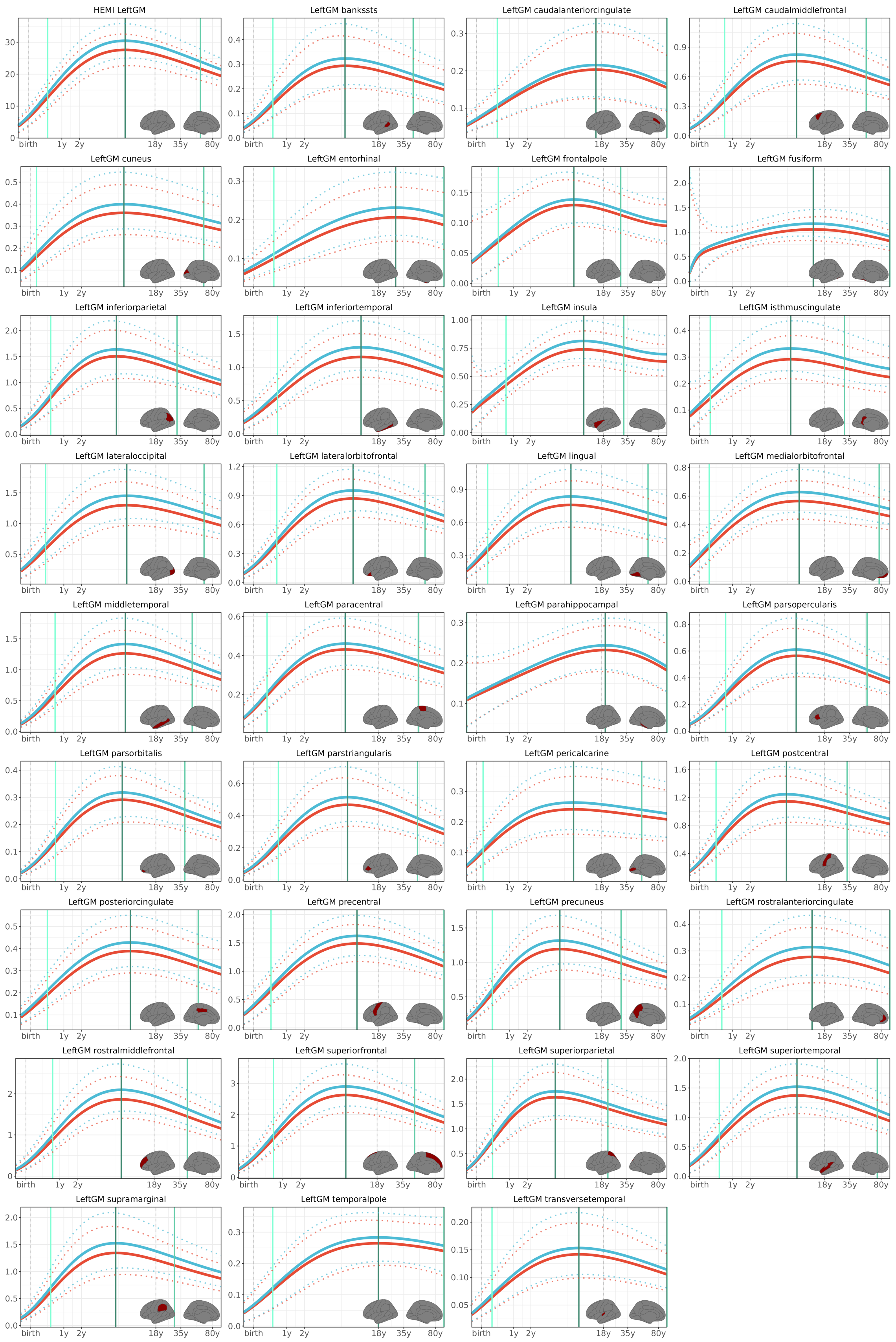

**Fig. 5: Regional left grey matter volume trajectories:** Female (red) and male (turquoise) lifespan trajectories of development. We denote three milestones of development: the peak rate of growth (light turquoise), the peak (dark green), and the peak rate of decline (light green).

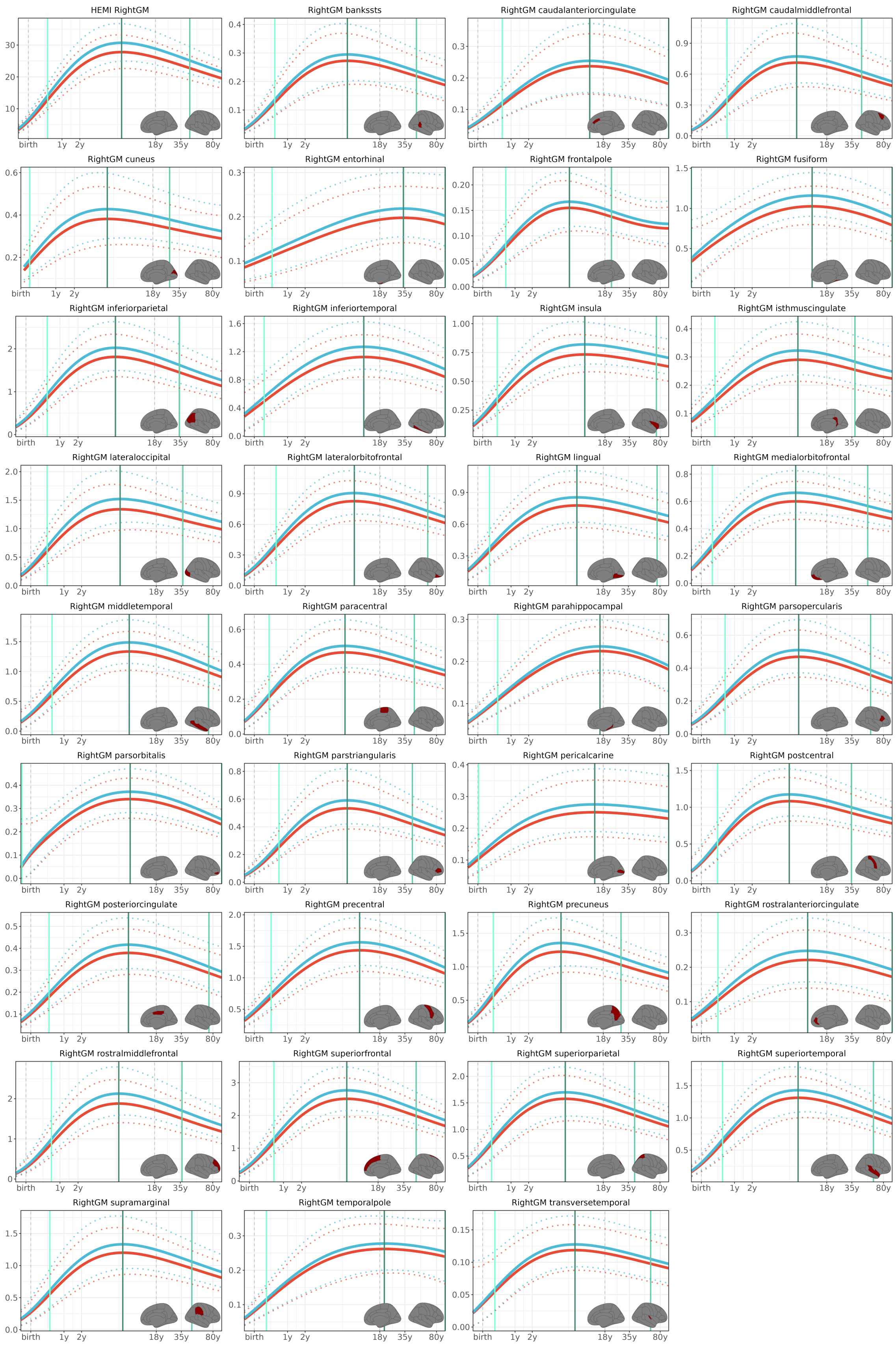

**Fig. 6: Regional right grey matter volume trajectories:** Female (red) and male (turquoise) lifespan trajectories of development. We denote three milestones of development: the peak rate of growth (light turquoise), the peak (dark green), and the peak rate of decline (light green).

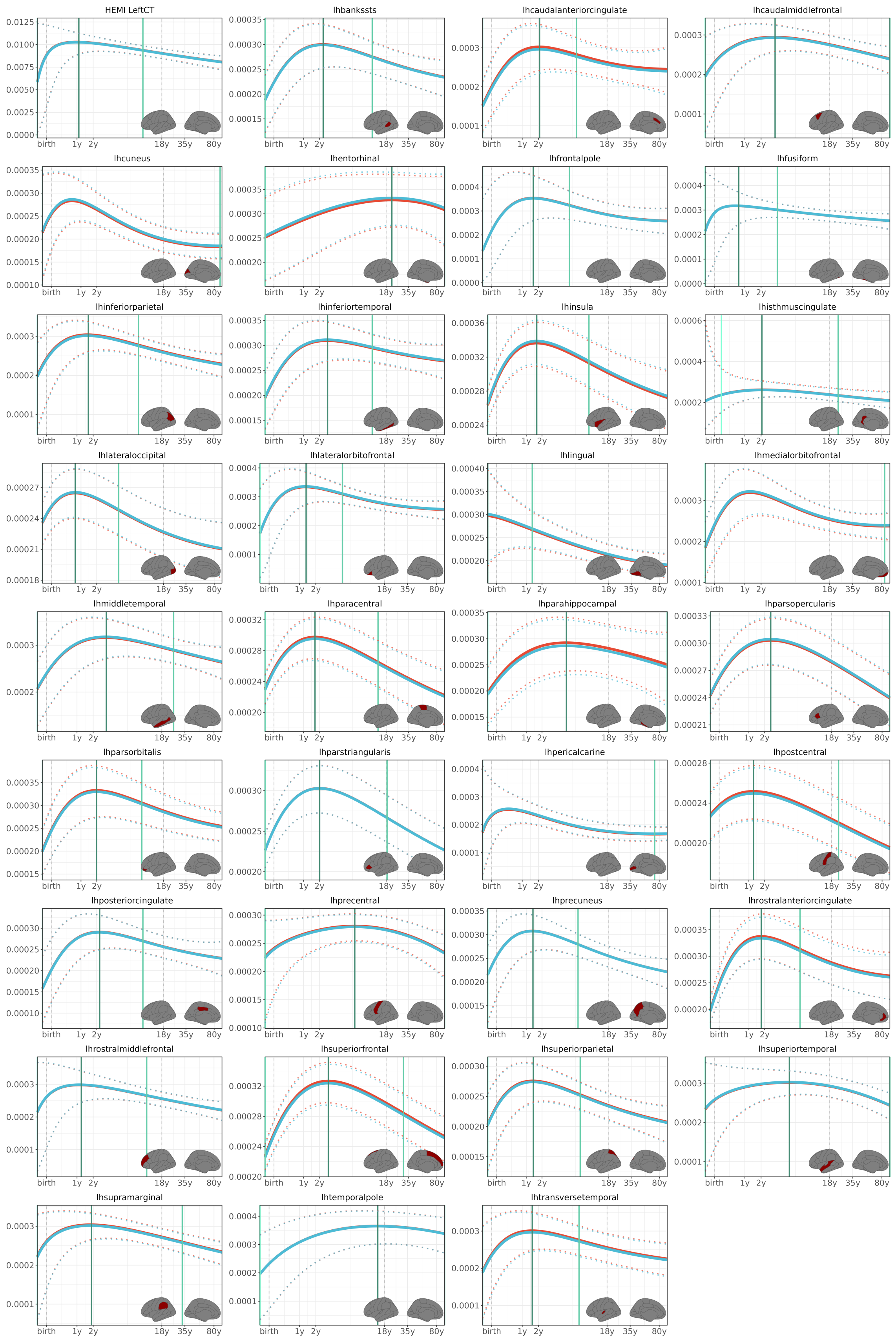

**Fig. 7: Regional left cortical thickness trajectories:** Female (red) and male (turquoise) lifespan trajectories of development. We denote three milestones of development: the peak rate of growth (light turquoise), the peak (dark green), and the peak rate of decline (light green).

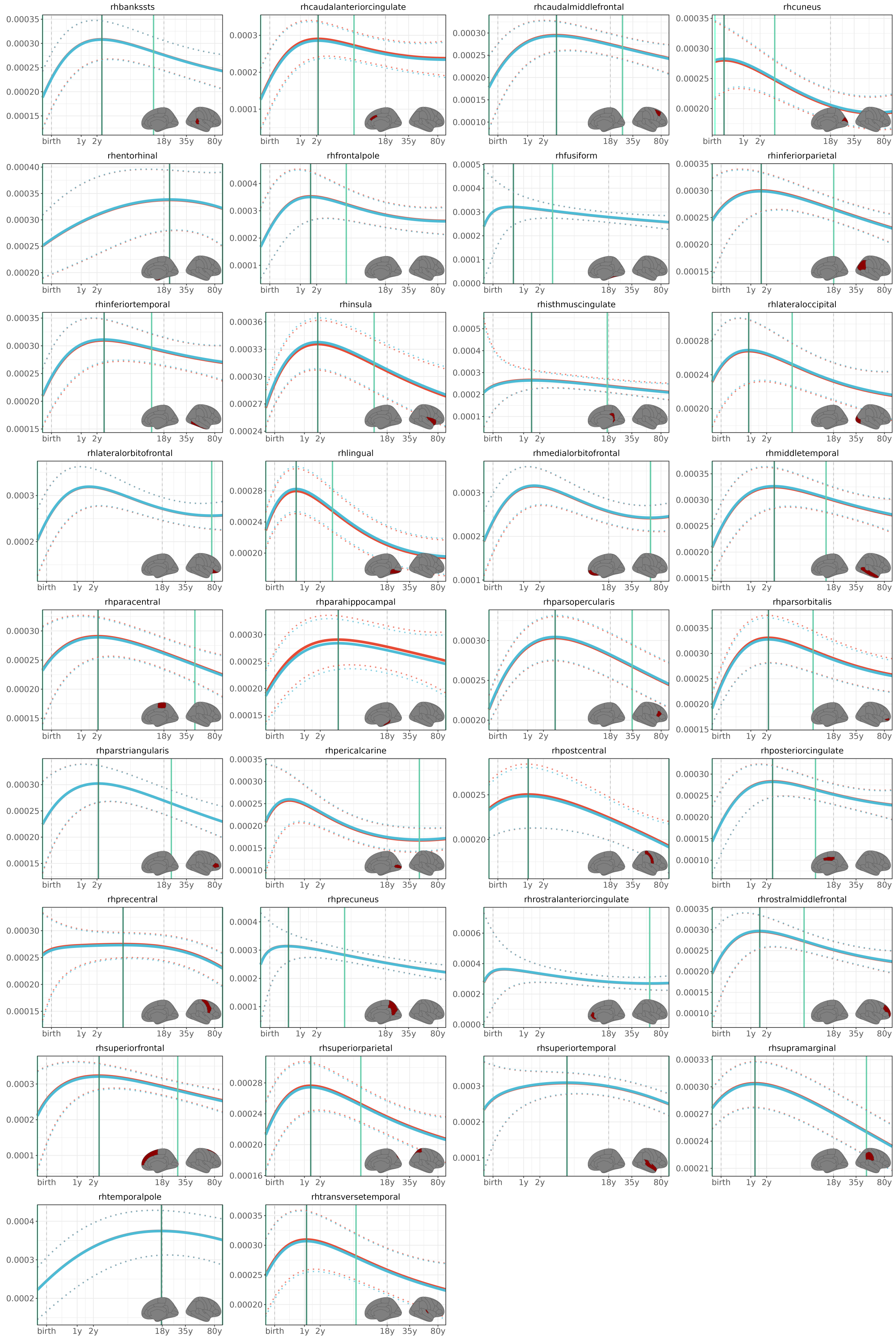

**Fig. 8: Regional right cortical thickness trajectories:** Female (red) and male (turquoise) lifespan trajectories of development. We denote three milestones of development: the peak rate of growth (light turquoise), the peak (dark green), and the peak rate of decline (light green).

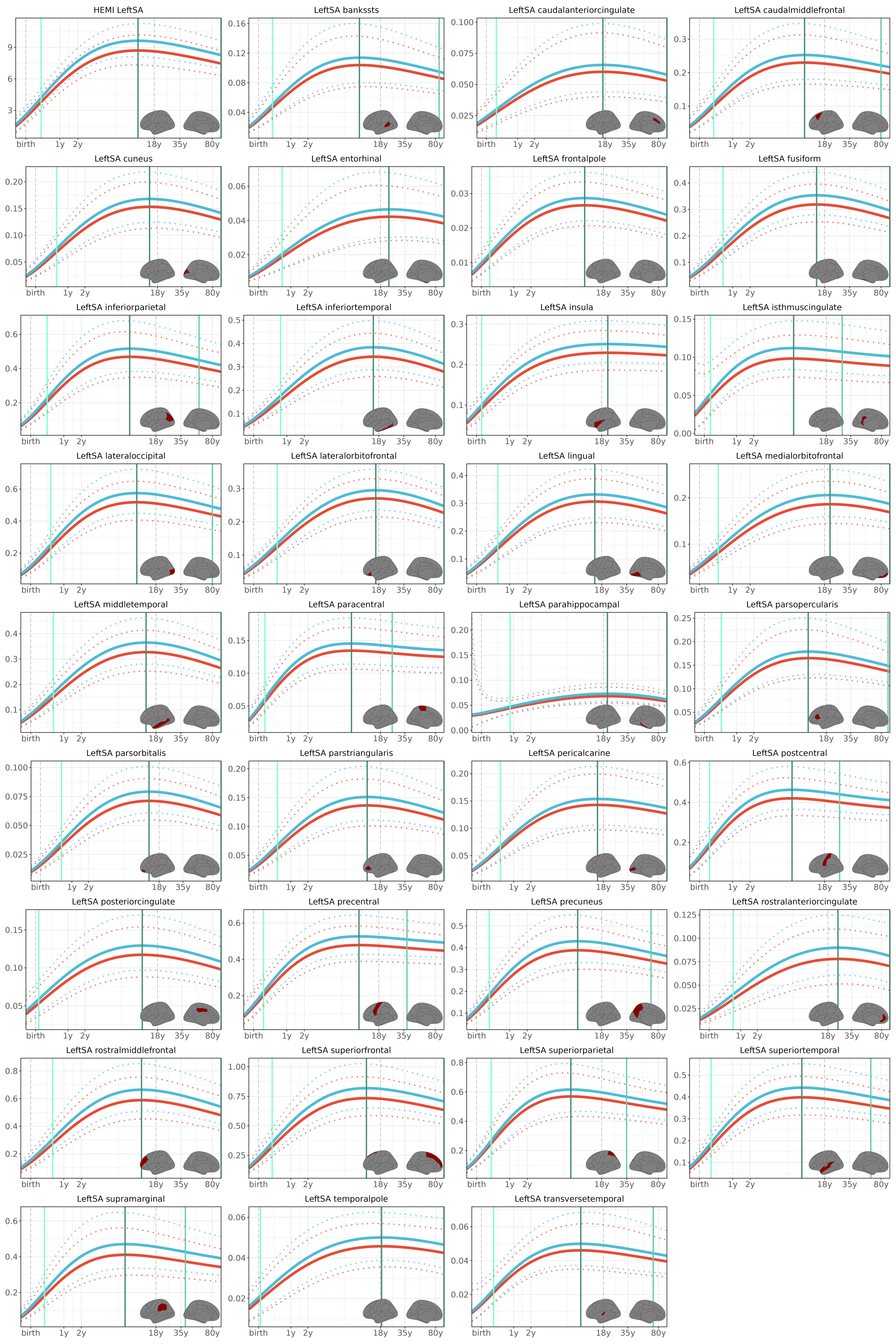

**Fig. 9: Regional left surface area trajectories:** Female (red) and male (turquoise) lifespan trajectories of development. We denote three milestones of development: the peak rate of growth (light turquoise), the peak (dark green), and the peak rate of decline (light green).

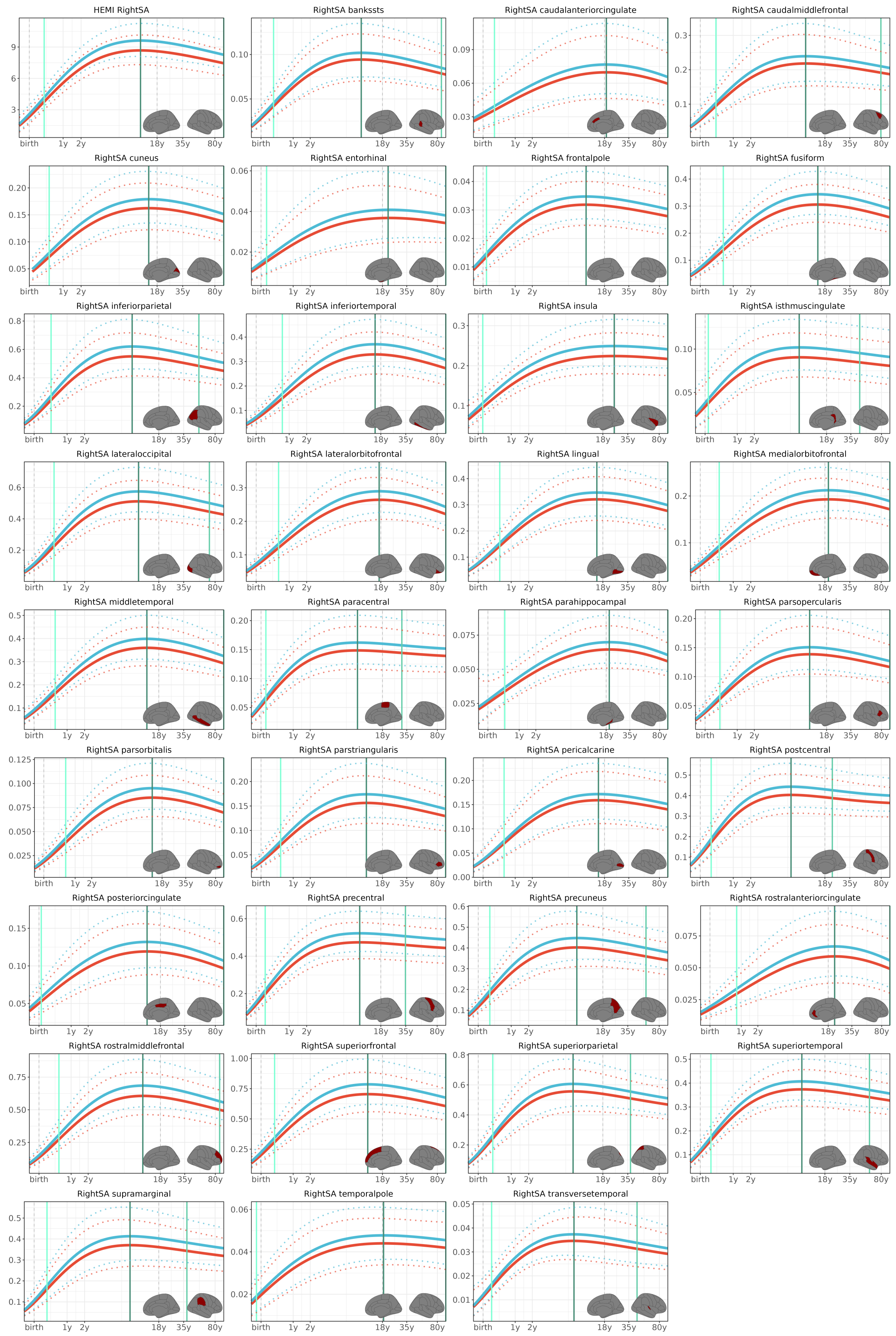

**Fig. 10: Regional right surface area trajectories:** Female (red) and male (turquoise) lifespan trajectories of development. We denote three milestones of development: the peak rate of growth (light turquoise), the peak (dark green), and the peak rate of decline (light green).

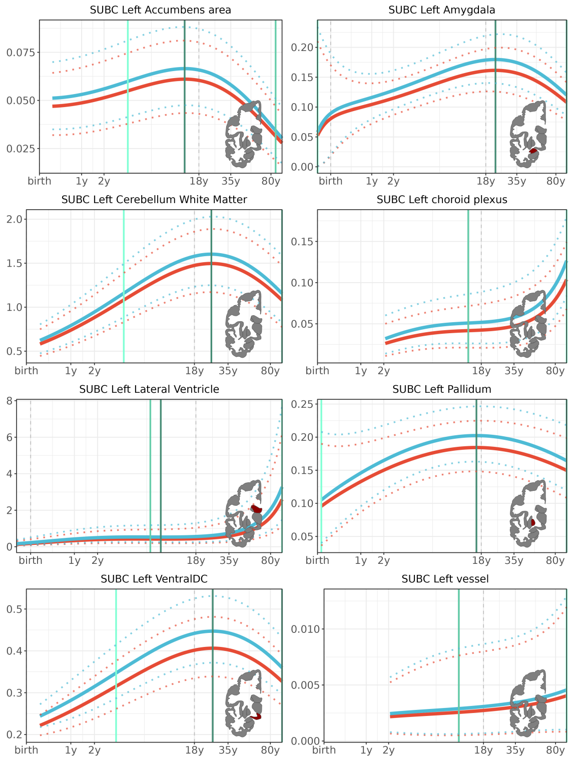

**Fig. 11: Regional left subcortical trajectories:** Female (red) and male (turquoise) lifespan trajectories of development. We denote three milestones of development: the peak rate of growth (light turquoise), the peak (dark green), and the peak rate of decline (light green).

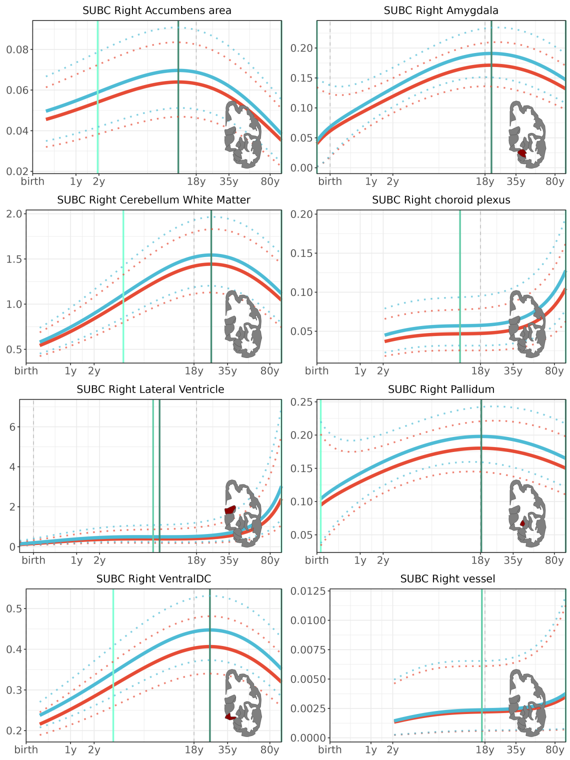

**Fig. 12: Regional right subcortical trajectories:** Female (red) and male (turquoise) lifespan trajectories of development. We denote three milestones of development: the peak rate of growth (light turquoise), the peak (dark green), and the peak rate of decline (light green).

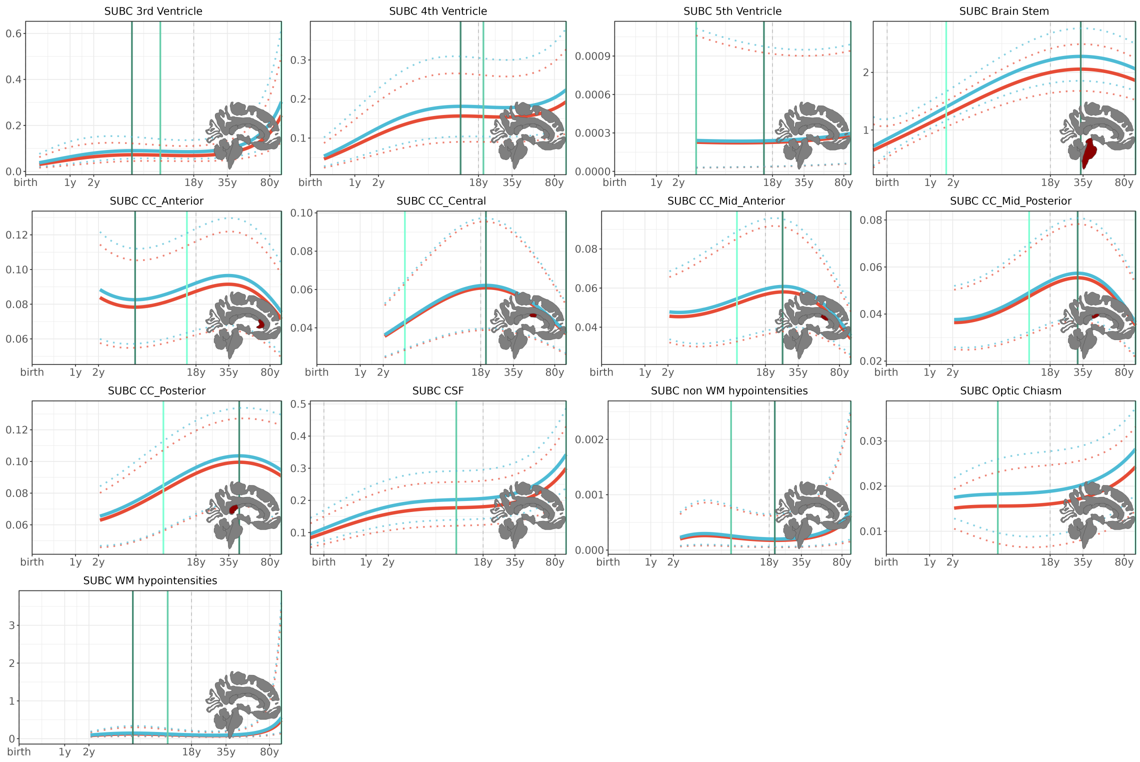

**Fig. 13: Regional midline subcortical trajectories:** Female (red) and male (turquoise) lifespan trajectories of development. We denote three milestones of development: the peak rate of growth (light turquoise), the peak (dark green), and the peak rate of decline (light green).

#### Developmental milestones

We characterized the lifespan trajectories of left and right regional structural phenotypes in terms of three key develpmental milestones: (i) the age at peak rate-of-growth, (ii) the age at peak, an (iii) the age at peak rate-of-decline. The results of the first two milestones are in line with previous bilaterally averaged findings [15]. Additionally, we report the peak rate of decline. We find that subcortical structures tend decline most strongly in the very end of life (here denoted as the last timepoint we map in our trajectories). Many surface area regions peak in their rate of decline around the same time, whereas more varied results are observed in grey matter volume and cortical thickness.

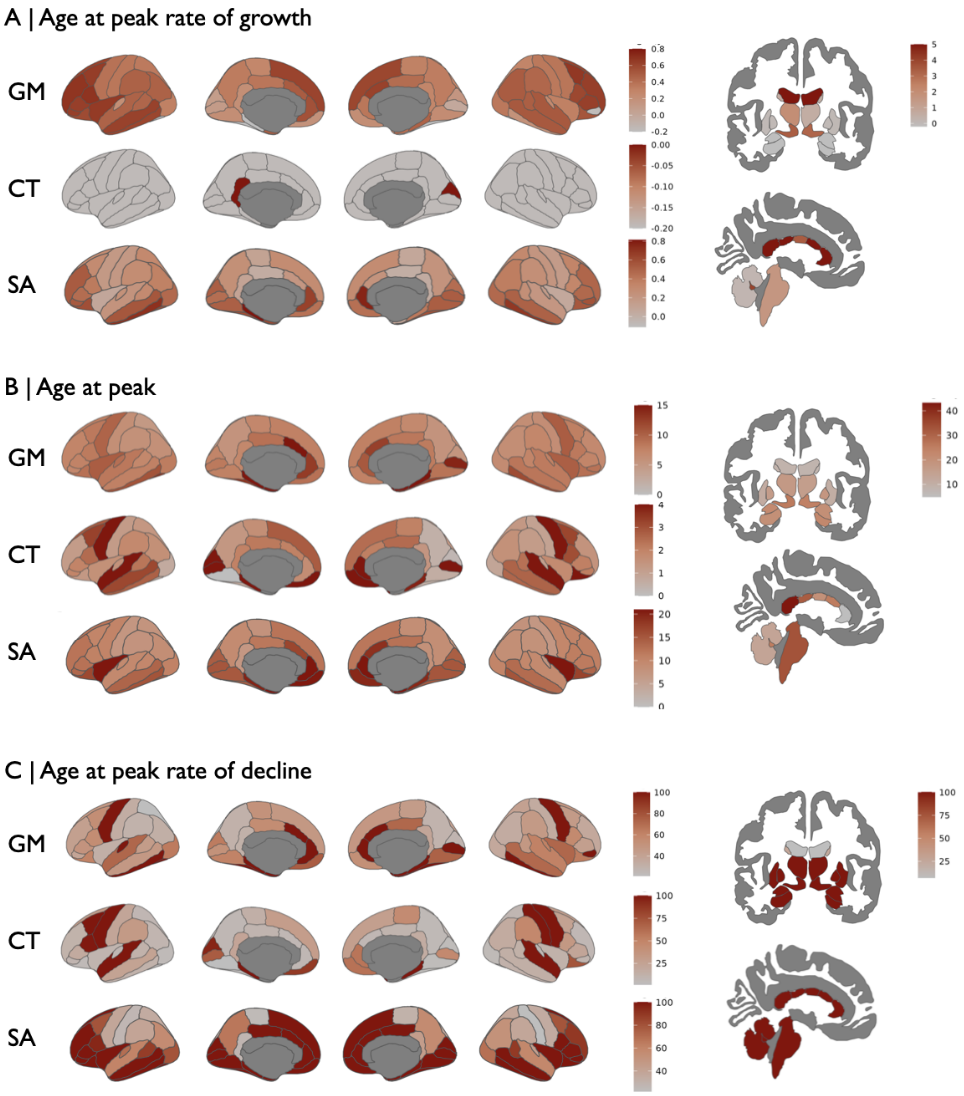

**Fig. 14: Developmental milestones:** We estimate the milestones of development in grey matter volume, cortical thickness, surface area and subcortical structures. (A) Maps of the age at peak rate-of-growth in each phenotype. (B) Maps of the age at peak grey matter volume, surface area, cortical thickness and subcortical tissue volume. (C) Maps of the age at peak rate-of-decline.

Next, we estimated the hemispheric difference in each regional milestones for each phenotype. We the determined significance of the difference using a bootstrap approach (see **Supplementary Methods**) We observe hemispheric differences in the peak of up to X years. Since most cortical thickness trajectories display their peak of growth at the first timepoint we can map, there are very few observable hemisperic differences in the peak rate of growth for this phenotype. In a similar fashion, almost all subcortical volume trajectories have their peak rate of decline at the last timepoint we map, leading to very few hemispheric differences in this phenotype.

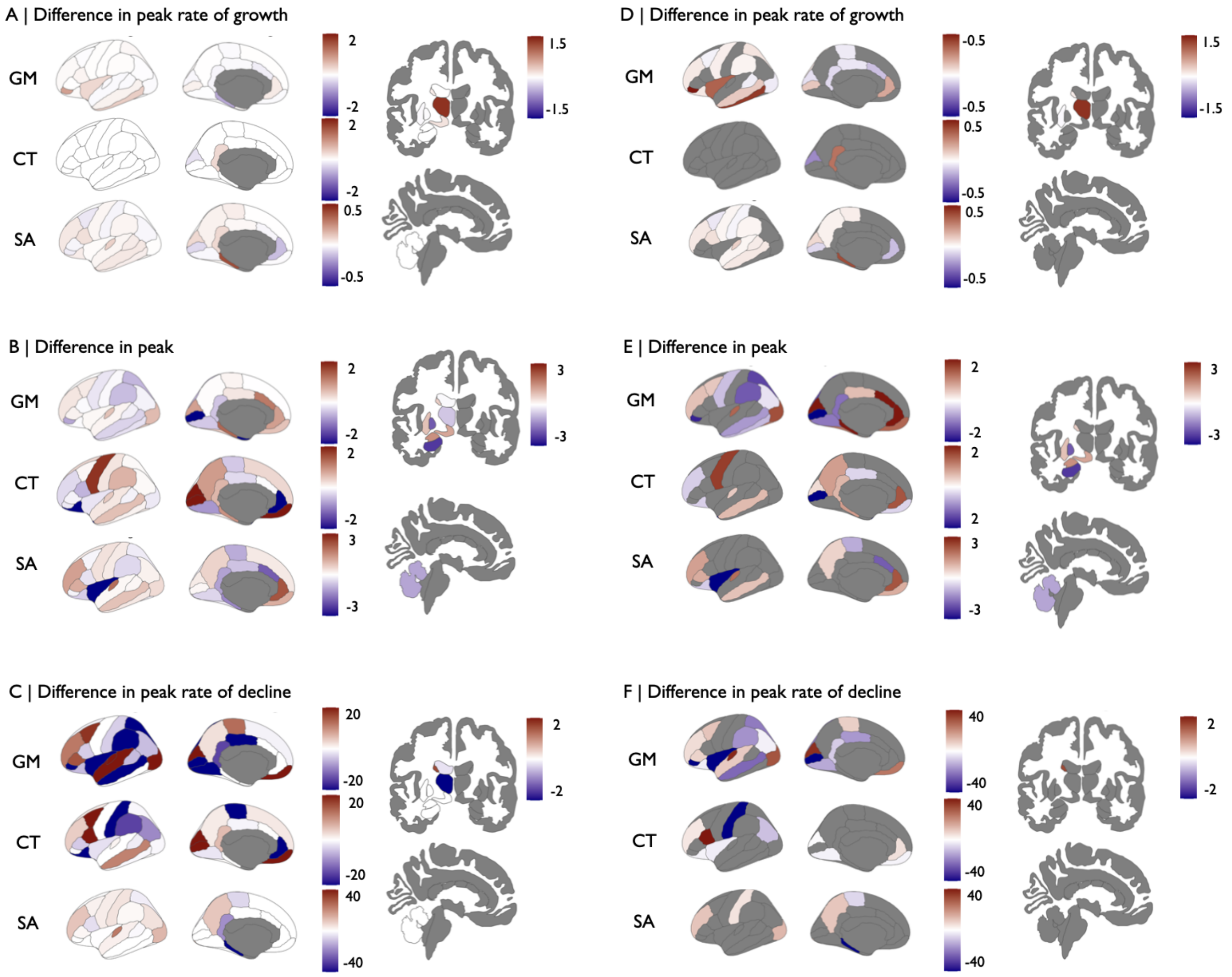

**Fig. 15: Hemispheric differences in developmental milestones:** We estimate the hemispheric difference in developmental milestones, i.e. the age difference in the milestone for all phenotypes. (A) Maps the left-right hemispheric difference in the age at peak rate-of-change, peak, and peak rate-of-decline in each phenotype. (B) We estimated the signifiance of the left-right difference in a bootstrap approach. Here, we shop the thresholded maps ($P < 0.05$).

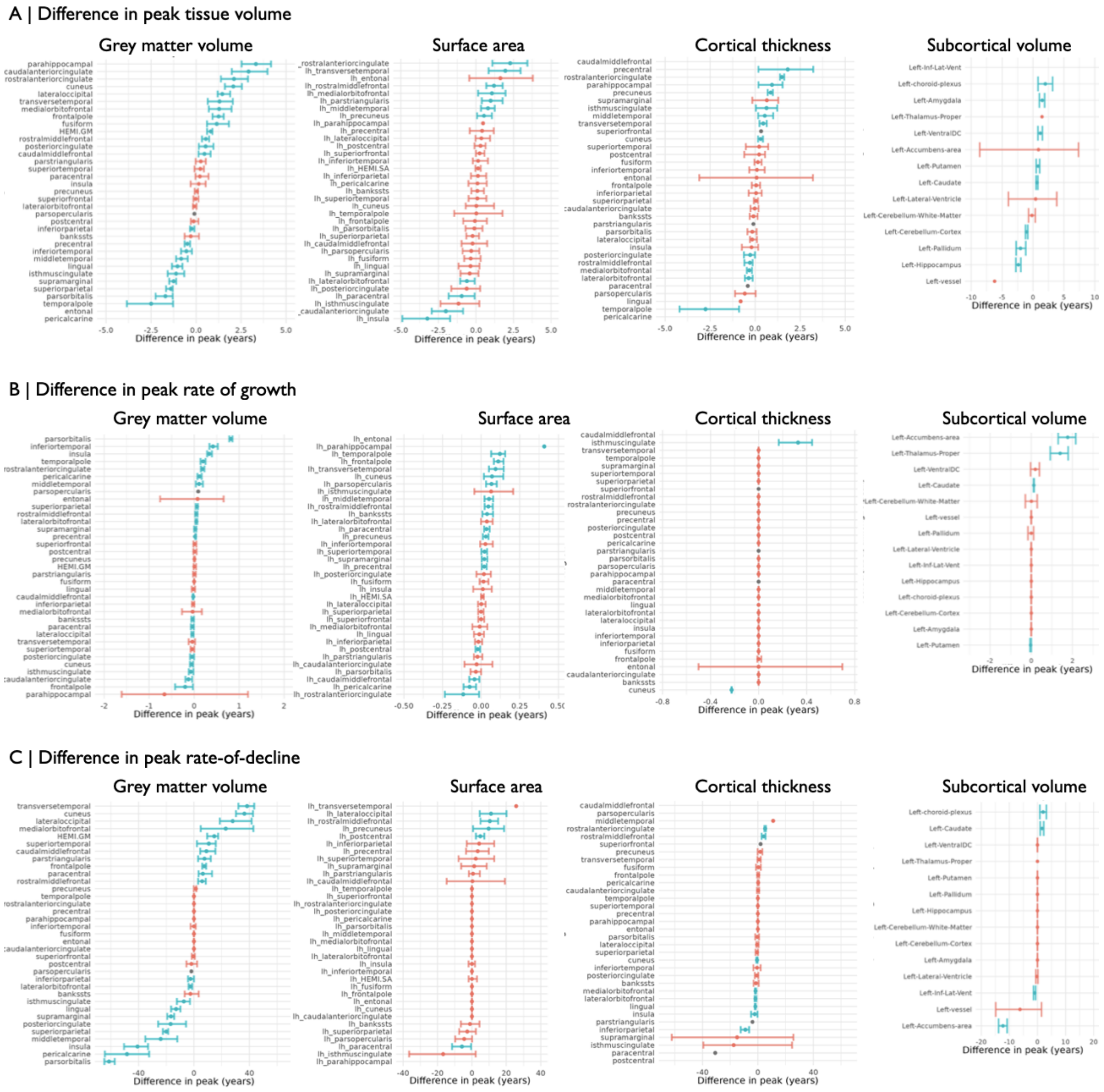

**Fig. 16: Bootstrap confidence around differences in developmental milestones:** We show the hemispheric difference in the age at which each of the three milestones were reached in each region for each phenotype. The regions (on the y-axis) are sorted by their difference in peak age. We further added the bootstrap derived confidence intervals around that difference. In turquoise, we highlight regions that show significant bilateral differences in milestones.

The 'First In, Last Out' hypothesis in neurodevelopment suggests that brain regions which mature earliest in development - in particular primary sensory and motor cortices - are the last to deteriorate with age or pathology. Conversely, higher-order association areas such as the prefrontal cortex, which are among the last to mature, tend to be first affected in neurodegeneration, psychiatric conditions, and cognitive decline [50].

In line with this previous work, we characterized the interplay of developmental milestones, i.e. the relationship between the peak rate of growth and declines across all phenotypes.

Overall, we find that surface area tends to align closely to this principle. Our data possible does not allow a closer examination of the principle for cortical thickness, since allmost all regions show their peak rate of growth early in development, likely before the onset of our trajectories (and with that before the current state-of-the-art prenatal MRIs).

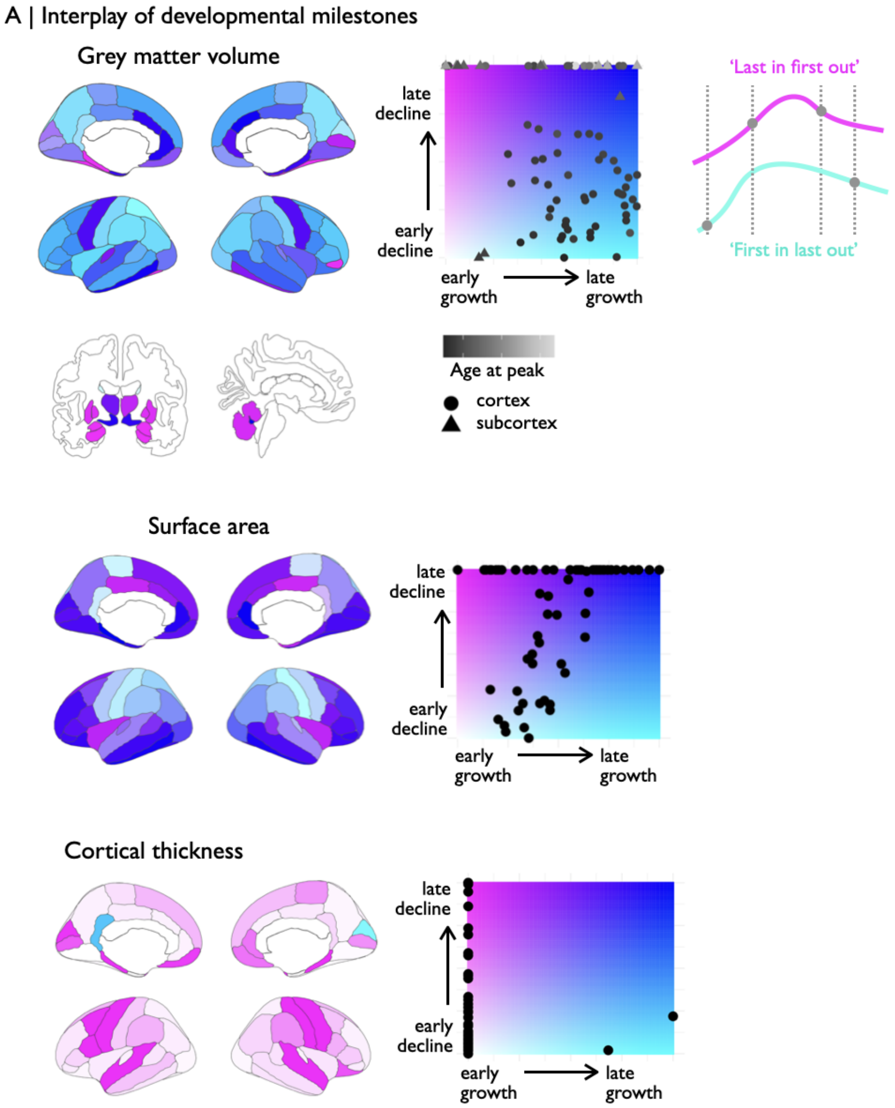

**Fig. 17: Interplay of developmental milestones:** For each phenotype, we illustrate the interplay between the rate of growth (x-axis) and the rate of decline (y-axis). We mapped the location on this field to colors on the cyan-pink spectrum. Pink regions tend to grow early, and decline late ('first in, last out'), bright turquoise/pink regions grow early and decline early ('first in, first out'), deeply turquoise regions grow late and decline early ('last in, first out'), blue regions grow late and decline late ('last in, last out').

#### Case-control differences in regional centile scores

Lastly, we estimated case-control differences in regional centile scores for three illustrative disorders, covering the lifespan from early life (Autism spectrum disorder, N=2421), through adolescence/early adulthood (Schizophrenia, N=1479), to late life (Alzheimer's disease, N=3070). We estimated the difference in centile scores for all pairwise comparisons of diagnostic groups and controls and characterized the effect size of this difference using Cohen's D, which describes the difference in group means in terms of the number of standard deviations that quantify the difference.

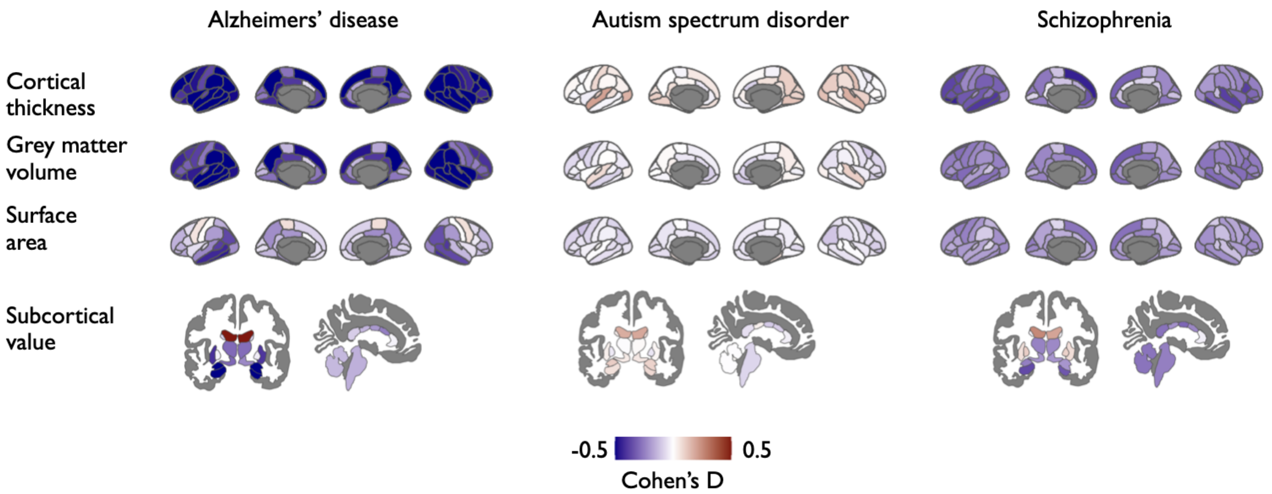

**Fig. 18: Regional case-control differences in centile scores:** Regional group mean case-control differences in centile scores for each phenotype (Cohen's D. Blue indicates regions that show decreases the case group in a given phenotype, compared to controls, whereas red indicates increases.

### Bootstrapped asymmetry trajectories

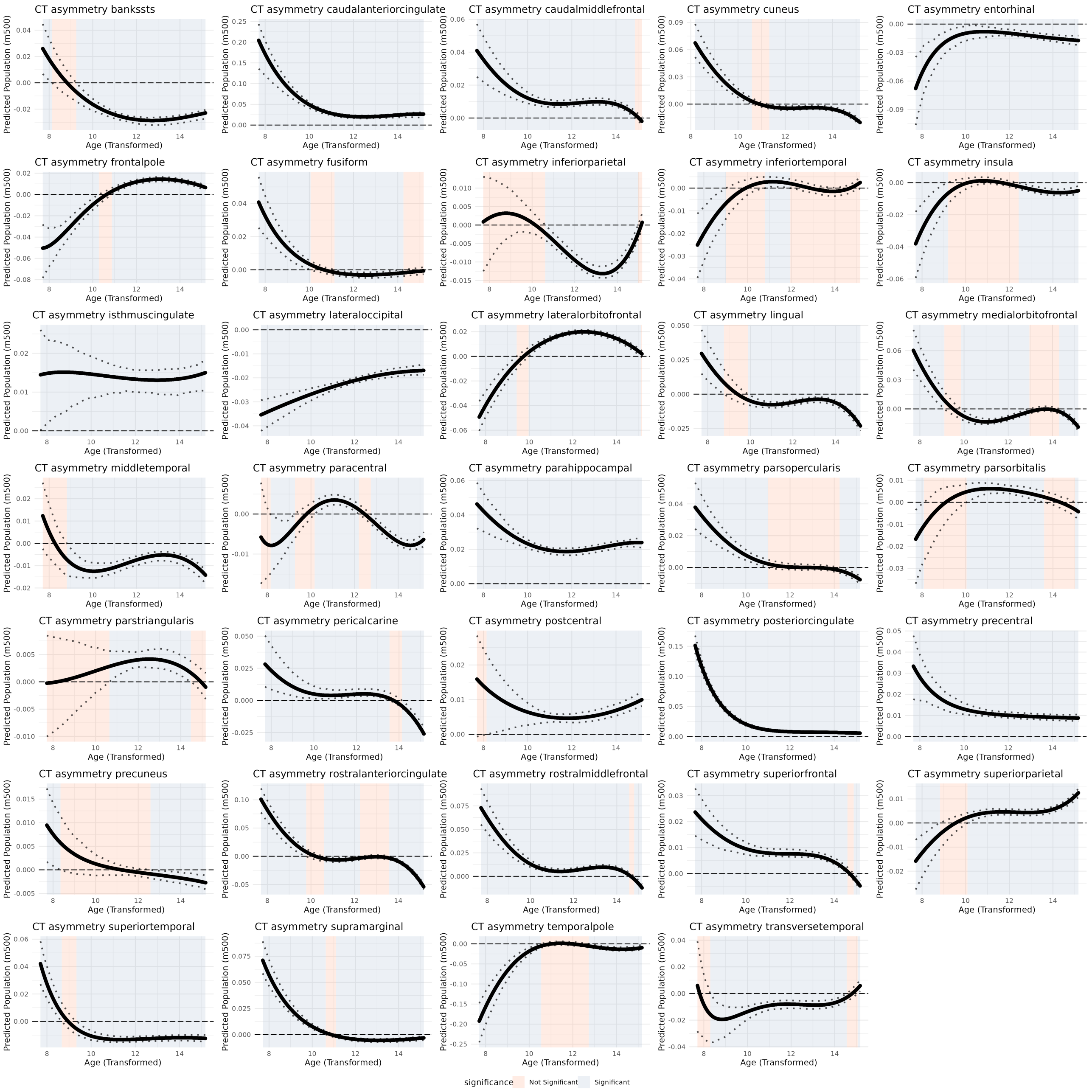

**Fig. 19: Bootstrap confidence of cortical thickness asymmetry trajectories: .**

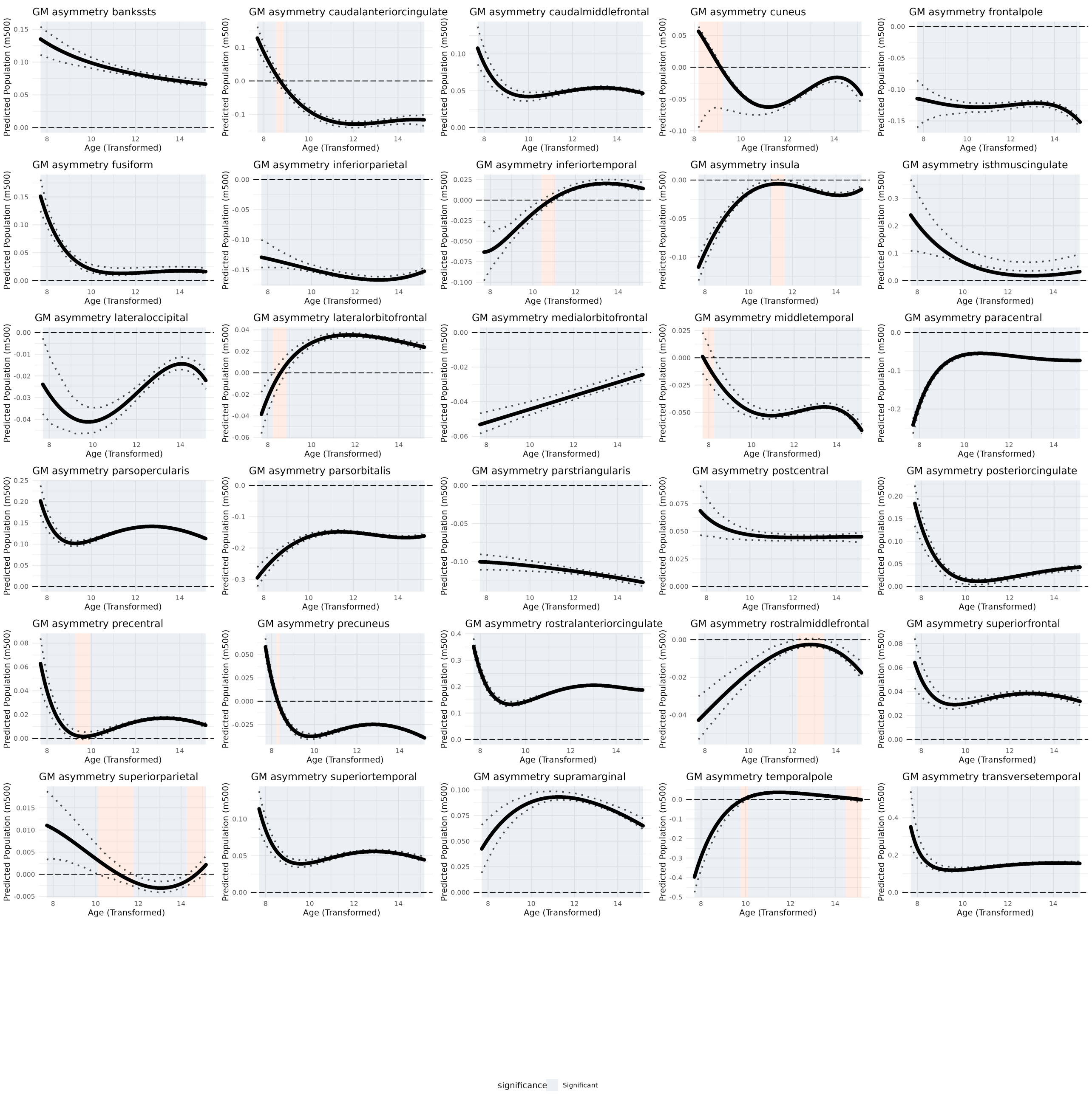

**Fig. 20: Bootstrap confidence of grey matter volume asymmetry trajectories: .**

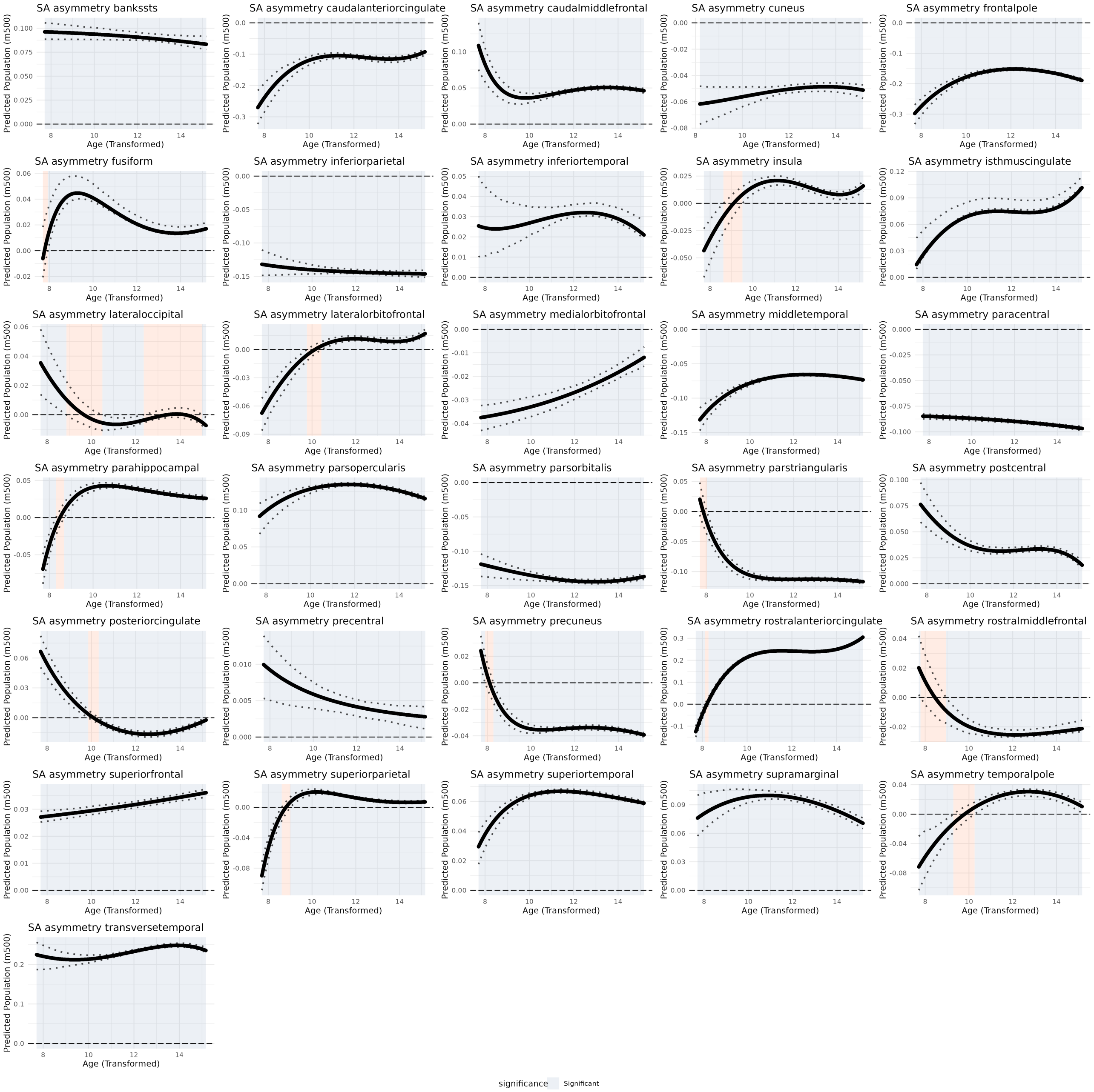

**Fig. 21: Bootstrap confidence of surface area asymmetry trajectories: .**

#### Quantification of magnitude of lifespan asymmetry

We characterized the magnitude of lifespan asymmetry for each phenotype by estimating the signed and absolute area under the curve of all regional asymmetry trajectories. A positive asymmetry value indicates left dominance, thus a positive area under the curves indicates that the region on average is left-dominant across the lifespan (**Main Fig. 3A**). We find that cortical thickness demonstrates a frontal-occipital gradient of left-to-right asymmetry. Grey matter volume and surface area show leftward asymmetry in tempero-parietal regions and a tendency for rightward asymmetry in frontal and occipital regions (**Fig. 22**). Further, overall grey matter volume and surface area asymmetry was greatest in multiple language regions (parsopercularis, parstriangularis, transversetemporal), as well as the anterior cingulate; whereas cortical thickness asymmetry was greatest in temporal regions, as well as the anterior cingulate (**Fig. 22**).

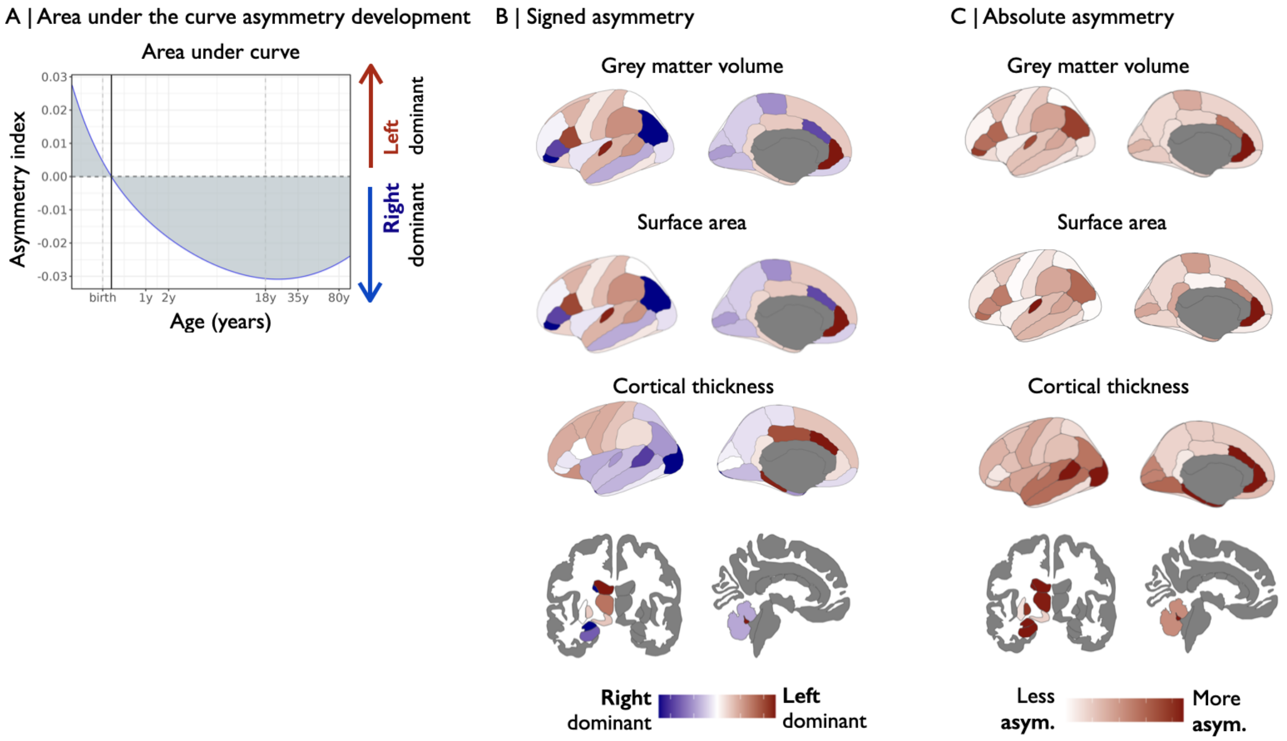

**Fig. 22: Area under the curve estimation of asymmetry:** (A) We aimed to estimate the magnitude of asymmetry over the lifespan by estimating the signed and unsigned area under the curve of the asymmetry trajectories. Negative asymmetry values indicate right dominance, thus a negative integral would indicate a right dominant region, whereas positive asymmetry values indicate left dominance, thus a positive integral would indicate a left dominant region. By estimating the summed integral, an average dominance value is estimated (B) Signed area under the curve. (C) Absolute area under the curve.

We next estimated the spatial correlation between epoch-specific asymmetry maps (**Main Fig. 3C**). This analysis revelead two clusters of asymmetry development: early and late development (**Fig. 23**), trends that were particularly pronounced in cortical thickness and subcortical volumes.

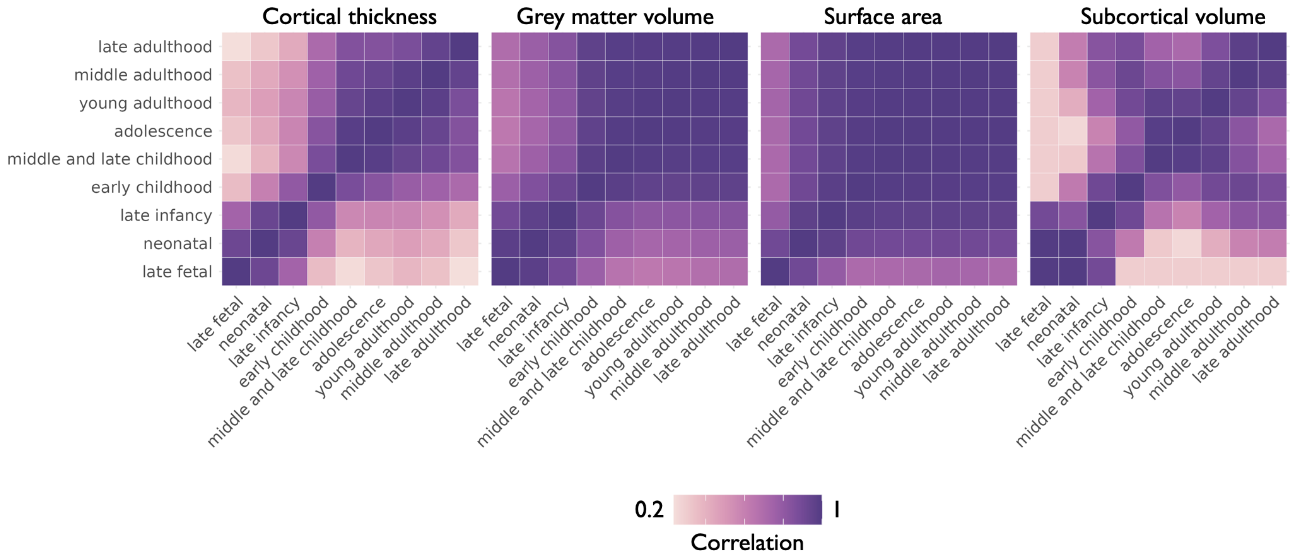

**Fig. 23: Developmental windows of asymmetry:** We estimated the spatial correlation between the age binned asymmetry maps by phenotype. This analysis really highlights two periods of asymmetry development, separated by early childhood: early development and ageing.

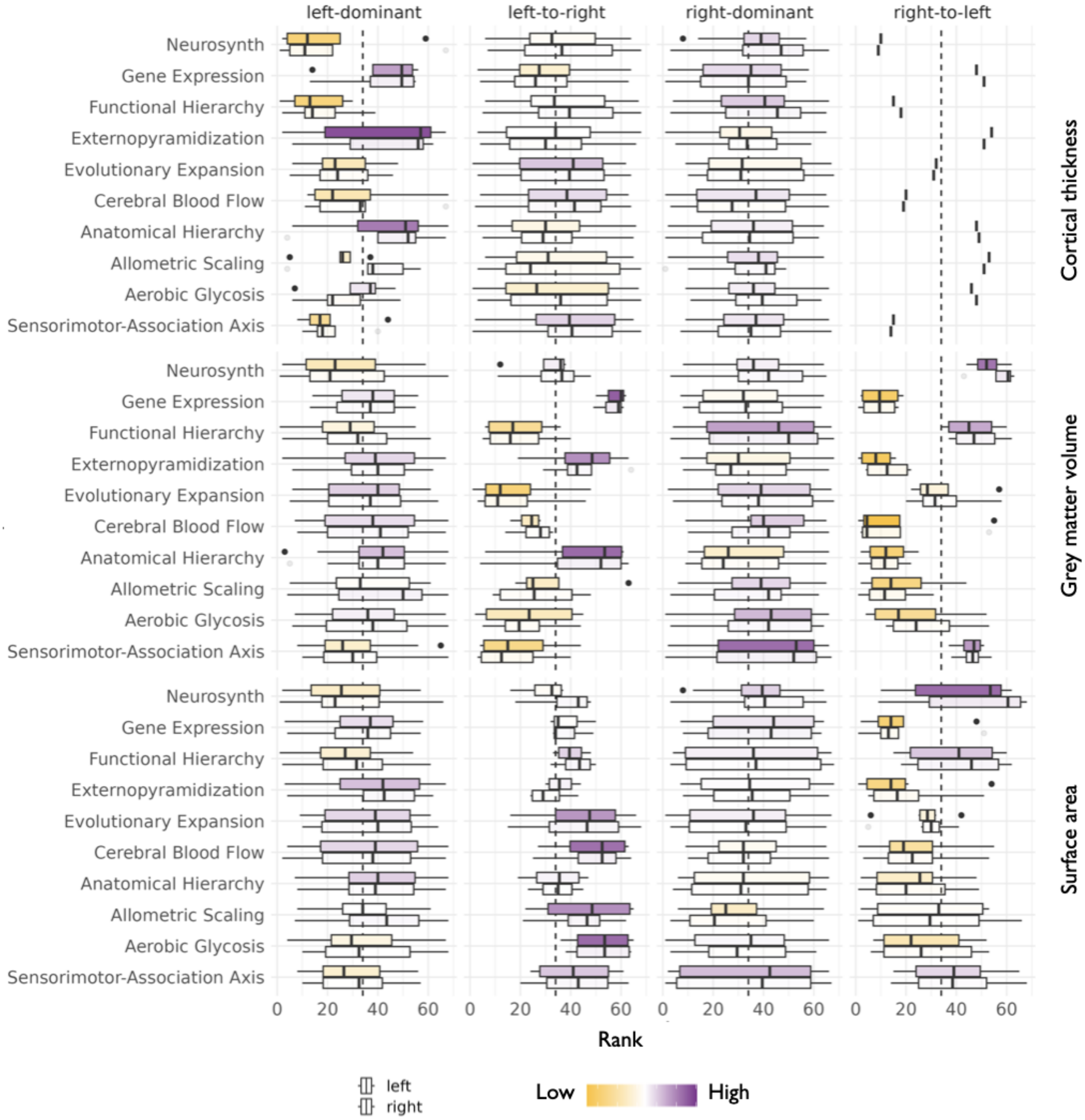

**Fig. 24: Asymmetry classes along the sensory-motor association axis:** We estimated the median value of the 10 composite maps of the sensory motor association axis map [129], as well as of the sensory-association axis map itself.

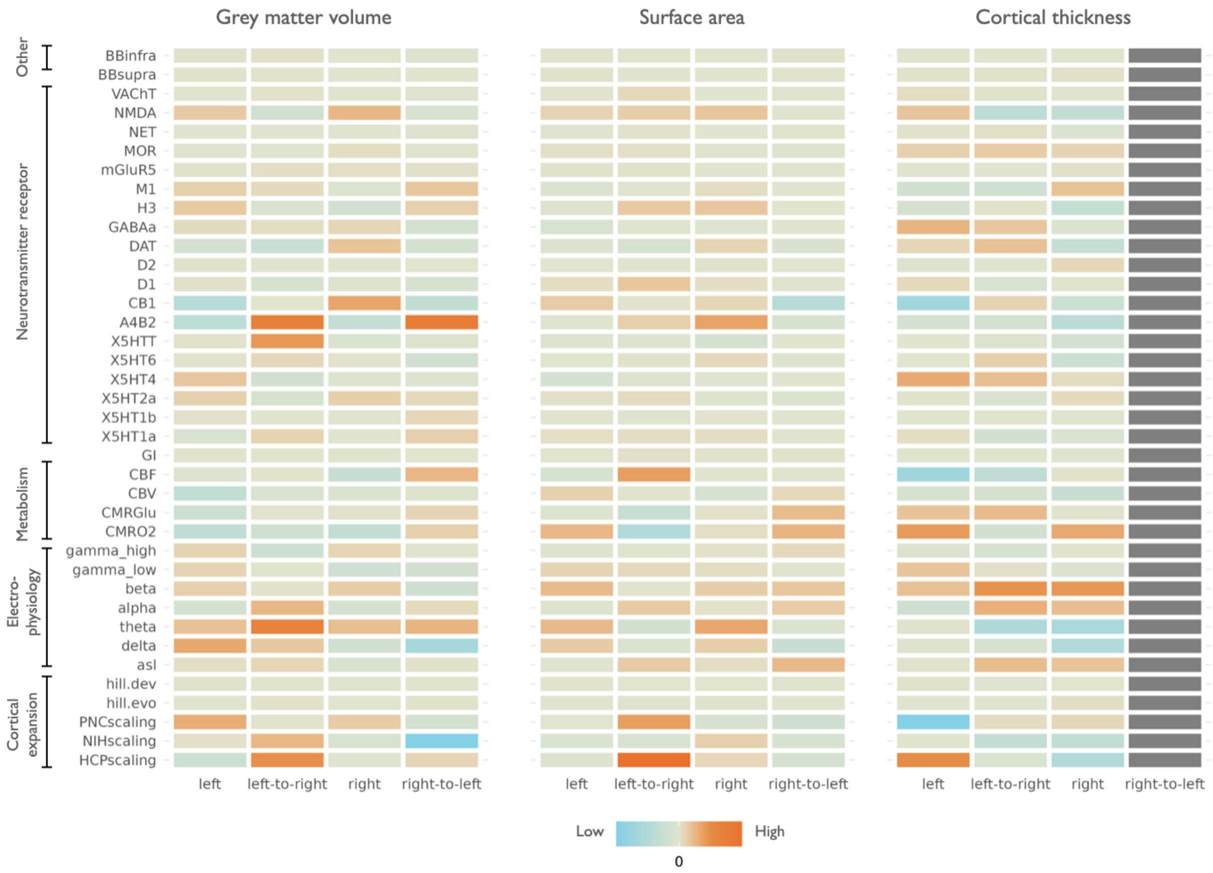

**Fig. 25: Structural and functional interpretation of asymmetry classes:** We contextualized each of four asymmetry classes using a set of reference maps of the human brain, including molecular, microstructural, electrophysiological, developmental and functional ontologies [81]. For each map, we averaged the respective values of regions assigned to each asymmetry class.

We assigned each region to one of four asymmetry classes, based on their asymmetry value at birth and and at the end of life: left-dominant regions (yellow), right-dominant regions (purple), left-to-right (pink) and right-to-left (blue) transitioning regions (**Fig. 26**). This analysis relvealed spatially consistent clusters of asymmetry classes.

The dual-origin theory posits that the mammalian cortex evolved from two distinct primordial cortical regions [2, 98]: (1) Hippocampal/ventral paleocortex origin (associated with medial and ventral structures like the hippocampus, entorhinal cortex, and parahippocampal gyrus); (2) Neocortical/dorsal archicortex origin (associated with dorsolateral structures like primary sensory cortices, prefrontal, and parietal cortex). We highlighted the 10 regions with the highest magnitude of changes in asymmetry over the lifespan (operationalized as the absolute difference in minimum vs maximum asymmetry over the lifespan). We find that these 10 regions clustered in regions associated with the dual origin theory of cortical evolution (**Fig. 26**). Specifically, highlighted regions included the periallocortical regions of the the cingulate, as well as multiple temporal regions.

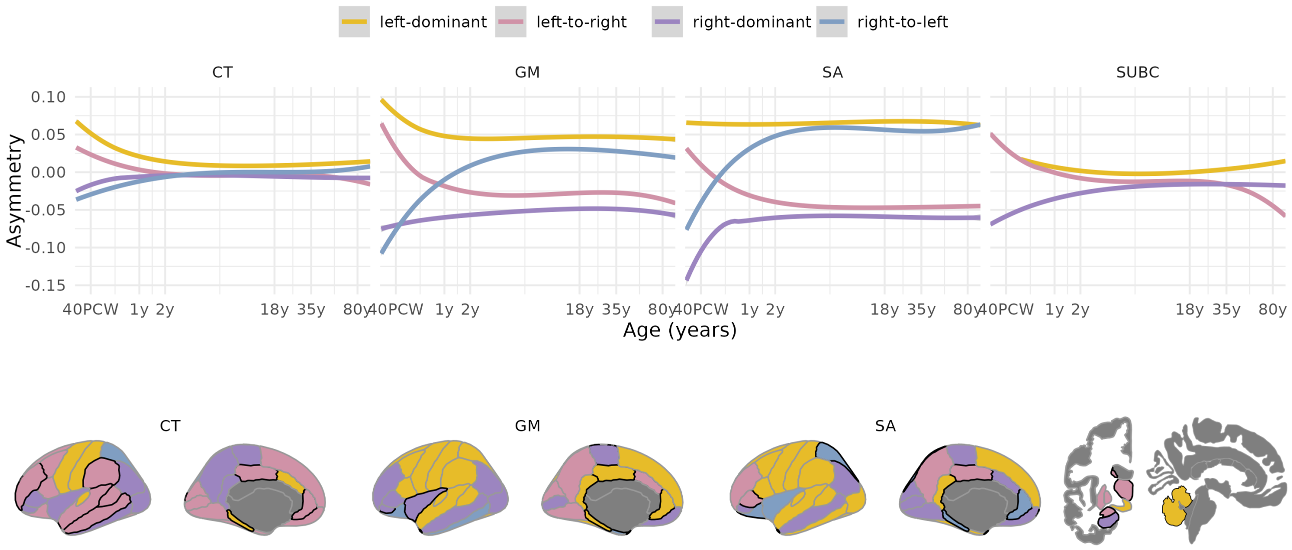

**Fig. 26: Largest changes in asymmetry across lifespan:** (A) We grouped asymmetry trajectories into four groups: left-dominant, left-to-right, right-dominant, right-to-left. We then averaged within these groups, across regions, and show the mean group trajectory. (B) For each phenotype, we found the 10 regions that showed the greated changes in asymmetry across the lifespan (highlighted by black outlines).

#### Case-control differences in asymmetry centile scores

We estimated case-control differences in asymmetry centile scores for four exemplary disorders. We characterized the effect size of the group mean difference using Cohen's D (**Fig. 27**). The interpration of this group mean difference is slightly more complicated than for the left/right regional models above. A negative Cohen's D value indicated that cases had decreased centile scores compared to controls. However, since asymmetry trajectories can be either left or right dominant (or close to symmetric), this can mean different things for different trajectories. In general, a decreased asymmetry centile score indicates a shift towards right-dominance. However, depending on the value of median trajectory in controls, this shift can mean one of many things. Hypothetically: (i) In controls, a region is strongly left-dominant, so decreased centile scores could mean that in cases, the region is symmetric, or right dominant. (ii) In controls, a region is right-dominant, so decreased centile scores would mean that a region is even more right-dominant in controls.

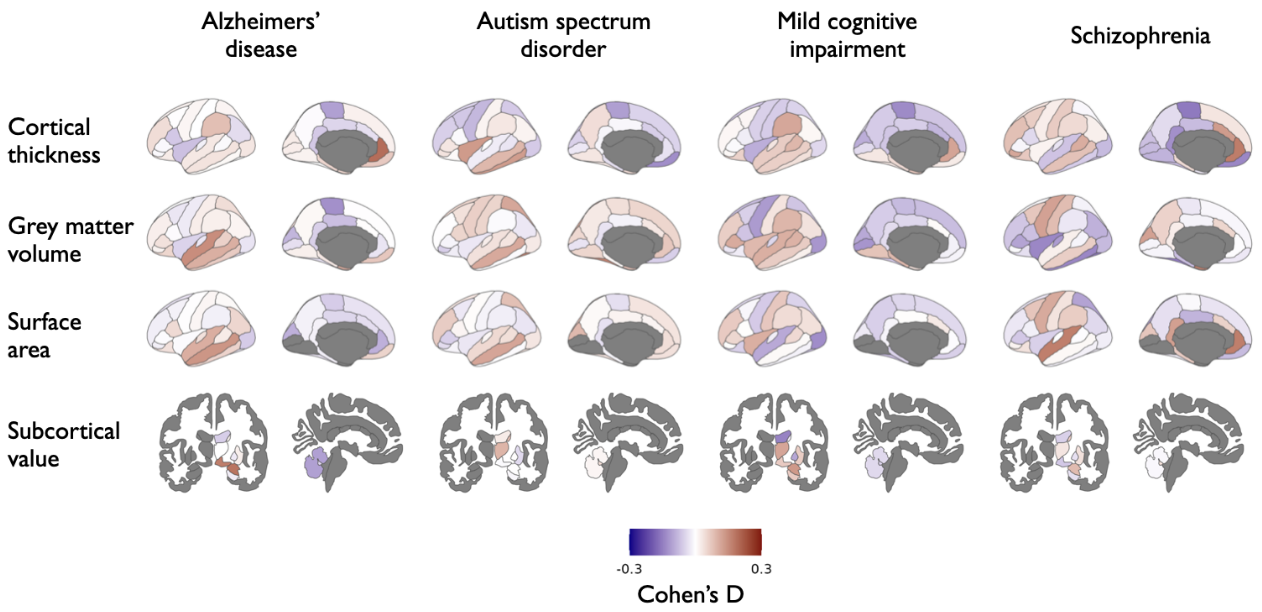

**Fig. 27: Case-control differences in asymmetry centile scores:** Effect size of group mean differences between cases and controls. Blue indicates decreased centile scores in cases compared to controls, red indicates increased centile scores in cases.

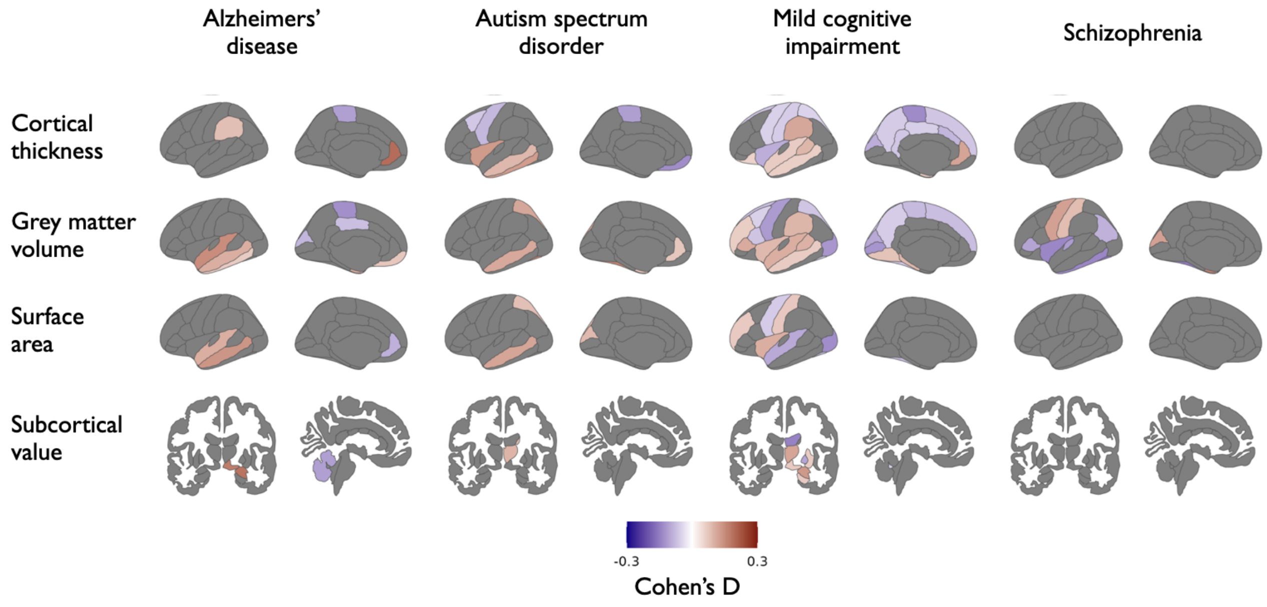

**Fig. 28: Significant case-control differences in asymmetry centile scores:** Thresholded map of case-control differences showing only regions that show significant differences after FDR-correction.

It is conceivable that neuropsychiatric disorders could lead to changes in the distribution of centile scores, compared to controls. For example, there may be changes in the number of subjects with extreme centile scores in neuropsychiatric disorders. For controls, the assumption is that 10% of subjects have centile scores below 0.1 and above 0.9, respectively. Here, we assessed whether this is true for subjects with neuropsychiatric disorders, too. We therefore estimated the percentage of subjects with centile scores below 0.1 and above 0.9.

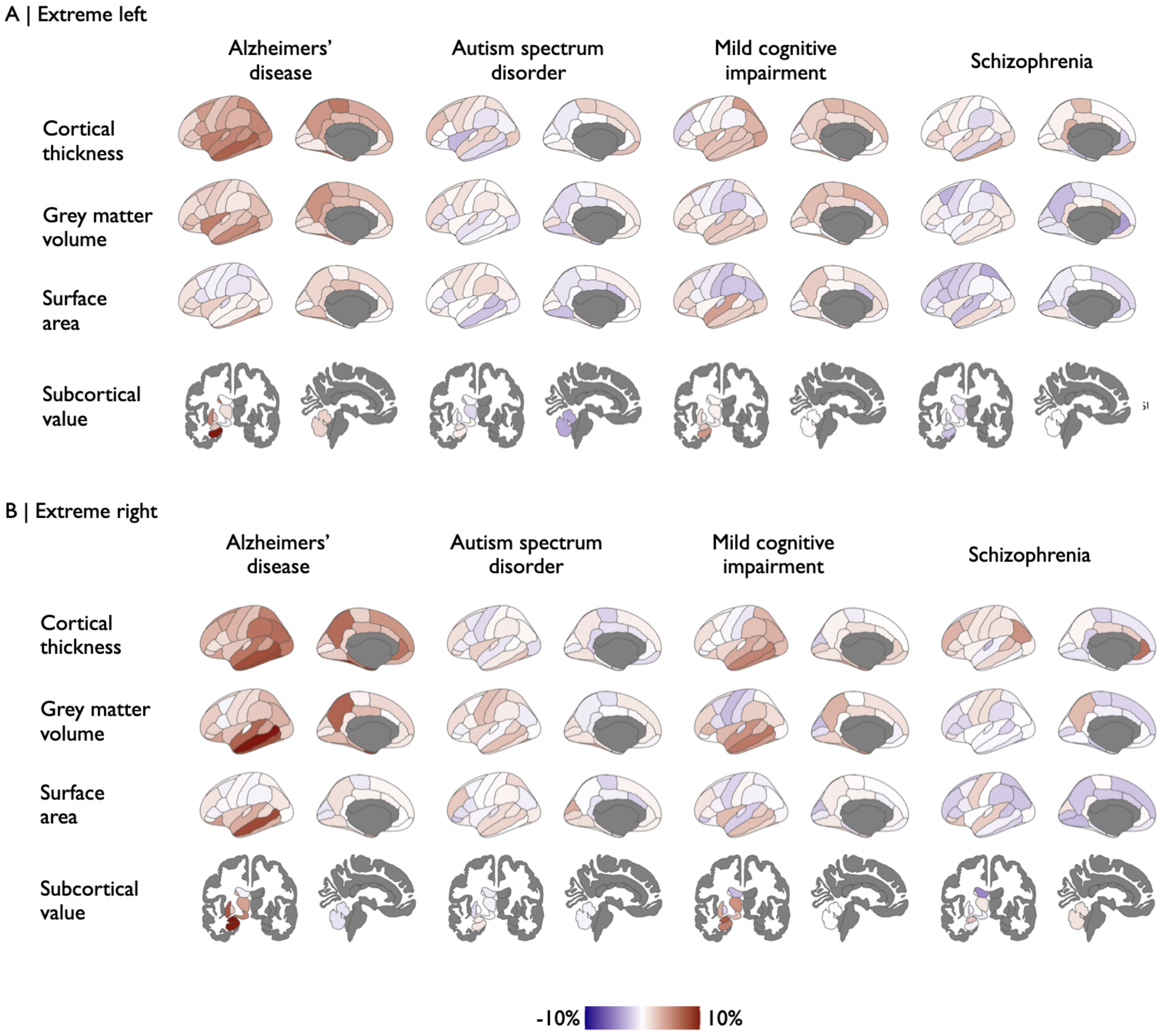

**Fig. 29: Extreme centile scores in cases:** We estimated the percentage of subjects with extreme centile scores, i.e. below 0.1 or above 0.9. For easy of interpretation, we subtracted 10 (the expected percentage of subjects in controls), such that the resulting values indicate deviations from normal.

Lastly, having observed changes in the number of subjects with extreme centile scores across disorders, we estimated the standard deviation in centile scores by diagnostic group.

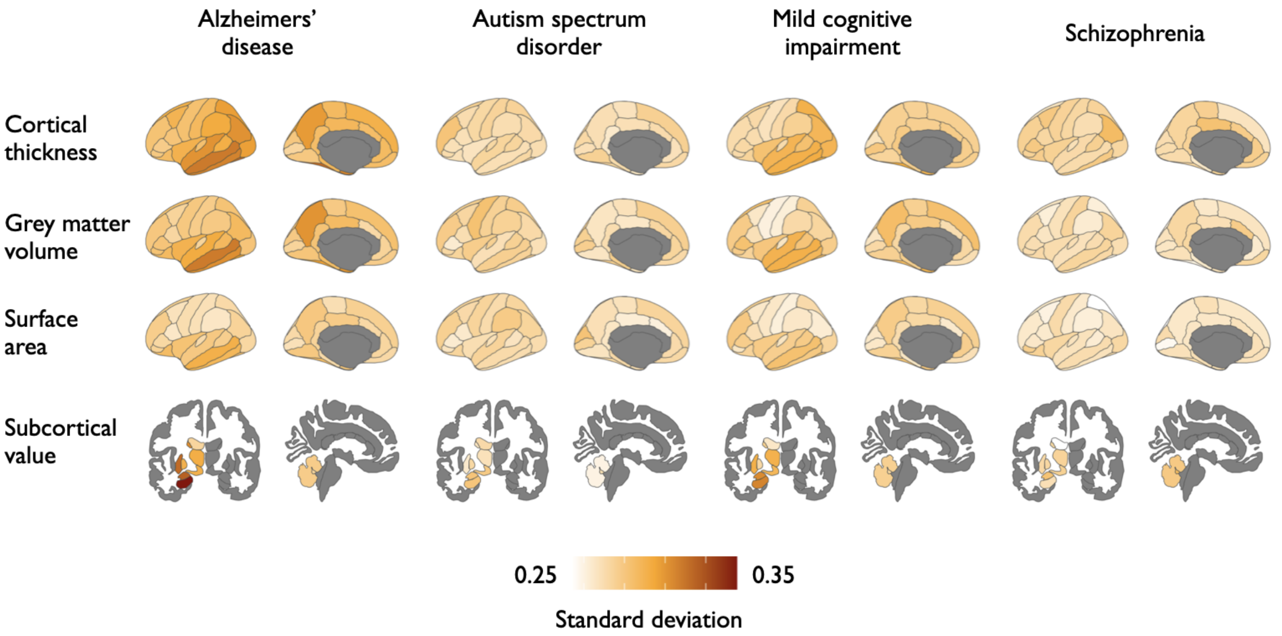

**Fig. 30: Distribution changes in asymmetry centile scores in cases:** Standard deviation in raw centile scores by diagnostic group.

#### Effect of handedness on asymmetry of centile scores

We assessed the relationship between handedness (right vs other) and asymmetry centile scores using the largest sample we had available to us. This included N subjects from 21 studies (**SI Table 5**). We used the handedness assignment as reported by the individual studies. For simplicity, we only included baseline data (one scan per subject) for this analysis. Due to the imbalance in the number of right handers compared to others, we combined left handers and ambidextruous subjects into a single category, we called 'non-righthandedness'.

| study | right | left | ambidextrous |
|---|---|---|---|
| study-ABCD | 9578 | 892 | 1720 |
| study-BGSP | 1437 | 109 | 11 |
| study-BIOCARD | 182 | 23 | 0 |
| study-CANBIND | 245 | 23 | 7 |
| study-CubanHCP | 152 | 19 | 5 |
| study-HCP | 947 | 67 | 99 |
| study-MCI | 172 | 7 | 12 |
| study-NACC | 4586 | 434 | 144 |
| study-NKI | 584 | 56 | 25 |
| study-OpenNeuroID1000 | 824 | 102 | 0 |
| study-OpenNeuroNIMHHV | 124 | 7 | 12 |
| study-OpenNeuroPIOP1 | 62 | 8 | 2 |
| study-OpenNeuroPixar | 139 | 10 | 3 |
| study-OpenNeuroSUDMEX | 112 | 9 | 7 |
| study-OpenNeuroWang | 206 | 4 | 0 |
| study-PACCT | 203 | 39 | 0 |
| study-PING | 603 | 80 | 30 |
| study-PNC | 1262 | 211 | 36 |
| study-QTIM | 1042 | 0 | 0 |
| study-SRPBS1600 | 1325 | 67 | 0 |
| study-UKB | 7752 | 804 | 130 |

**Table 5: Subjects with handedness data by study:** We list the number of unique subjects with handedness information (left/right/ambidextrous) as reported by the individual studies.

We estimated the effect of non-righthandedness on regional asymmetry centile scores in each phenotype using linear mixed effects models with a width fixed effects of age and sex, and a random effect of study. Due to the small number of subjects that were non-righthanders at some sites, we opted to control for study, rather than site in this analysis.

A negative effect of handedness indicates a lower centile scores in non-righthanders compared to right-handers. In a left-dominant region, this means that non-righthanders are less left-dominant, or more symmetric. In a right-dominant regions, it means that non-righthanders are more right-dominant or less symmetric (**SI Fig 31A**).

We also estimated the relationship between the handedness effects and overall (absolute) asymmetry. These effects are relatively low, overall, but we do observe a low negative correlation in subcortex ($\rho = -0.7, P < 0.05$) and grey matter volume ($\rho = -0.3, P = 0.1$). This relationship possibly suggests that non-righthanders have tangentially more symmetric brains (**SI Fig 31B**).

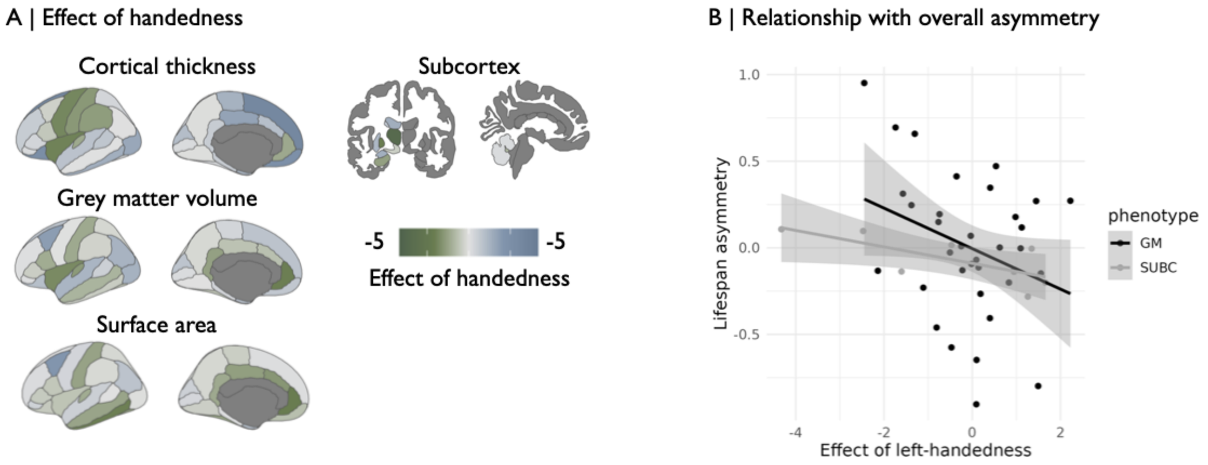

**Fig. 31: Effect of handedness on asymmetry centile scores:** (A) Effect of non-righthandedness on asymmetry centile scores, t-values. A negative t-value (green) indicates a lower centile scores in non-righthanders compared to right-handers. (B)

Lastly, we also estimated the effects of handedness in the two largest individual datasets available (**SI Fig 32**): ABCD (childhood) and UK Biobank (late adulthood). Here, we used linear models, with fixed effects of age and sex.

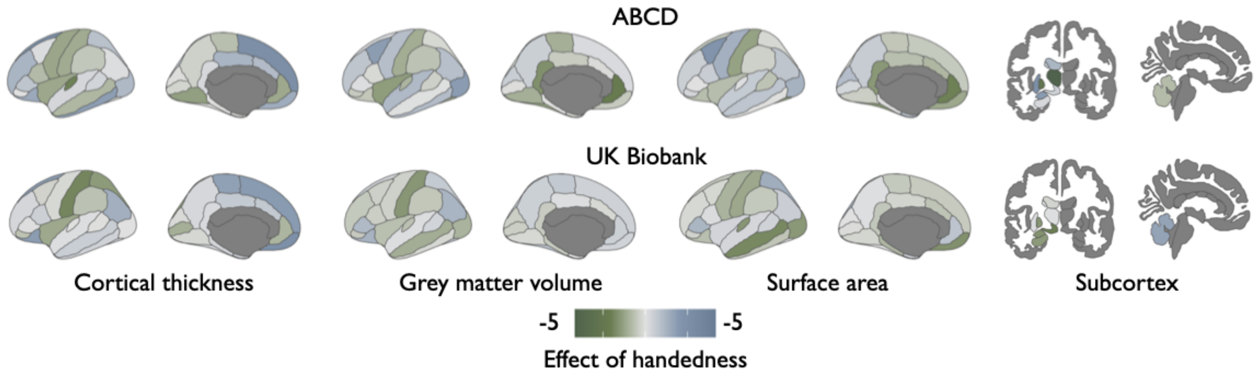

**Fig. 32: Effect of handedness in UKB and ABCD:** (A) .

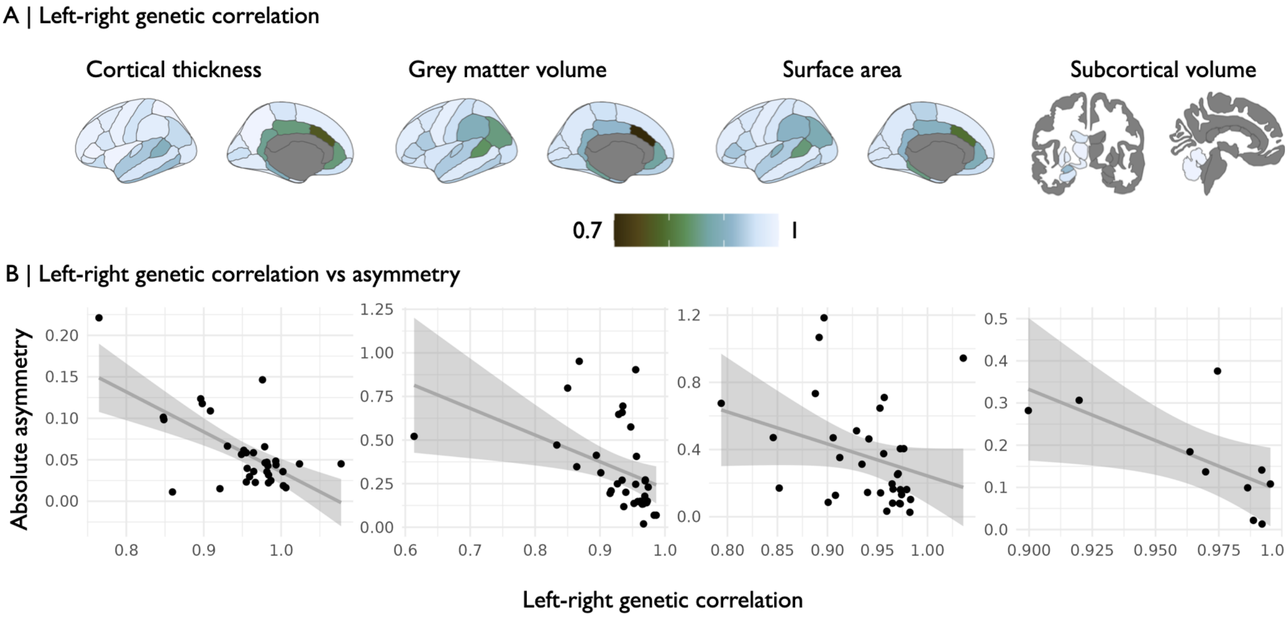

**Fig. 33: Relationship between genetic hemispheric similarity and asymmetry:** (A) Left-right genetic correlation for grey matter volume, cortical thickness and surface area. (B) Correlation between the absolute asymmetry map and the map of left-right genetic correlation values for each phenotype. Across phenotypes, we observed a negative correlation, indicating that regions that have lower left-right genetic correlation, tend to be more asymmetric ($\rho_{GM} = -0.63, P_{GM} < 0.01, P_{spin} < 0.01$; $\rho_{CT} = -0.46, P_{CT} < 0.01, P_{spin} < 0.01$; $\rho_{SA} = -0.43, P_{SA} < 0.05, P_{spin} < 0.01$; $\rho_{SUBC} = -0.69, P_{SUBC} < 0.05$).

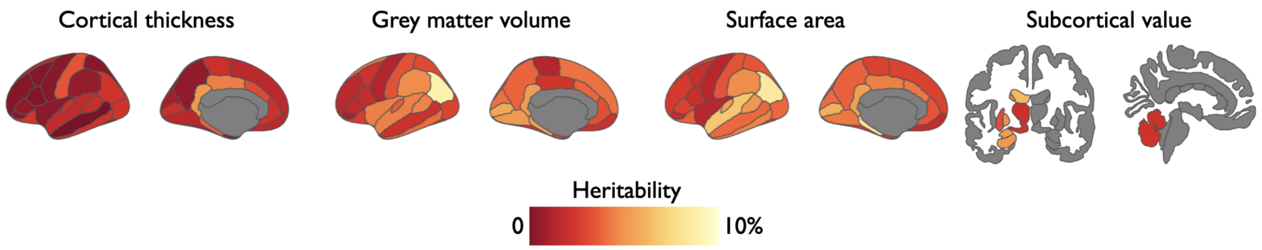

**Fig. 34: SNP-based heritability of asymmetry centile scores:** We estimated the SNP-based heritability of asymmetry centile scores.

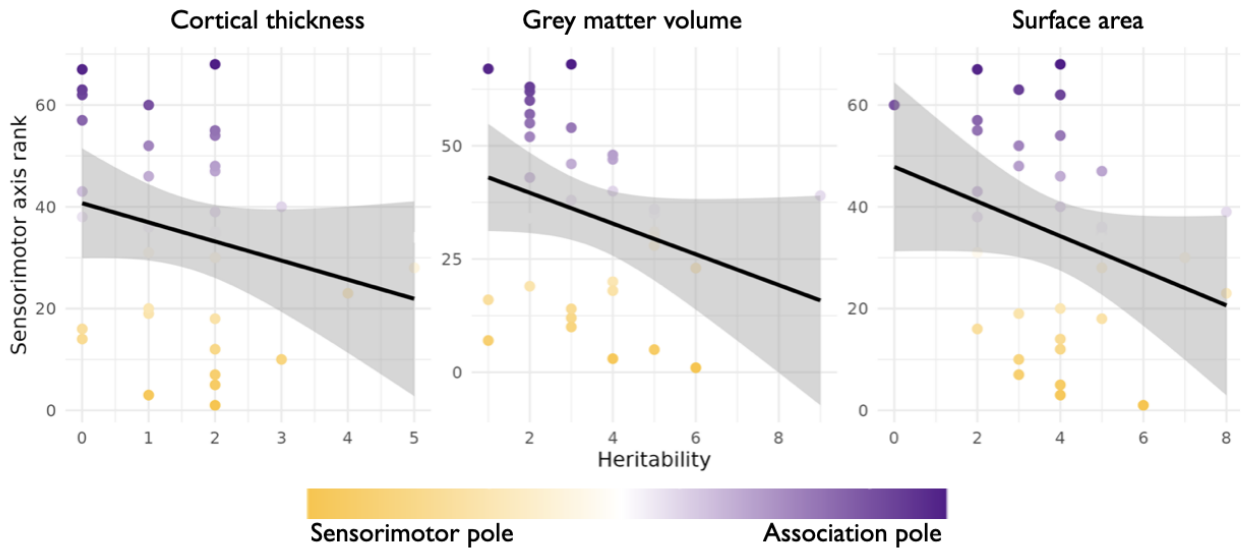

**Fig. 35: Asymmetry heritability develops along sensorimotor axis:** We estimated the correlation between the sensorimotor axis [129] and the maps of asymmetry centile score heritability for grey matter volume ($\rho_{GM} = -0.4, P < 0.05, P_{spin} < 0.05$), surface area ($\rho_{SA} = -0.35; P < 0.05, P_{spin} < 0.05$), and ($\rho_{CT} = -0.28; P = 0.1, P_{spin} = 0.07$).

### Translational potential of centile scores

Recent advances have been made to widen the geographical and demographic inclusivity of MRI data. Such efforts include the establishment of quick-scan MRI protocols [140], and low-field scanning consortia [34, 1], as well as the use of mobile MRI scanners [34] which allow access to previously less-studied communities.

To assess the translational potential of our braincharts, we assessed the reliability of centile scores estimated in ultra low resolution MRI data. We gained access to a dataset that included low (64mT; Hyperfine Swoop, Guildford,CT; hardware version: 1.7; software version: 8.6.0) and high (3T; GE Healthcare SIGNA Premier, Waukesha, WI) resolution anatomical scans acquired in the same set of 23 adult (20–64 years) subjects [141].

Acquisition and processing can be found in [141], but briefly: The low resolution scans were acquired using three product sequences, with higher resolution within either the axial, coronal or sagittal plane ($1.6 \times 1.6 \times 5mm^3$; AXI, COR, SAG respectively). These scans were subsequently combined into a single scan within each participant, using multi-resolution registration [34]. High resolution scans were acquired at $1mm^3$ isotropic resolution. All data were segmented using SynthSeg+ [17].

We generated out-of-sample centile scores for grey matter volume in the set of high and low resolution scans. We then estimated the Spearman correlation between high and low resolution centile scores within each region across subjects (**SI Fig. 36A**).

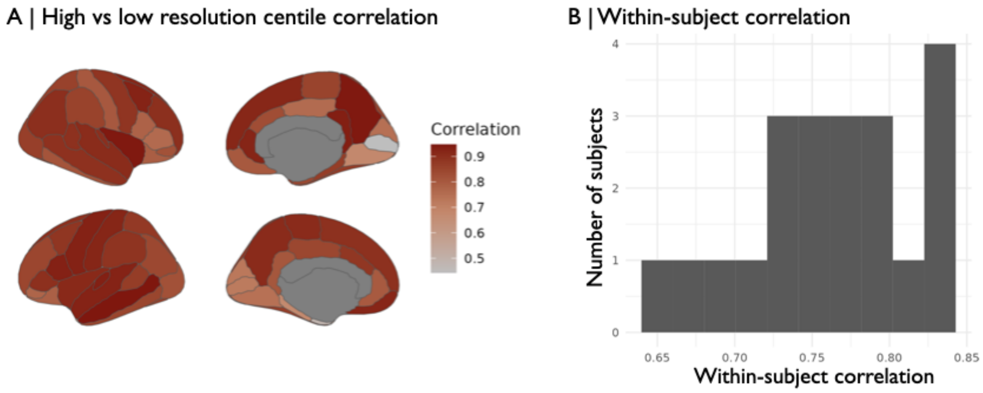

**Fig. 36: Ultra low resolution centile score consistency:** (A) Correlation of high vs low resolution centile scores within region across subjects. (B) Histogram of correlation values of within-subject high vs low resolution centile scores using LDSC.

### References

[1] F Abate et al. "UNITY: A low-field magnetic resonance neuroimaging initiative to characterize neurodevelopment in low and middle-income settings". In: *Developmental Cognitive Neuroscience* 69 (2024), p. 101397.

[2] Andrew Arthur Abbie. "Cortical lamination in the monotremata". In: *Journal of Comparative Neurology* 72.3 (1940), pp. 429–467.

[3] Jean Addington et al. "North American prodrome longitudinal study (NAPLS 3): methods and baseline description". In: *Schizophrenia research* 243 (2022), pp. 262–267.

[4] CJ Aine et al. "Multimodal neuroimaging in schizophrenia: description and dissemination". In: *Neuroinformatics* 15 (2017), pp. 343–364.

[5] Marilyn Albert et al. "Cognitive changes preceding clinical symptom onset of mild cognitive impairment and relationship to ApoE genotype". In: *Current Alzheimer Research* 11.8 (2014), pp. 773–784.

[6] Bonnie Alexander et al. "A new neonatal cortical and subcortical brain atlas: the Melbourne Children's Regional Infant Brain (M-CRIB) atlas". In: *Neuroimage* 147 (2017), pp. 841–851.

[7] Lindsay M Alexander et al. "An open resource for transdiagnostic research in pediatric mental health and learning disorders". In: *Scientific data* 4.1 (2017), pp. 1–26.

[8] C Robert Almli, Michael J Rivkin, Robert C McKinstry, Brain Development Cooperative Group, et al. "The NIH MRI study of normal brain development (Objective-2): newborns, infants, toddlers, and preschoolers". In: *Neuroimage* 35.1 (2007), pp. 308–325.

[9] Nicola Amoroso, Marianna La Rocca, Alfonso Monaco, Roberto Bellotti, and Sabina Tangaro. "Complex networks reveal early MRI markers of Parkinson's disease". In: *Medical image analysis* 48 (2018), pp. 12–24.

[10] Nancy C Andreasen, Peg Nopoulos, Vincent Magnotta, Ronald Pierson, Steven Ziebell, and Beng-Choon Ho. "Progressive brain change in schizophrenia: a prospective longitudinal study of first-episode schizophrenia". In: *Biological psychiatry* 70.7 (2011), pp. 672–679.

[11] Anahit Babayan et al. "A mind-brain-body dataset of MRI, EEG, cognition, emotion, and peripheral physiology in young and old adults". In: *Scientific data* 6.1 (2019), pp. 1–21.

[12] Pierre Bellec, Carlton Chu, Francois Chouinard-Decorte, Yassine Benhajali, Daniel S Margulies, and R Cameron Craddock. "The neuro bureau ADHD-200 preprocessed repository". In: *Neuroimage* 144 (2017), pp. 275–286.

[13] David A Bennett, Aron S Buchman, Patricia A Boyle, Lisa L Barnes, Robert S Wilson, and Julie A Schneider. "Religious orders study and rush memory and aging project". In: *Journal of Alzheimer's disease* 64.s1 (2018), S161–S189.

[14] Lilah Besser et al. "Version 3 of the national Alzheimer's coordinating center's uniform data set". In: *Alzheimer Disease & Associated Disorders* 32.4 (2018), pp. 351–358.

[15] Richard AI Bethlehem et al. "Brain charts for the human lifespan". In: *Nature* 604.7906 (2022), pp. 525–533.

[16] R Bilder, R Poldrack, T Cannon, E London, N Freimer, E Congdon, K Karlsgodt, and F Sabb. *"UCLA Consortium for Neuropsychiatric Phenomics LA5c Study"*. OpenNeuro, 2020. DOI: `10.18112/openneuro.ds000030.v1.0.0`.

[17] Benjamin Billot, Douglas N Greve, Oula Puonti, Axel Thielscher, Koen Van Leemput, Bruce Fischl, Adrian V Dalca, Juan Eugenio Iglesias, et al. "SynthSeg: Segmentation of brain MRI scans of any contrast and resolution without retraining". In: *Medical image analysis* 86 (2023), p. 102789.

[18] Elaine Borghi et al. "Construction of the World Health Organization child growth standards: selection of methods for attained growth curves". In: *Statistics in medicine* 25.2 (2006), pp. 247–265.

[19] Rotem Botvinik-Nezer, Bogdan Petre, Marta Ceko, Naomi Friedman, and Tor Wager. *"Paingen˙placebo"*. OpenNeuro, 2023. DOI: `doi:10.18112/openneuro.ds004746.v1.0.1`.

[20] Rotem Botvinik-Nezer, Bogdan Petre, Marta Ceko, Martin A Lindquist, Naomi P Friedman, and Tor D Wager. "Placebo treatment affects brain systems related to affective and cognitive processes, but not nociceptive pain". In: *Nature communications* 15.1 (2024), p. 6017.

[21] Niall J Bourke et al. "Brain volume abnormalities and clinical outcomes following paediatric traumatic brain injury". In: *Brain* 145.8 (2022), pp. 2920–2934.

[22] Katharina Brueggen et al. "The European DTI Study on Dementia—A multicenter DTI and MRI study on Alzheimer's disease and mild cognitive impairment". In: *Neuroimage* 144 (2017), pp. 305–308.

[23] Brendan Bulik-Sullivan et al. "An atlas of genetic correlations across human diseases and traits". In: *Nature genetics* 47.11 (2015), pp. 1236–1241.

[24] Brendan K Bulik-Sullivan et al. "LD Score regression distinguishes confounding from polygenicity in genome-wide association studies". In: *Nature genetics* 47.3 (2015), pp. 291–295.

[25] Clare Bycroft et al. "The UK Biobank resource with deep phenotyping and genomic data". In: *Nature* 562.7726 (2018), pp. 203–209.

[26] Chris C Camp, Stephanie Noble, Dustin Scheinost, Argyris Stringaris, and Dylan M Nielson. "Test-retest reliability of functional connectivity in depressed adolescents". In: *medRxiv* (2022), pp. 2022–10.

[27] Betty Jo Casey et al. "The adolescent brain cognitive development (ABCD) study: imaging acquisition across 21 sites". In: *Developmental cognitive neuroscience* 32 (2018), pp. 43–54.

[28] Enrica Cavedo et al. "The Italian Alzheimer's Disease Neuroimaging Initiative (I-ADNI): Validation of structural MR imaging". In: *Journal of Alzheimer's Disease* 40.4 (2014), pp. 941–952.

[29] Anni Copeland et al. "Infant and child MRI: a review of scanning procedures". In: *Frontiers in neuroscience* 15 (2021), p. 666020.

[30] Nicolas A Crossley et al. "Structural brain abnormalities in schizophrenia in adverse environments: examining the effect of poverty and violence in six Latin American cities". In: *The British Journal of Psychiatry* 218.2 (2021), pp. 112–118.

[31] Alexander Dagley et al. "Harvard aging brain study: dataset and accessibility". In: *Neuroimage* 144 (2017), pp. 255–258.

[32] Ana M Daugherty and Naftali Raz. "A virtual water maze revisited: Two-year changes in navigation performance and their neural correlates in healthy adults". In: *NeuroImage* 146 (2017), pp. 492–506.

[33] Kara Dempster, Peter Jeon, Michael MacKinley, Peter Williamson, Jean Théberge, and Lena Palaniyappan. "Early treatment response in first episode psychosis: a 7-T magnetic resonance spectroscopic study of glutathione and glutamate". In: *Molecular psychiatry* 25.8 (2020), pp. 1640–1650.

[34] Sean CL Deoni et al. "Development of a mobile low-field MRI scanner". In: *Scientific reports* 12.1 (2022), p. 5690.

[35] Adriana Di Martino et al. "The autism brain imaging data exchange: towards a large-scale evaluation of the intrinsic brain architecture in autism". In: *Molecular psychiatry* 19.6 (2014), pp. 659–667.

[36] A David Edwards et al. "The developing human connectome project neonatal data release". In: *Frontiers in neuroscience* 16 (2022), p. 886772.

[37] Lloyd T Elliott, Kevin Sharp, Fidel Alfaro-Almagro, Sinan Shi, Karla L Miller, Gwenaëlle Douaud, Jonathan Marchini, and Stephen M Smith. "Genome-wide association studies of brain imaging phenotypes in UK Biobank". In: *Nature* 562.7726 (2018), pp. 210–216.

[38] Kathryn A Ellis et al. "The Australian Imaging, Biomarkers and Lifestyle (AIBL) study of aging: methodology and baseline characteristics of 1112 individuals recruited for a longitudinal study of Alzheimer's disease". In: *International psychogeriatrics* 21.4 (2009), pp. 672–687.

[39] Antao Chen Feng, Jiang Qiu, Xu Chen, Xun Liu, and Xi-Nian Zuo. "Chinese color nest project (CCNP) I: Growing up in China". In: *Chin. Sci. Bull* 62 (2017), pp. 3008–3022.

[40] Noa Fingher, Ilan Dinstein, Michal Ben-Shachar, Shlomi Haar, Anders M Dale, Lisa Eyler, Karen Pierce, and Eric Courchesne. "Toddlers later diagnosed with autism exhibit multiple structural abnormalities in temporal corpus callosum fibers". In: *Cortex* 97 (2017), pp. 291–305.

[41] Giovanni B Frisoni et al. "Markers of Alzheimer's disease in a population attending a memory clinic". In: *Alzheimer's & Dementia* 5.4 (2009), pp. 307–317.

[42] Bari Fuchs, Alaina Pearce, Barbara Rolls, Stephen Wilson, Emma Jane Rose, Charles Geier, Hugh Garavan, and Kathleen Keller. *"Food and Brain Study"*. OpenNeuro, 2023. DOI: doi:10.18112/openneuro.ds004697.v1.0.2.

[43] Peng Gao, Hao-Ming Dong, Si-Man Liu, Xue-Ru Fan, Chao Jiang, Yin-Shan Wang, Daniel Margulies, Hai-Fang Li, and Xi-Nian Zuo. "A Chinese multi-modal neuroimaging data release for increasing diversity of human brain mapping". In: *Scientific data* 9.1 (2022), p. 286.

[44] Peng Gao, Hao-Ming Dong, Yin-Shan Wang, Chun-Shui Yu, and Xi-Nian Zuo. *"Imaging Chinese Young Brains (I See Your Brain)"*. Version V1. Science Data Bank, 2021. DOI: doi:10.11922/sciencedb.00740.

[45] Antoine Garnier-Crussard et al. "White matter hyperintensities across the adult lifespan: relation to age, A$\beta$ load, and cognition". In: *Alzheimer's research & therapy* 12 (2020), pp. 1–11.

[46] Eduardo A Garza-Villarreal et al. "The effect of crack cocaine addiction and age on the microstructure and morphology of the human striatum and thalamus using shape analysis and fast diffusion kurtosis imaging". In: *Translational psychiatry* 7.5 (2017), e1122–e1122.

[47] Eduardo A. Garza-Villarreal, Jorge Julio Gonzalez Olvera, Thania Balducci, Diego Angeles Valdez, Alely Valencia, and Jalil Rasgado. *"SUDMEX˙CONN: The Mexican dataset of cocaine use disorder patients."*. OpenNeuro, 2021. DOI: doi:10.18112/openneuro.ds003346.v1.1.2.

[48] Jianqiao Ge et al. "Increasing diversity in connectomics with the Chinese Human Connectome Project". In: *Nature Neuroscience* 26.1 (2023), pp. 163–172.

[49] Matthew F Glasser et al. "The minimal preprocessing pipelines for the Human Connectome Project". In: *Neuroimage* 80 (2013), pp. 105–124.

[50] Nitin Gogtay et al. "Dynamic mapping of human cortical development during childhood through early adulthood". In: *Proceedings of the national academy of sciences* 101.21 (2004), pp. 8174–8179.

[51] H Hill Goldsmith, Kathryn Lemery-Chalfant, Nicole L Schmidt, Carrie L Arneson, and Cory K Schmidt. "Longitudinal analyses of affect, temperament, and childhood psychopathology". In: *Twin Research and Human Genetics* 10.1 (2007), pp. 118–126.

[52] Randy L Gollub and Dennis A Turner. *Neuroinformatics in Clinical and Translational Medicine—Novel Approaches*. 2010.

[53] Randy L Gollub et al. "The MCIC collection: a shared repository of multi-modal, multi-site brain image data from a clinical investigation of schizophrenia". In: *Neuroinformatics* 11 (2013), pp. 367–388.

[54] Neil Samuel Nyholm Graham et al. "Multicentre longitudinal study of fluid and neuroimaging BIOmarkers of AXonal injury after traumatic brain injury: the BIO-AX-TBI study protocol". In: *BMJ open* 10.11 (2020), e042093.

[55] Katrina L Grasby et al. "The genetic architecture of the human cerebral cortex". In: *Science* 367.6484 (2020), eaay6690.

[56] Michael P Harms et al. "Extending the Human Connectome Project across ages: Imaging protocols for the Lifespan Development and Aging projects". In: *Neuroimage* 183 (2018), pp. 972–984.

[57] Avram J Holmes et al. "Brain Genomics Superstruct Project initial data release with structural, functional, and behavioral measures". In: *Scientific data* 2.1 (2015), pp. 1–16.

[58] Joni Holmes, Annie Bryant, CALM Team calm@ mrc-cbu. cam. ac. uk, and Susan Elizabeth Gathercole. "Protocol for a transdiagnostic study of children with problems of attention, learning and memory (CALM)". In: *BMC pediatrics* 19 (2019), pp. 1–11.

[59] Brittany R Howell et al. "The UNC/UMN Baby Connectome Project (BCP): An overview of the study design and protocol development". In: *NeuroImage* 185 (2019), pp. 891–905.

[60] Clifford R Jack Jr et al. "The Alzheimer's disease neuroimaging initiative (ADNI): MRI methods". In: *Journal of Magnetic Resonance Imaging: An Official Journal of the International Society for Magnetic Resonance in Medicine* 27.4 (2008), pp. 685–691.

[61] Terry L Jernigan et al. "The pediatric imaging, neurocognition, and genetics (PING) data repository". In: *Neuroimage* 124 (2016), pp. 1149–1154.

[62] Longda Jiang, Zhili Zheng, Ting Qi, Kathryn E Kemper, Naomi R Wray, Peter M Visscher, and Jian Yang. "A resource-efficient tool for mixed model association analysis of large-scale data". In: *Nature genetics* 51.12 (2019), pp. 1749–1755.

[63] Tuomas Jukuri et al. "Default mode network in young people with familial risk for psychosis—The Oulu Brain and Mind Study". In: *Schizophrenia research* 143.2-3 (2013), pp. 239–245.

[64] Hyo Jung Kang et al. "Spatio-temporal transcriptome of the human brain". In: *Nature* 478.7370 (2011), pp. 483–489.

[65] Kristen M Kennedy, Karen M Rodrigue, Gérard N Bischof, Andrew C Hebrank, Patricia A Reuter-Lorenz, and Denise C Park. "Age trajectories of functional activation under conditions of low and high processing demands: an adult lifespan fMRI study of the aging brain". In: *Neuroimage* 104 (2015), pp. 21–34.

[66] Manfred G Kitzbichler et al. "Peripheral inflammation is associated with micro-structural and functional connectivity changes in depression-related brain networks". In: *Molecular psychiatry* 26.12 (2021), pp. 7346–7354.

[67] Ece Kocagoncu, David Nesbitt, Tina Emery, Laura E Hughes, Richard N Henson, James B Rowe, et al. "Neurophysiological and brain structural markers of cognitive frailty differ from Alzheimer's disease". In: *Journal of Neuroscience* 42.7 (2022), pp. 1362–1373.

[68] Jakub Kopal, Lucina Q Uddin, and Danilo Bzdok. "The end game: respecting major sources of population diversity". In: *Nature Methods* 20.8 (2023), pp. 1122–1128.

[69] William S Kremen et al. "Genetic and environmental influences on the size of specific brain regions in midlife: the VETSA MRI study". In: *Neuroimage* 49.2 (2010), pp. 1213–1223.

[70] Maria Kuklisova-Murgasova et al. "A dynamic 4D probabilistic atlas of the developing brain". In: *NeuroImage* 54.4 (2011), pp. 2750–2763.

[71] Raymond W Lam et al. "Discovering biomarkers for antidepressant response: protocol from the Canadian biomarker integration network in depression (CAN-BIND) and clinical characteristics of the first patient cohort". In: *BMC psychiatry* 16 (2016), pp. 1–13.

[72] Pamela J. LaMontagne et al. "OASIS-3: Longitudinal Neuroimaging, Clinical, and Cognitive Dataset for Normal Aging and Alzheimer Disease". In: *medRxiv* (2019). Publisher: Cold Spring Harbor Laboratory Press _eprint: https://www.medrxiv.org/content/early/2019/12/15/2019.12.13.19014902.full.pdf. DOI: 10.1101/2019.12.13.19014902. URL: https://www.medrxiv.org/content/early/2019/12/15/2019.12.13.19014902.

[73] Annie Lee et al. "Maternal care in infancy and the course of limbic development". In: *Developmental Cognitive Neuroscience* 40 (2019), p. 100714.

[74] John D Lewis et al. "The emergence of network inefficiencies in infants with autism spectrum disorder". In: *Biological psychiatry* 82.3 (2017), pp. 176–185.

[75] Siman Liu et al. "Chinese Color Nest Project : An accelerated longitudinal brain-mind cohort". In: *Developmental Cognitive Neuroscience* 52 (2021), p. 101020. ISSN: 1878-9293. DOI: https://doi.org/10.1016/j.dcn.2021.101020. URL: https://www.sciencedirect.com/science/article/pii/S1878929321001109.

[76] Wei Liu et al. "Longitudinal test-retest neuroimaging data from healthy young adults in southwest China". In: *Scientific data* 4.1 (2017), pp. 1–9.

[77] Markus Loeffler et al. "The LIFE-Adult-Study: objectives and design of a population-based cohort study with 10,000 deeply phenotyped adults in Germany". In: *BMC public health* 15 (2015), pp. 1–14.

[78] Michael V Lombardo, Emma Ashwin, Bonnie Auyeung, Bhismadev Chakrabarti, Meng-Chuan Lai, Kevin Taylor, Gerald Hackett, Edward T Bullmore, and Simon Baron-Cohen. "Fetal programming effects of testosterone on the reward system and behavioral approach tendencies in humans". In: *Biological psychiatry* 72.10 (2012), pp. 839–847.

[79] Antonios Makropoulos et al. "The developing human connectome project: A minimal processing pipeline for neonatal cortical surface reconstruction". In: *Neuroimage* 173 (2018), pp. 88–112.

[80] Kenneth Marek et al. "The Parkinson's progression markers initiative (PPMI)–establishing a PD biomarker cohort". In: *Annals of clinical and translational neurology* 5.12 (2018), pp. 1460–1477.

[81] Ross D Markello et al. "Neuromaps: structural and functional interpretation of brain maps". In: *Nature Methods* 19.11 (2022), pp. 1472–1479.

[82] John C Mazziotta, Arthur W Toga, Alan Evans, Peter Fox, Jack Lancaster, et al. "A probabilistic atlas of the human brain: theory and rationale for its development". In: *Neuroimage* 2.2 (1995), pp. 89–101.

[83] Karla L Miller et al. "Multimodal population brain imaging in the UK Biobank prospective epidemiological study". In: *Nature neuroscience* 19.11 (2016), pp. 1523–1536.

[84] José Luis Molinuevo et al. "The ALFA project: a research platform to identify early pathophysiological features of Alzheimer's disease". In: *Alzheimer's & Dementia: Translational Research & Clinical Interventions* 2.2 (2016), pp. 82–92.

[85] Sarah Morgan et al. *Data for "Cortical patterning of abnormal morphometric similarity in psychosis is associated with brain expression of schizophrenia-related genes"*. Apr. 2019. DOI: 10.6084/m9.figshare.7908488.v1. URL: https://figshare.com/articles/dataset/Data_for_Cortical_patterning_of_abnormal_morphometric_similarity_in_psychosis_is_associated_with_brain_expression_of_schizophrenia-related_genes_/7908488.

[86] Sarah Morgan et al. *Data for "Cortical patterning of abnormal morphometric similarity in psychosis is associated with brain expression of schizophrenia-related genes"*. Apr. 2019. DOI: 10.6084/m9.figshare.7908488.v1. URL: https://figshare.com/articles/dataset/Data_for_Cortical_patterning_of_abnormal_morphometric_similarity_in_psychosis_is_associated_with_brain_expression_of_schizophrenia-related_genes_/7908488.

[87] Sarah E Morgan et al. "Cortical patterning of abnormal morphometric similarity in psychosis is associated with brain expression of schizophrenia-related genes". In: *Proceedings of the National Academy of Sciences* 116.19 (2019), pp. 9604–9609.

[88] Hajer Nakua et al. "Cortico-amygdalar connectivity and externalizing/internalizing behavior in children with neurodevelopmental disorders". In: *Brain structure and function* 227.6 (2022), pp. 1963–1979.

[89] Ádám Nárai et al. *"Movement-related artefacts (MR-ART) dataset"*. OpenNeuro, 2022. DOI: doi:10.18112/openneuro.ds004173.v1.0.2.

[90] Ádám Nárai et al. "Movement-related artefacts (MR-ART) dataset of matched motion-corrupted and clean structural MRI brain scans". In: *Scientific data* 9.1 (2022), p. 630.

[91] Samuel A Nastase, Yaroslav O Halchenko, Andrew C Connolly, M Ida Gobbini, and James V Haxby. "Neural responses to naturalistic clips of behaving animals in two different task contexts". In: *Frontiers in neuroscience* 12 (2018), p. 316.

[92] Samuel A. Nastase et al. *"Narratives"*. OpenNeuro, 2025. DOI: 10.18112/openneuro.ds002345.v1.1.4.

[93] Dylan M. Nielson et al. *"National Institute of Mental Health Characterization and Treatment of Adolescent Depression (NIMH CAT-D)"*. OpenNeuro, 2023. DOI: doi:10.18112/openneuro.ds004627.v1.0.0.

[94] Aki Nikolaidis et al. "Heterogeneity in caregiving-related early adversity: Creating stable dimensions and subtypes". In: *Development and Psychopathology* 34.2 (2022), pp. 621–634.

[95] Allison C Nugent et al. "The NIMH intramural healthy volunteer dataset: A comprehensive MEG, MRI, and behavioral resource". In: *Scientific Data* 9.1 (2022), p. 518.

[96] Allison C. Nugent et al. *"The NIMH Healthy Research Volunteer Dataset"*. OpenNeuro, 2025. DOI: doi:10.18112/openneuro.ds005752.v2.1.0.

[97] N Charlotte Onland-Moret et al. "The YOUth study: Rationale, design, and study procedures". In: *Developmental cognitive neuroscience* 46 (2020), p. 100868.

[98] Deepak Pandya, Michael Petrides, and Patsy Benny Cipolloni. *Cerebral cortex: architecture, connections, and the dual origin concept*. Oxford University Press, 2015.

[99] Vishnu Parthasarathy et al. "Triglycerides are negatively correlated with cognitive function in nondemented aging adults." In: *Neuropsychology* 31.6 (2017), p. 682.

[100] Zdenka Pausova et al. "Genes, maternal smoking, and the offspring brain and body during adolescence: design of the Saguenay Youth Study". In: *Human brain mapping* 28.6 (2007), pp. 502–518.

[101] Russell A Poldrack et al. "A phenome-wide examination of neural and cognitive function". In: *Scientific data* 3.1 (2016), pp. 1–12.

[102] Jalil Rasgado-Toledo, Apurva Shah, Madhura Ingalhalikar, and Eduardo A Garza-Villarreal. "Neurite orientation dispersion and density imaging in cocaine use disorder". In: *Progress in Neuro-Psychopharmacology and Biological Psychiatry* 113 (2022), p. 110474.

[103] Siddharth Ray et al. "Structural and functional connectivity of the human brain in autism spectrum disorders and attention-deficit/hyperactivity disorder: A rich club-organization study". In: *Human brain mapping* 35.12 (2014), pp. 6032–6048.

[104] PK Reardon et al. "Normative brain size variation and brain shape diversity in humans". In: *Science* 360.6394 (2018), pp. 1222–1227.

[105] Jess E Reynolds, Xiangyu Long, Dmitrii Paniukov, Mercedes Bagshawe, and Catherine Lebel. "Calgary Preschool magnetic resonance imaging (MRI) dataset". In: *Data in brief* 29 (2020), p. 105224.

[106] Hilary Richardson, Grace Lisandrelli, Alexa Riobueno-Naylor, and Rebecca Saxe. *"MRI data of 3-12 year old children and adults during viewing of a short animated film"*. OpenNeuro, 2019.

[107] Hilary Richardson, Grace Lisandrelli, Alexa Riobueno-Naylor, and Rebecca Saxe. "Development of the social brain from age three to twelve years". In: *Nature communications* 9.1 (2018), p. 1027.

[108] Jonathan D Rohrer et al. "Presymptomatic cognitive and neuroanatomical changes in genetic frontotemporal dementia in the Genetic Frontotemporal dementia Initiative (GENFI) study: a cross-sectional analysis". In: *The Lancet Neurology* 14.3 (2015), pp. 253–262.

[109] Caitlin K Rollins et al. "Regional brain growth trajectories in fetuses with congenital heart disease". In: *Annals of neurology* 89.1 (2021), pp. 143–157.

[110] Chris Rorden and John Absher. *"Stroke Outcome Optimization Project (SOOP)"*. OpenNeuro, 2023. DOI: doi:10.18112/openneuro.ds004889.v1.0.0.

[111] Adon FG Rosen et al. "Quantitative assessment of structural image quality". In: *Neuroimage* 169 (2018), pp. 407–418.

[112] Therese Rydberg Sterner et al. "The Gothenburg H70 Birth cohort study 2014–16: design, methods and study population". In: *European journal of epidemiology* 34 (2019), pp. 191–209.

[113] Neda Sadeghi et al. "Mood and behaviors of adolescents with depression in a longitudinal study before and during the COVID-19 pandemic". In: *Journal of the American Academy of Child & Adolescent Psychiatry* 61.11 (2022), pp. 1341–1350.

[114] Giovanni Abrahão Salum et al. "High risk cohort study for psychiatric disorders in childhood: rationale, design, methods and preliminary results". In: *International journal of methods in psychiatric research* 24.1 (2015), pp. 58–73.

[115] Kazunori Sato, Yasuyuki Taki, Hiroshi Fukuda, and Ryuta Kawashima. "Neuroanatomical database of normal Japanese brains". In: *Neural networks* 16.9 (2003), pp. 1301–1310.

[116] Theodore D Satterthwaite et al. "Neuroimaging of the Philadelphia neurodevelopmental cohort". In: *Neuroimage* 86 (2014), pp. 544–553.

[117] Jenna M Schabdach et al. "Brain Growth Charts for Quantitative Analysis of Pediatric Clinical Brain MRI Scans with Limited Imaging Pathology". In: *Radiology* 309.1 (2023), e230096.

[118] Gunter Schumann et al. "The IMAGEN study: reinforcement-related behaviour in normal brain function and psychopathology". In: *Molecular psychiatry* 15.12 (2010), pp. 1128–1139.

[119] Roni Setton et al. "Age differences in the functional architecture of the human brain". In: *Cerebral Cortex* 33.1 (2023), pp. 114–134.

[120] Meredith A Shafto et al. "The Cambridge Centre for Ageing and Neuroscience (Cam-CAN) study protocol: a cross-sectional, lifespan, multidisciplinary examination of healthy cognitive ageing". In: *BMC neurology* 14 (2014), pp. 1–25.

[121] Mariana da Silva, Kaili Liang, Kara E Garcia, Jorge Cardoso, and Emma Claire Robinson. "Differential patterns of cortical expansion in fetal and preterm brain development". In: *bioRxiv* (2025), pp. 2025–05.

[122] Lukas Snoek, Maite van der Miesen, Andries van der Leij, Tinka Beemsterboer, Annemarie Eigenhuis, and Steven Scholte. *"AOMIC-ID1000"*. OpenNeuro, 2021. DOI: `10.18112/openneuro.ds003097.v1.2.1`.

[123] Lukas Snoek, Maite van der Miesen, Andries van der Leij, Tinka Beemsterboer, Annemarie Eigenhuis, and Steven Scholte. *"AOMIC-PIOP1"*. OpenNeuro, 2020. DOI: `10.18112/openneuro.ds002785.v2.0.0`.

[124] Lukas Snoek, Maite van der Miesen, Andries van der Leij, Tinka Beemsterboer, Annemarie Eigenhuis, and Steven Scholte. *"AOMIC-PIOP2"*. OpenNeuro, 2020. DOI: `10.18112/openneuro.ds002790.v2.0.0`.

[125] Lukas Snoek, Maite M van der Miesen, Tinka Beemsterboer, Andries Van Der Leij, Annemarie Eigenhuis, and H Steven Scholte. "The Amsterdam Open MRI Collection, a set of multimodal MRI datasets for individual difference analyses". In: *Scientific data* 8.1 (2021), p. 85.

[126] R. Nathan Spreng et al. *"Neurocognitive aging data release with behavioral, structural, and multi-echo functional MRI measures"*. OpenNeuro, 2022. DOI: `doi:10.18112/openneuro.ds003592.v1.0.9`.

[127] D Mikis Stasinopoulos and Robert A Rigby. "Generalized additive models for location scale and shape (GAMLSS) in R". In: *Journal of Statistical Software* 23 (2008), pp. 1–46.

[128] Lachlan T Strike, Narelle K Hansell, Baptiste Couvy-Duchesne, Paul M Thompson, Greig I de Zubicaray, Katie L McMahon, and Margaret J Wright. "Genetic complexity of cortical structure: differences in genetic and environmental factors influencing cortical surface area and thickness". In: *Cerebral cortex* 29.3 (2019), pp. 952–962.

[129] Valerie J Sydnor et al. "Neurodevelopment of the association cortices: Patterns, mechanisms, and implications for psychopathology". In: *Neuron* 109.18 (2021), pp. 2820–2846.

[130] Saori C Tanaka et al. "A multi-site, multi-disorder resting-state magnetic resonance image database". In: *Scientific data* 8.1 (2021), p. 227.

[131] Jason R Taylor, Nitin Williams, Rhodri Cusack, Tibor Auer, Meredith A Shafto, Marie Dixon, Lorraine K Tyler, Richard N Henson, et al. "The Cambridge Centre for Ageing and Neuroscience (Cam-CAN) data repository: Structural and functional MRI, MEG, and cognitive data from a cross-sectional adult lifespan sample". In: *neuroimage* 144 (2017), pp. 262–269.

[132] Loreen Tisdall and Rui Mata. *"AgeRisk"*. OpenNeuro, 2023. DOI: `doi:10.18112/openneuro.ds004711.v1.0.0`.

[133] Loreen Tisdall, Simon Mugume, David Kellen, and Rui Mata. "Lifespan trajectories of risk preference, impulsivity, and self-control: A dataset containing self-report, informant-report, behavioral, hormone and functional neuroimaging measures from a cross-sectional human sample". In: *Data in Brief* 52 (2024), p. 109968.

[134] Russell H Tobe et al. "A longitudinal resource for studying connectome development and its psychiatric associations during childhood". In: *Scientific data* 9.1 (2022), p. 300.

[135] Nicolas Traut et al. "Insights from an autism imaging biomarker challenge: promises and threats to biomarker discovery". In: *NeuroImage* 255 (2022), p. 119171.

[136] Jennifer Tremblay-Mercier et al. "Open science datasets from PREVENT-AD, a longitudinal cohort of pre-symptomatic Alzheimer's disease". In: *NeuroImage: Clinical* 31 (2021), p. 102733.

[137] Matilde M Vaghi et al. "Specific frontostriatal circuits for impaired cognitive flexibility and goal-directed planning in obsessive-compulsive disorder: evidence from resting-state functional connectivity". In: *Biological psychiatry* 81.8 (2017), pp. 708–717.

[138] Pedro A Valdes-Sosa et al. "The Cuban Human Brain Mapping Project, a young and middle age population-based EEG, MRI, and cognition dataset". In: *Scientific Data* 8.1 (2021), p. 45.

[139] Jennifer Vannest et al. "Factors determining success of awake and asleep magnetic resonance imaging scans in nonsedated children". In: *Neuropediatrics* 45.06 (2014), pp. 370–377.

[140] František Váša et al. "Rapid processing and quantitative evaluation of structural brain scans for adaptive multimodal imaging". In: *Human Brain Mapping* 43.5 (2022), pp. 1749–1765.

[141] František Váša et al. "Ultra-low-field brain MRI morphometry: test-retest reliability and correspondence to high-field MRI". In: *bioRxiv* (2024), pp. 2024–08.

[142] Lana Vasung, Caitlin K Rollins, Clemente Velasco-Annis, Hyuk Jin Yun, Jennings Zhang, Simon K Warfield, Henry A Feldman, Ali Gholipour, and P Ellen Grant. "Spatiotemporal differences in the regional cortical plate and subplate volume growth during fetal development". In: *Cerebral Cortex* 30.8 (2020), pp. 4438–4453.

[143] Lana Vasung et al. "Quantitative in vivo MRI assessment of structural asymmetries and sexual dimorphism of transient fetal compartments in the human brain". In: *Cerebral Cortex* 30.3 (2020), pp. 1752–1767.

[144] L. M. Villa et al. "Sex differences in brain development in fetuses and infants who are at low or high likelihood for autism". In: *medRxiv* (2021). Publisher: Cold Spring Harbor Laboratory Press _eprint: https://www.medrxiv.org/content/ DOI: 10.1101/2021.03.08.21251862. URL: https://www.medrxiv.org/content/early/2021/03/11/2021.03.08.21251862.

[145] Jin Wang, Marisa N Lytle, Yael Weiss, Brianna L Yamasaki, and James R Booth. "A longitudinal neuroimaging dataset on language processing in children ages 5, 7, and 9 years old". In: *Scientific Data* 9.1 (2022), p. 4.

[146] Jin Wang, Marisa N. Lytle, Yael Weiss, Brianna L. Yamasaki, and James R. Booth. *"A longitudinal neuroimaging dataset on language processing in children ages 5, 7, and 9 years old"*. OpenNeuro, 2021. DOI: 10.18112/openneuro.ds003604.v1.0.2.

[147] Cassandra MJ Wannan et al. "Accelerating Medicines Partnership® Schizophrenia (AMP® SCZ): rationale and study design of the largest global prospective cohort study of clinical high risk for psychosis". In: *Schizophrenia bulletin* 50.3 (2024), pp. 496–512.

[148] Varun Warrier et al. "Genetic insights into human cortical organization and development through genome-wide analyses of 2,347 neuroimaging phenotypes". In: *Nature genetics* 55.9 (2023), pp. 1483–1493.

[149] Dongtao Wei, Kaixiang Zhuang, Lei Ai, Qunlin Chen, Wenjing Yang, Wei Liu, Kangcheng Wang, Jiangzhou Sun, and Jiang Qiu. "Structural and functional brain scans from the cross-sectional Southwest University adult lifespan dataset". In: *Scientific data* 5.1 (2018), pp. 1–10.

[150] Margaret L Westwater, Flavia Mancini, Adam X Gorka, Jane Shapleske, Jaco Serfontein, Christian Grillon, Monique Ernst, Hisham Ziauddeen, and Paul C Fletcher. "Prefrontal responses during proactive and reactive inhibition are differentially impacted by stress in anorexia and bulimia nervosa". In: *Journal of Neuroscience* 41.20 (2021), pp. 4487–4499.

[151] Kirstie J Whitaker et al. "Adolescence is associated with genomically patterned consolidation of the hubs of the human brain connectome". In: *Proceedings of the National Academy of Sciences* 113.32 (2016), pp. 9105–9110.

[152] Ayumu Yamashita et al. "Harmonization of resting-state functional MRI data across multiple imaging sites via the separation of site differences into sampling bias and measurement bias". In: *PLoS biology* 17.4 (2019), e3000042.

[153] Michal Rafal Zareba, Magdalena Fafrowicz, Tadeusz Marek, Ewa Beldzik, Halszka Oginska, and Aleksandra Domagalik. *"Structural (t1) images of 136 young healthy adults; study of effects of chronotype, sleep quality and daytime sleepiness on brain structure."*. OpenNeuro, 2022. DOI: doi:10.18112/openneuro.ds003826.v3.0.1.

[154] Michal Rafal Zareba, Magdalena Fafrowicz, Tadeusz Marek, Ewa Beldzik, Halszka Oginska, and Aleksandra Domagalik. "Late chronotype is linked to greater cortical thickness in the left fusiform and entorhinal gyri". In: *Biological Rhythm Research* 53.10 (2022), pp. 1626–1638.

[155] Xi-Nian Zuo and CCNP Consortium. *"Chinese Color Nest Project (CCNP)"*. Version V2. Science Data Bank, 2023. DOI: 10.57760/sciencedb.07478.
